# *Tyr* is Responsible for the *Cctq1a* QTL and Links Developmental Environment to Central Corneal Thickness Determination

**DOI:** 10.1101/2021.09.22.461410

**Authors:** Kacie J. Meyer, Demelza R. Larson, S. Scott Whitmore, Carly J. van der Heide, Adam Hedberg-Buenz, Laura M. Dutca, Swanand Koli, Nicholas Pomernackas, Hannah E. Mercer, Maurisa N. Mansaray, William J. Paradee, Kai Wang, K. Saidas Nair, Todd E. Scheetz, Michael G. Anderson

**Affiliations:** Department of Molecular Physiology and Biophysics, University of Iowa, Iowa City, Iowa; Department of Ophthalmology and Visual Sciences, University of Iowa, Iowa City, Iowa; Center for the Prevention and Treatment of Visual Loss, Iowa City VA Health Care System, Iowa City, Iowa; Department of Ophthalmology, University of California, San Francisco, California; Genome Editing Core Facility, University of Iowa, Iowa City, Iowa; Department of Biostatistics, University of Iowa, Iowa City, Iowa; Department of Anatomy, University of California, San Francisco, California

## Abstract

Central corneal thickness is a quantitative trait with important associations to human health. In a phenotype-driven approach studying corneal thickness of congenic derivatives of C57BLKS/J and SJL/J mice, the critical region for a quantitative trait locus influencing corneal thickness, *Cctq1a*, was delimited to a 10-gene interval. Exome sequencing, RNAseq, and studying independent mutations eliminated multiple candidate genes and confirmed one. Though the causative gene, *Tyr*, has no obvious direct function in the transparent cornea, studies with multiple alleles on matched genetic backgrounds, both in isolation and genetic complementation crosses, confirmed allelism of *Tyr-Cctq1a*; albino mice lacking *Tyr* function had thin corneas. Albino mice also had increased axial length. Because albinism exposes eyes to increased light, the effect of dark-rearing was tested and found to rescue central corneal thickness. In sum, the results point to an epiphenomenon; developmental light exposure interacts with genotype as an important determinate of adult corneal thickness.

## Introduction

Central corneal thickness has important associations with ocular disease, but its natural determining factors remain predominately elusive. The cornea consists of three cellular layers (an outermost epithelium, middle stroma, and innermost endothelium) separated by two thinner basement membranes. The combined central thickness of these layers (central corneal thickness, CCT) increases rapidly through infancy and early childhood, reaches adult values in pre-teen ages, and remains relatively stable thereafter, with eventual modest age-related thinning^1-3, 4^. For largely unknown reasons, average CCT can vary by dozens of microns between ethnicities^4,5^. Thin CCT is associated with several corneal diseases, such as corneal dystrophy, brittle cornea syndrome, keratoconus, and cornea plana; diseases of connective tissue, such as Marfan syndrome, Ehlers-Danlos syndrome, Loeys-Dietz syndrome, and osteogenesis imperfecta; and at least two diseases in which the nature of, and/or reason for, the association is unclear, including myopia and primary open angle glaucoma^6^.

CCT is a highly heritable trait^7,8^, leading to many genetic studies. Variants influencing CCT have been identified from familial studies of Mendelian syndromes and GWAS of various large populations^9-12^. From these studies, some themes have begun to emerge. For example, some of the same CCT loci are identified in both multigenic and Mendelian disease studies^10,13,14^. It has also been common to identify variants related to collagen matrix integrity^10,12,14,15^. However, it is also clear that much remains unknown. Among the known associations, most known SNPs occur in non-coding regions, and the nearest genes typically have no obvious link to known structural components of the cornea^15^. It is also clear that many important genes remain to be discovered. Known SNPs give rise to a SNP-based heritability estimate of 42.5% and account for only 14.2% of the CCT variance^9^.

Here, an unique approach complementary to others is undertaken using inbred mouse strains to identify quantitative trait loci (QTL) that influence CCT^16,17^. Similar to humans, there is natural variation of CCT among inbred mouse strains^18^. Previous work using a quantitative approach with intercrosses between two such strains (C57BLKS/J [KS] mice with thin corneas and SJL/J [SJL] mice with thick corneas) identified the first CCT QTL, *Central corneal thickness QTL 1* (*Cctq1*) on mouse chromosome 7^16^. Here, *Cctq1* was resolved into two closely linked regions, *Cctq1a* and *Cctq1b*, which each influence CCT. Through multiple genetic approaches, a mutation in the tyrosinase gene (*Tyr*) is identified as causative of the *Cctq1a* phenotype, which appears to influence CCT via an epiphenomenon dependent on developmental light exposure.

## Results

### *Cctq1* contains two adjacent interacting QTL

The original 95% Bayesian credible interval of *Cctq1* spanned a 38.3 cM region on chromosome 7 (34.1 cM – 72.4 cM)^16^. To reduce this interval, recombination mapping was used with 92 recombinant N4 intercross mice. However, the interval was originally recalcitrant to division— more than one interval conferred the increased CCT phenotype. This suggested that there was too much genetic heterogeneity for this trait at the N4 generation, potentially including the presence of more than one CCT-regulating gene within or near *Cctq1*. To address these possibilities, the original F2 dataset was subjected to additional evaluations while backcrossing of the congenic mice to the N10 generation was continued.

The original analysis of (KS Χ SJL) F2 mice was based upon a significance threshold determined empirically by stratified permutation testing with 1000 permutations^16,19^, and did not identify any loci that significantly interacted with *Cctq1*. Prompted by the recombination mapping, the dataset was re-examined by performing a pairwise scan of the markers on chromosome 7 using scantwo analysis with R/qtl^19,20^. Of the possible interactions, the markers that produced the highest LOD scores were *D7Mit31* and *rs13479535* (Figure 1A; Figure 1 – Source Data 1; full LOD score = 10.33, interactive LOD score = 5.05). *D7Mit31* lies within the 95% Bayes credible interval of *Cctq1*; *rs13479535* is 2 cM distal to the end of the interval at 74.3 cM. The putative linked loci were subsequently subjected to multiple regression analysis in which each locus and the interaction component were sequentially dropped from the 2-QTL model. This analysis indicated that both loci and their interaction had significant contributions to the model (Figure 1 – Source Data 2). The QTL at *D7Mit31* was responsible for 31% of the phenotypic variability while the QTL at *rs13479535* accounted for another 24% of the variability. These data indicate that both loci are true QTL. *Cctq1* was thus resolved into two QTL, *Cctq1a* (95% Bayes credible interval: *D7Mit318*–*D7Mit220*, spanning 49.0 cM, peak at *D7Mit31*), and *Cctq1b* (95% Bayes credible interval: *D7Mit105*–*rs13479545*, spanning 74.3 cM, peak at *rs13479535*; Figure 1B).

**Figure 1.**
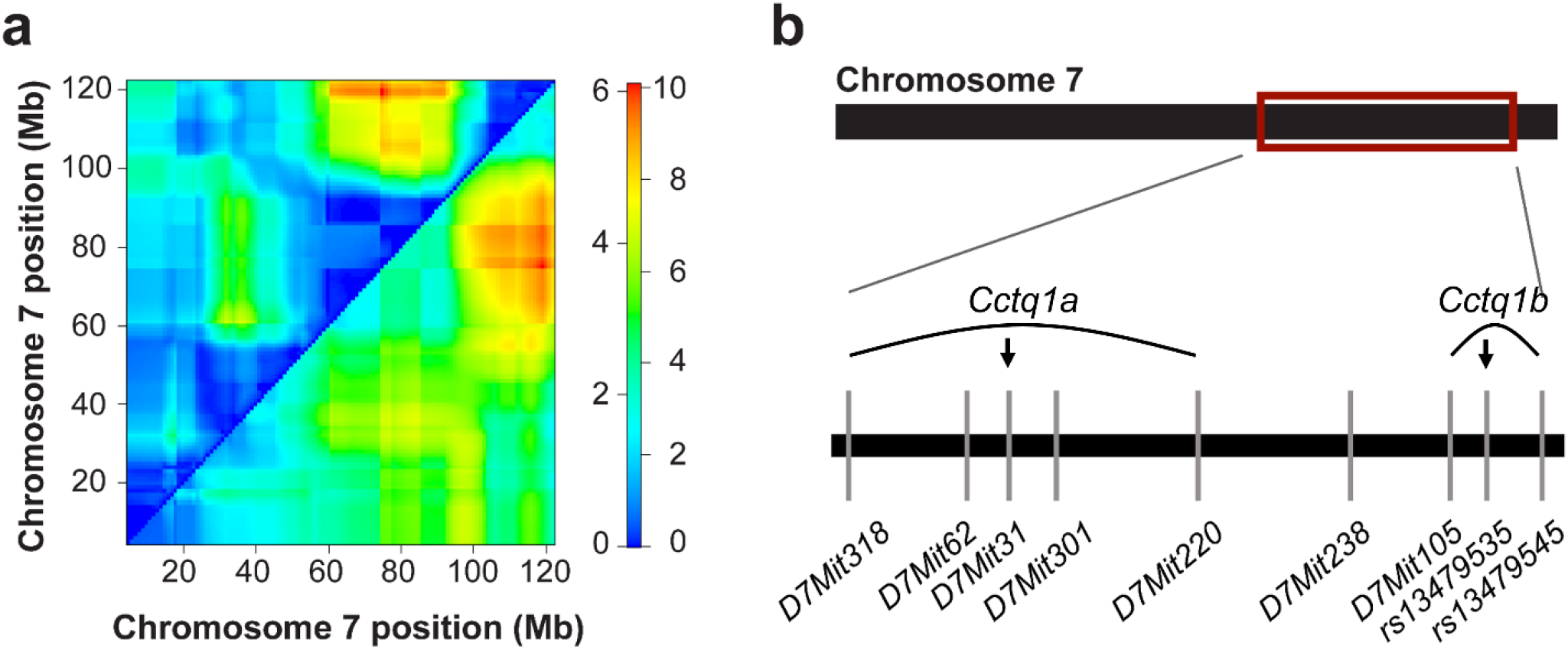
*Cctq1* contains two adjacent interacting QTL, *Cctq1a* and *Cctq1b*. **a** Heatmap of the chromosome 7 pairwise scan. The *upper left triangle* displays the interactive LOD score (LOD_*i*_; left side of the heat map scale) and the *lower right triangle* displays the full LOD score (LOD_*f*_; right side of the heat map scale). Chromosome 7 positions (Mb) are based on NCBI Build 33. **b** Genetic map of chromosome 7 showing the *Cctq1a* and *Cctq1b* loci. The area *boxed in red* is the original *Cctq1* interval blown up to show the intervals of *Cctq1a* and *Cctq1b*, and a subset of the polymorphic markers used for genotyping. The *arrows* indicate the peaks of the two QTL.

To reduce genetic heterogeneity, N4 mice were further backcrossed onto the KS background to the N10 generation. Because the congenic interval was relatively large, a panel of six markers was used at each generation of backcrossing to keep the interval intact. At generation N10, congenic mice were intercrossed, recombination within the interval was allowed, and mice with all nine genetic combinations of *Cctq1a* and *Cctq1b* were phenotyped for CCT (Table 1; Table 1 – Source Data). Congenic control mice (KS.SJL-*Cctq1a*^*KS*^,*Cctq1b*^*KS*^) had a CCT indistinguishable from inbred KS mice (94.8± 2.4 μm vs. 94.5 ± 3.2 μm, respectively; one-way ANOVA with Tukey post-test; Table 1). Congenic mice with SJL genotypes at *Cctq1a* (*i*.*e*., KS.SJL-*Cctq1a*^*SJL*^) had significantly thinner corneas than inbred KS mice (87.9 ± 3.8 μm; *n* = 39; *p* < 0.001; Student’s two-tailed *t*-test) independent of the genotype at *Cctq1b* (*n* = 13 and *p* < 0.05 for each of three genotypes at *Cctq1b*; one-way ANOVA with Tukey post-test; Table 1; Supplementary Figure 1). The difference in thickness mediated by *Cctq1a* is predominantly due to the thickness of the stroma (Δ 9.6 μm, *p* < 0.001; Student’s two-tailed *t*-test), though there is a small (Δ 1.2 μm) and marginally significant (*p* = 0.022 Student’s two-tailed *t*-test) decrease in thickness of the epithelium as well. Congenic mice with KS genotypes at *Cctq1a* and SJL genotypes at *Cctq1b* (i.e., KS.SJL-*Cctq1a*^*KS*^,*Cctq1b*^*SJL*^) also had significantly thinner corneas (90.8 ± 2.6 μm; *n* = 13 mice; *p* < 0.05; one way ANOVA with Tukey post-test) than inbred KS mice (Table 1, Supplementary Figure 1). No other genetic combinations caused significant changes in CCT compared to inbred KS controls. As predicted from the analysis of the original F2 Dataset with R/qtl, these data with congenic mice independently support that both *Cctq1a* and *Cctq1b* are true QTL capable of altering the phenotypic variability of CCT on a uniform genetic background.

**Table 1.**
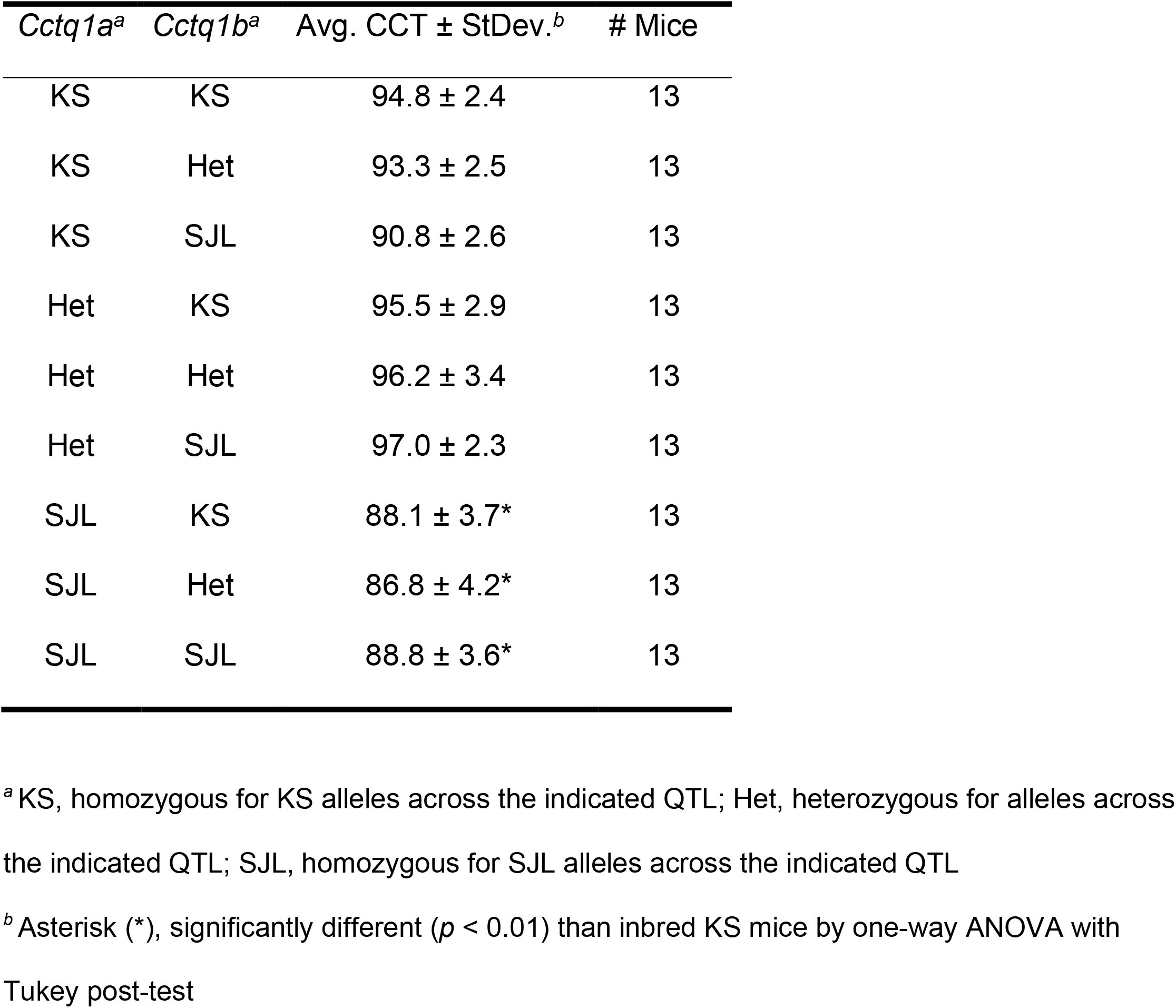
CCT phenotype results of adult N10F2 mice.

### Mapping of *Cctq1a* using recombination mapping and sub-congenics

Because of its larger effect, initial efforts were focused on fine-scale mapping for *Cctq1a*. To identify the gene underlying *Cctq1a, Cctq1a*-recombinant N10F2 mice, with KS genotypes at *Cctq1b*, were used to narrow the critical region. From this recombination analysis, the gene underlying *Cctq1a* was deduced to be between SNP markers *rs108403472* at 48.51 cM and *rs6247100* at 50.26 cM. Simultaneously, sub-congenic mice were created by continued backcrossing of the KS.SJL-*Cctq1a*^*HET*^*;Cctq1b*^*KS*^ N10 mice. Eyes of N10 congenic mice were overtly healthy, differing only in pigmentation between genotypes (Supplementary Figure 1). At N12, *Cctq1a* was physically reduced to a 15.9 cM region (KS.SJL-*Cctq1a(15*.*9cM)*) flanked by *D7mit347* and *D7mit321* and characterized by an association of SJL homozygosity with decreased CCT (87.6 ± 2.3 μm vs. 76.8 ± 2.4 μm, *n* = 15 mice per genotype, *p* < 0.001, Student’s two-tailed *t*-test). At N15, *Cctq1a* was physically reduced to a 9.9 cM region (KS.SJL-*Cctq1a(9*.*9cM)*) flanked by *rs3672782* and *D7mit321*; again characterized by an association of SJL homozygosity with decreased CCT (91.6 ± 2.2 μm vs. 80.7 ± 2.9 μm, *n* = 7 mice per genotype, *p* < 0.001, Student’s two-tailed *t*-test). One N15F2 mouse harbored a recombination event within the minimal sub-congenic interval. The phenotype of this mouse indicated the gene underlying *Cctq1a* lies proximal to marker *rs13479393* (Figure 2; Figure 2 – Source Data). Using this recombinant mouse in a progeny test, additional intercrossing to generation N15F7 confirmed the distal breakpoint proximal to marker *rs13479393* at 49.65 cM, again characterized by an association of SJL homozygosity with decreased CCT (85.7 ± 1.6 μm vs. 94.4 ± 3.3 μm, *n* = 8–10 mice per genotype; *p* < 0.001, Student’s two-tailed *t*-test; Figure 3A; Figure 3 – Source data 1−3).

**Figure 2.**
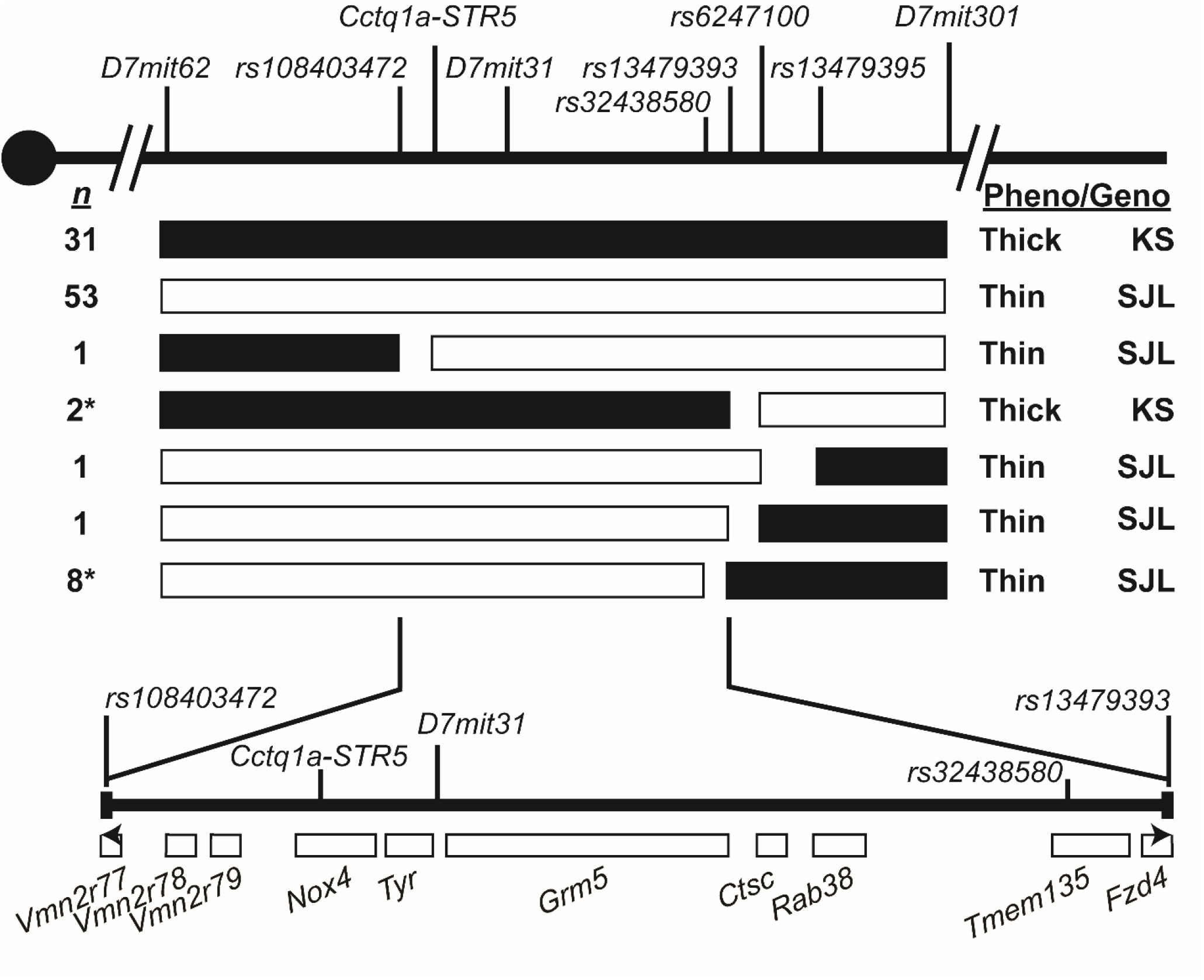
Genetic mapping of *Cctq1a* on mouse chromosome 7 using intercrosses of KS.SJL-*Cctq1a* congenic and sub-congenic mice. *Black boxes* represent the KS or HET genotype associated with a thick cornea and *white boxes* represent the SJL genotype associated with a thin cornea. The number of mice (*n*) with each haplotype is listed to the left of each row, with progeny-tested mice denoted with an asterisk (*). The adult CCT phenotype (*pheno*; measured by optical coherence tomography) relative to littermate controls and the deduced genotype (*geno*) for each haplotype is listed to the right of each row as “Thick KS” or “Thin SJL”. The *vertical lines* across the chromosome represent markers that are polymorphic between KS and SJL mice. *Black arrowheads* indicate partial genes.

**Figure 3.**
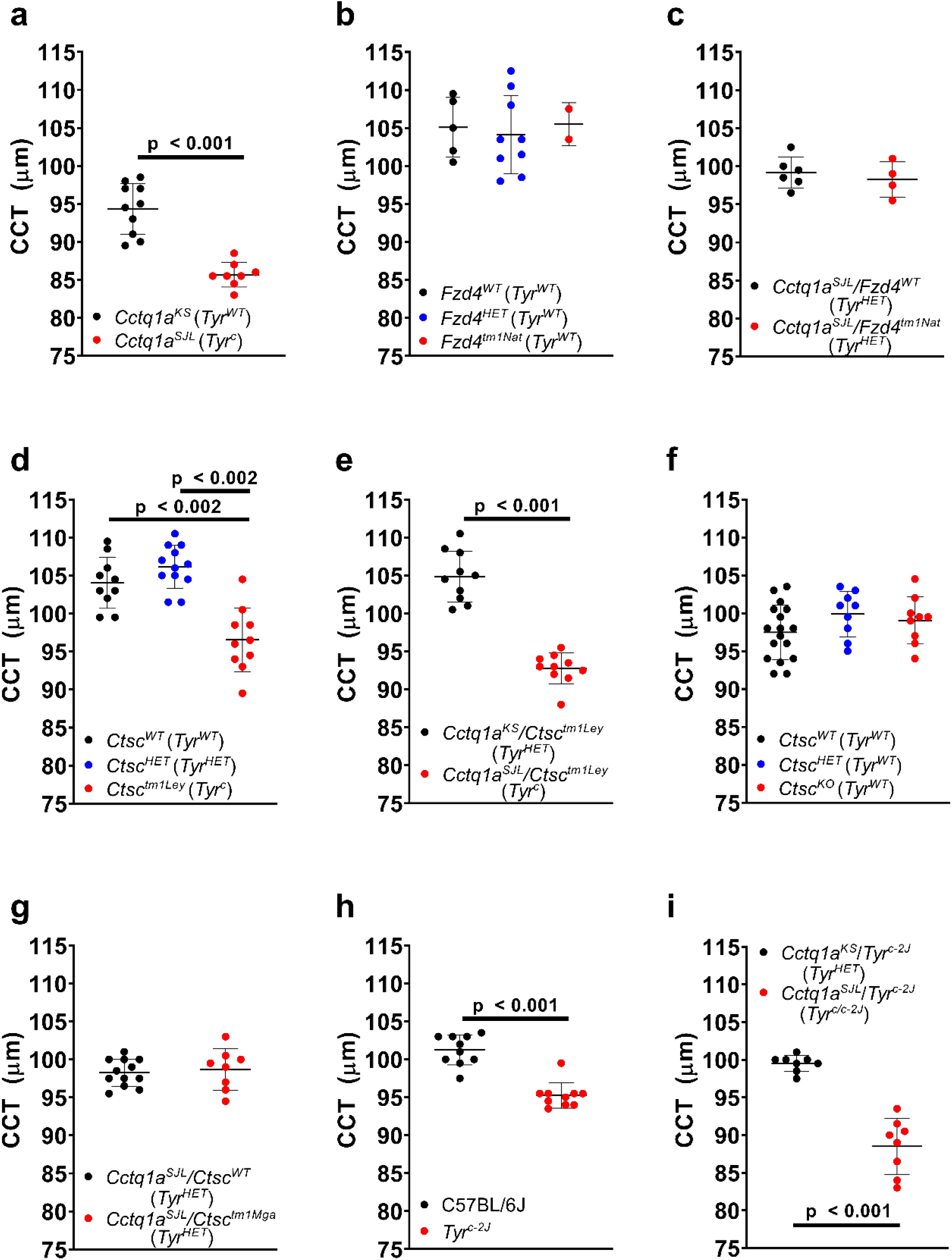
Testing the influence of *Cctq1a* positional candidate genes on central corneal thickness (CCT). Each point on the graph represents the average CCT measured by optical coherence tomography from one adult mouse with age-matching across genotypes and *error bars* = mean ± standard deviation. **a** Plot comparing CCT for *Cctq1a*^*KS*^ vs. *Cctq1a*^*SJL*^ genotypes in KS.SJL-*Cctq1a* N15F7 sub-congenic littermate mice 12 to13 weeks old (Student’s two-tailed *t*-test). **b** Plot comparing CCT between *Fzd4* genotypes for littermate mice 12 to 13 weeks old (one-way ANOVA with Tukey test). **c** Plot comparing the CCT for mice with a *Fzd4*^*tm1Nat*^ allele in trans with a *Cctq1a*^*SJL*^ allele to littermate controls having a *Fzd4*^*WT*^ allele in trans with a *Cctq1a*^*SJL*^ allele (Student’s two-tailed *t*-test; 17 to 33 weeks old). **d** Plot comparing CCT between *Ctsc*^*tm1Ley*^ genotypes for 11-week-old littermate mice (one-way ANOVA with Tukey test). **e** Plot comparing CCT for mice with a *Ctsc*^*tm1Ley*^ allele in trans with a *Cctq1a*^*SJL*^ allele compared to littermate controls having a *Ctsc*^*tm1Ley*^ allele in trans with a *Cctq1a*^*KS*^ allele (Student’s two-tailed *t*-test; 15 to18 weeks old). **f** Plot comparing CCT for mice harboring one of four *Ctsc*^*KO*^ alleles on a pure B6 background to littermate controls (one-way ANOVA with Tukey test; 10 to 12 weeks old). **g** Plot comparing CCT for mice with a *Ctsc*^*tm1Mga*^ allele, made on a pure B6 background, in trans with a *Cctq1a*^*SJL*^ allele to littermate controls having a *Ctsc*^*WT*^ allele in trans with a *Cctq1a*^*SJL*^ allele (Student’s two-tailed *t*-test; 10 to 11 weeks old). **h** Plot comparing CCT for B6.*Tyr*^*c-2J*^ mice to B6 mice (Student’s two-tailed *t*-test; 11 to 13 weeks old). **i** Plot comparing CCT for mice with a *Tyr*^*c-2J*^ allele in trans with a *Cctq1a*^*SJL*^ allele to littermate controls having a *Tyr*^*c-2J*^ allele in trans with a *Cctq1a*^*KS*^ allele (Student’s two-tailed *t*-test; 10 to 12 weeks old).

In sum, physical recombination mapping utilizing multiple generations of congenic and sub-congenic mice conclusively indicated that the gene underlying *Cctq1a* lies on chromosome 7 between markers *rs108403472* at 48.51 cM and *rs13479393* at 49.32 cM, a 0.81 cM region containing the entirety of eight RefSeq genes (*Vmn2r78, Vmn2r79, Nox4, Tyr, Grm5, Ctsc, Rab38*, and *Tmem135*), the 3’ portion of one gene (*Vmn2r77*), and the 5’ portion of one gene (*Fzd4*) (Figure 2).

### Candidate identification and prioritization

To identify all the possible exonic variants within the *Cctq1a* critical region, whole exome sequencing was conducted on KS and SJL inbred mice. In the entire exome, 15,261 missense, frameshift, and splice-site mutations were found between KS and SJL (Supplementary File 1). There were six amino acid altering variants within the *Cctq1a* critical region, located within *Vmn2r79* (A223T, L243M, T257I, I265V), *Tyr* (C103S), and *Fzd4* (F27L). Vomeronasal receptor genes, such as *Vmn2r79*, have an increased rate of coding sequence variants ^21^ and the four altered residues between KS and SJL in *Vmn2r79* are poorly conserved. The amino acid change in *Tyr* from cysteine to serine is the albinism-causing *Tyr*^*c*^ allele conferred by the albino SJL strain. The *Fzd4* amino acid variant is within the signal sequence of the protein; KS mice conferred the phenylalanine amino acid residue (the same residue as C57BL/6J mice) while SJL mice conferred the mammalian-conserved leucine residue.

Transcriptional profiling was additionally used to prioritize candidates. Using previously published microarray data comparing adult corneal RNA expression profiles in KS and SJL mice^18^, *Nox4, Ctsc, Rab38, Tmem135*, and *Fzd4* were all present in the adult cornea; there was no evidence for adult corneal expression of *Vmn2r77, Vmn2r78, Vmn2r79, Tyr*, or *Grm5*. Of those expressed, *Nox4, Ctsc, and Fzd4* showed differential expression between the two strains. *Nox4* and *Ctsc* were both down-regulated 2.5-fold in SJL, while *Fzd4* was up-regulated 1.6-fold in SJL.

Additionally, RNAseq was performed on corneas of 3-week-old KS.SJL-*Cctq1a(15*.*9cM)*^*SJL*^ N12F3 mice. Analysis was focused on three comparisons: 1) KS.SJL-*Cctq1a(15*.*9cM)*^*SJL*^ vs. KS (experimental, identifying genes with altered corneal expression in the congenic interval), 2) KS.SJL-*Cctq1a(15*.*9cM)*^*SJL*^ vs. KS.SJL-*Cctq1a(15*.*9cM)*^*KS*^ (experimental, also identifying genes with altered corneal expression in the congenic interval), and 3) KS.SJL-*Cctq1a(15*.*9cM)*^*KS*^ vs. KS (control, identifying genes in the background of the congenic strain with altered corneal expression not associated with the congenic interval) (Supplementary File 2). In each comparison, genes were first filtered for those with a Q-value ≤ 0.001 and a FPKM ≥ 1 in at least one of the strains. Gene lists were subsequently compared to one another, identifying 87 genes consistently altered in both experimental comparisons but not in the control comparison (Supplementary File 2). Among these 87, only one was localized to the *Cctq1a* critical region, *Ctsc*, which was modestly (−0.8 log_2_fold) but consistently and significantly (*p* = 5 × 10^−5^; *Q* = 0.0009) down regulated in comparing the SJL allele to the KS allele. Web Gestalt^22^ was used for over-representation analysis comparing the list of 87 differentially expressed genes to a background list of all genes expressed in the cornea with a FPKM ≥ 1 in any strain. Results of the analysis indicate a strong signal for several collagen-related categories (fibrillar collagen trimer, abnormal cutaneous collagen fibril morphology, collagen biosynthesis and modifying enzymes, collagen degradation, etc.), extracellular-matrix-related categories (extracellular matrix component, ECM proteoglycans, degradation of the extracellular matrix, etc.) and ocular-related categories (decreased corneal stroma thickness, abnormal corneal epithelium morphology, abnormal cornea morphology, and abnormal eye morphology) (Supplementary Figure 2).

### Functional tests of lead candidates

Based on candidate prioritization criteria, *Fzd4* and *Ctsc* were initially considered the top candidates. To test the influence of *Fzd4* on CCT, we tested a strain with a targeted mutation of *Fzd4* (B6;129-*Fzd4*^*tm1Nat*^/J) on a segregating B6 and 129 background^23^. *Fzd4*^*tm1Nat*^ homozygotes had a chocolate coat color not expected from either background. This observation is meaningful as the gene responsible for this phenotype, *Rab38*, is physically near *Fzd4* and within the *Cctq1* critical region, i.e., the strain is likely a double mutant for *Fzd4* (genotype verified) and *Rab38* (mutation unknown). However, there was no correlation between *Fzd4* ^*tm1Nat*^ genotype and CCT (*p* = 0.819; one-way ANOVA comparing all three genotypes among littermates; Figure 3B; Supplementary Data 3–5). In genetic complementation crosses, *Fzd4* ^*tm1Nat*^ mutation complemented the congenic interval ([*Fzd4*^*WT*^/*Cctq1a*^*SJL*^ F1] vs. [*Fzd4*^*HET*^/*Cctq1a*^*SJL*^ F1]; *p* = 0.528; Student’s two-tailed *t*-test; *n* = 4-6 per genotype; Figure 3C; Figure 3 – Source Data 1−3). The complementation cross also highlighted the coat color phenotype associated with the *Fzd4*^*tm1Nat*^ mutation, with *Fzd4*^*WT*^/*Cctq1a*^*SJL*^ F1 mice (*Tyr*^*HET*^) having a black coat color with normally pigmented eyes and *Fzd4*^*HET*^/*Cctq1a*^*SJL*^ F1 mice (*Tyr*^*HET*^) having an unmistakable lightened (“light chocolate”) coat color and light brown irides instead of brown (Figure 3 – Source Data 5). Therefore, *Fzd4* was ruled out as causative of the *Cctq1a* phenotype, and *Rab38* further deprioritized as a candidate.

To test the influence of *Ctsc* on CCT, we imported a strain with a targeted mutation of *Ctsc* (B6.Cg-*Ctsc*^*tm1Ley*^) on an N10 congenic B6 background^24^. Because the targeted mutation was generated on an albino 129 background, and *Ctsc* is physically linked to *Tyr, Ctsc*^*tm1Ley*^ homozygotes are albino, i.e., the strain is a double mutant for *Ctsc* and *Tyr*. This is meaningful because the SJL/J strain is albino and the *Tyr*^*c*^ mutation is within the KS.SJL-*Cctq1a* congenic interval. Homozygotes had a statistically significant decrease in CCT compared to littermate controls (*p* < 0.002 for albino *Ctsc*^*tm1Ley*^ vs. pigmented *Ctsc*^*HET*^; *p* < 0.002 for albino *Ctsc*^*tm1Ley*^ vs. pigmented *Ctsc*^*WT*^; one-way ANOVA with Tukey post-test comparing all three genotypes among littermates; *n*=10–12 mice per genotype; Figure 3D; Figure 3 – Source Data 1−3). In genetic complementation crosses, *Ctsc*^*tm1Ley*^ failed to complement *Cctq1a*^*SJL*^ (*p* < 0.001 for pigmented [*Ctsc*^*tm1Ley*^/*Cctq1a*^*KS*^ F1] vs. albino [*Ctsc*^*tm1Ley*^/*Cctq1a*^*SJL*^ F1]; Student’s two-tailed *t*-test; *n* = 10– 11 mice per genotype; Figure 3E; Figure 3 – Source Data 1−3). Therefore, *Ctsc or Tyr* were determined to be causative of the *Cctq1a* phenotype.

To differentiate *Ctsc* and *Tyr* as the causative mutation, independent alleles on a pure B6 background were assessed. To test *Ctsc*, four new mutations predicted to result in null protein were generated in B6 mice with CRISPR-Cas9 technology (Figure 3 – Source Data 4). To test *Tyr*, a well-known spontaneous mutation that was commercially available, *Tyr*^*c-2J*^, was analyzed^25,26^. There was no association between *Ctsc*^*KO*^ genotype and CCT (*p* = 0.237; one-way ANOVA comparing all three genotypes among littermates; *n* = 9–17 mice per group; Figure 3F; Figure 3 – Source Data 1−4). In genetic complementation crosses with the congenic strain, the *Ctsc*^*tm1Mga*^ mutation (46bp-deletion in the coding sequence of exon 1 leading to no detectable CTSC protein; Figure 3 – Source Data 4) complemented the congenic phenotype (*p* = 0.696; Student’s two-tailed *t*-test; *n* = 8–12 mice per group; Figure 3G; Figure 3 – Source Data 1−4). In contrast, albino *Tyr*^*c-2J*^ mice had decreased CCT relative to pigmented B6 (*p* < 0.001; Student’s two-tailed *t*-test; *n* = 9–10 mice per group; Figure 3H; Figure 3 – Source Data 1–3). In genetic complementation crosses with the congenic strain, the *Tyr*^*c-2J*^ mutation failed to complement the congenic phenotype (*p* < 0.001 for pigmented [*Cctq1a*^*KS*^/*Tyr*^*c-2J*^ F1] vs. albino [*Cctq1a*^*SJL*^/*Tyr*^*c-2J*^ F1]; Student’s two-tailed *t*-test; *n* = 8 mice per genotype; Figure 3I; Figure 3 Source Data 1–3). Thus, only *Tyr* mutation was left as a feasible candidate for *Cctq1a*.

CRISPR-Cas9 technology was also used to generate new *Tyr* mutations on a C57BL/6J background (Figure 4 – Source Data 1). One allele was selected for propagation, *Tyr*^*tm4Mga*^, an albinism-causing 4 bp-deletion in the coding sequence of exon 1 that is predicted to cause a frameshift leading to a premature stop codon and RNA-mediated decay; i.e., a presumed null mutation. *Tyr*^*tm4Mga*^ mice had a significantly thinner cornea than littermate controls (*p* < 0.002 for *Tyr*^*WT*^ vs. *Tyr*^*tm4Mga*^; *p* < 0.002 for *Tyr*^*HET*^ vs. *Tyr*^*tm4Mga*^; one-way ANOVA with Tukey post-test comparing all three genotypes among littermates; *n* = 6–10 mice per genotype; Figure 4; Figure 4 – Source Data 2−3). Thus, a presumed null allele of *Tyr* caused the same albinism and thinning of CCT as found for the *c* and *c-2J* alleles.

**Figure 4.**
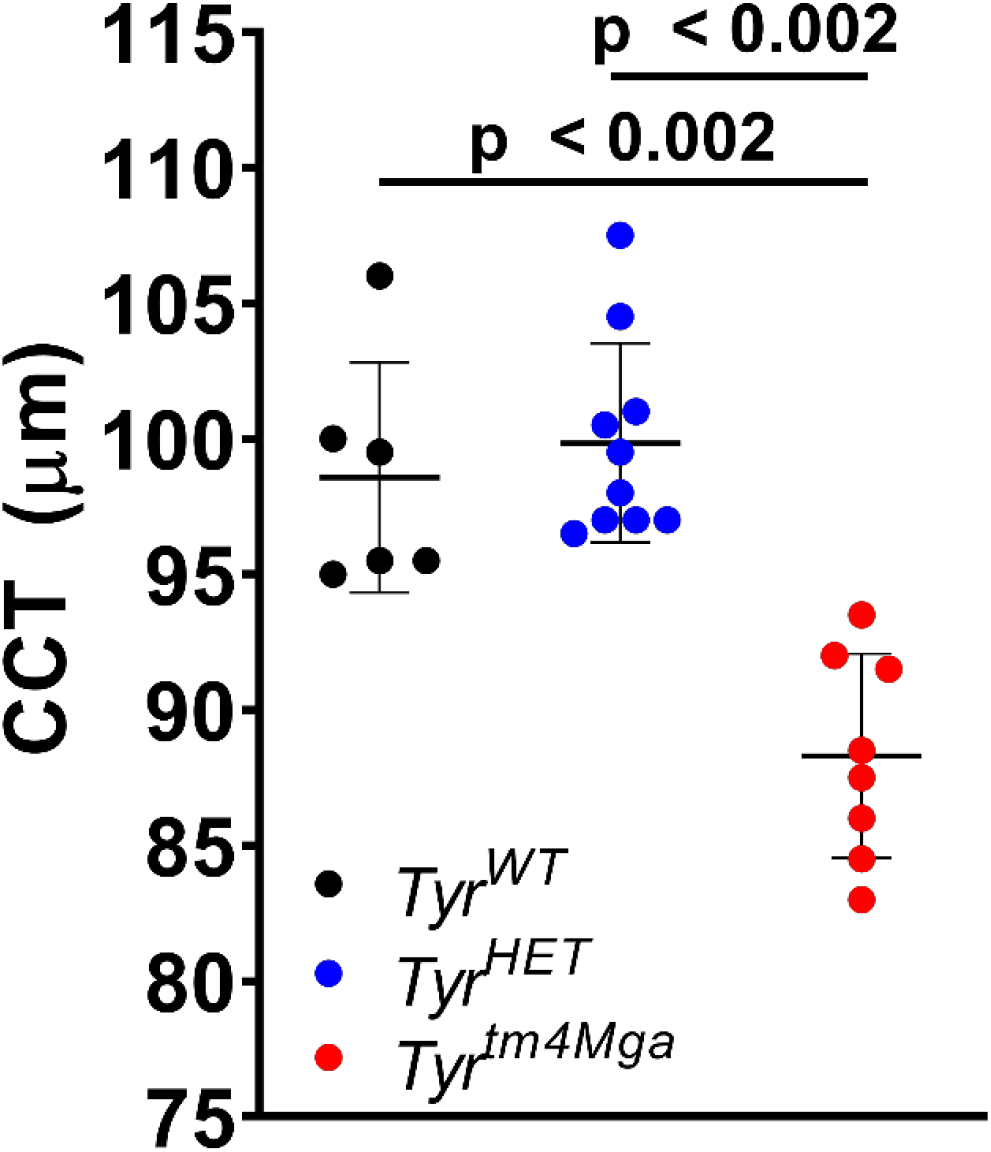
The influence of a null *Tyr* mutation on central corneal thickness (CCT). Each point on the graph represents the average CCT measured by optical coherence tomography from one adult mouse, 13 to 17 weeks old, with age-matching across genotypes and *error bars* = mean ± standard deviation. The plot compares CCT between mice homozygous for the *Tyr*^*tm4Mga*^ allele on a pure B6 background to littermate controls with heterozygous (HET) or wild-type (WT) alleles (one-way ANOVA with Tukey test).

### Mechanism of *Tyr* function on CCT

In considering the possible mechanism through which *Tyr* might influence CCT, three hypotheses were tested:

The first candidate mechanism centered on the role of DOPA, which is a cofactor in the oxidation of tyrosine by TYR, leading to melanin production^27^, and a substrate for tyrosine hydroxylase (TH), leading to dopamine synthesis. Dopamine is considered a key molecule in ocular growth^28,29^, and mice with a conditional knock-out of *Th* in the retina have previously been shown to have decreased CCT^30^. Rationalizing that some DOPA may normally escape from pigment producing cells to influence CCT via a TH-dependent mechanism, we tested whether providing supplemental DOPA in the drinking water of albino mice would rescue the decreased CCT associated with albinism. No statistically significant effect was observed (Figure 5; Figure 5 – Source Data 1−2), discounting the hypothesis that a DOPA deficit was rate-limiting for CCT determination in *Tyr* mutant mice.

**Figure 5.**
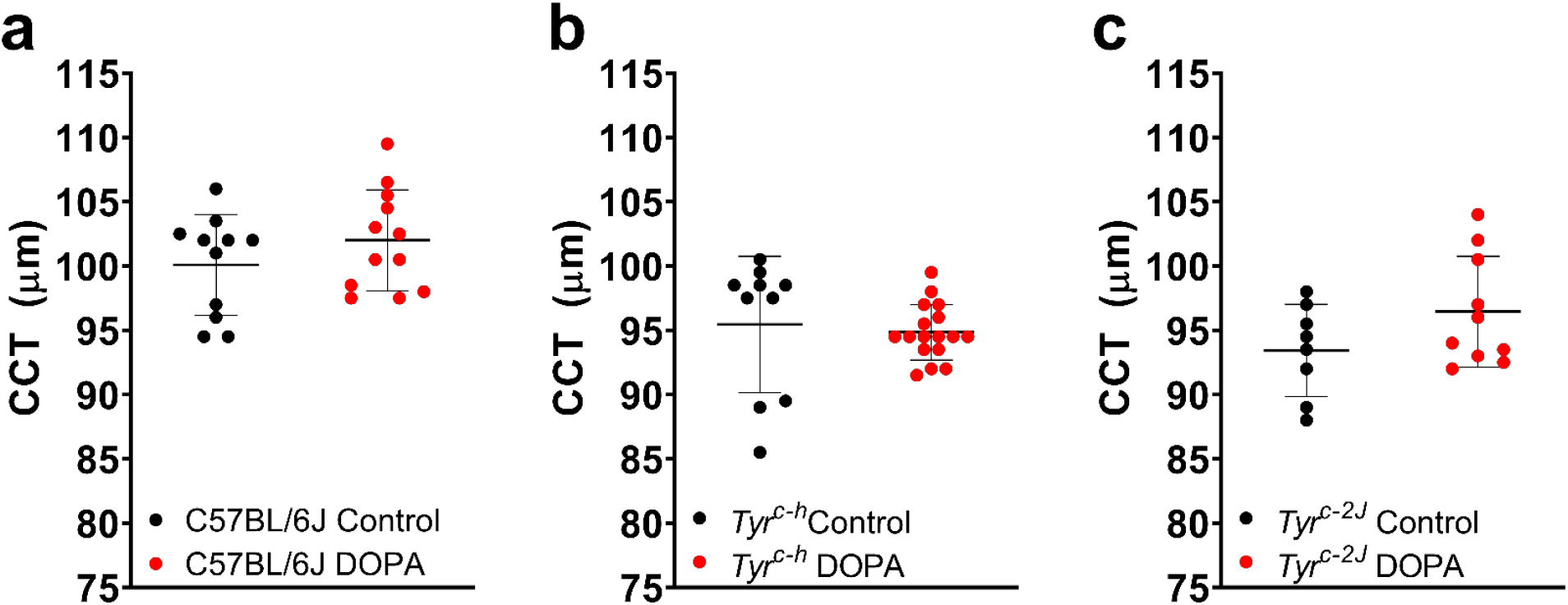
Testing the influence of DOPA on central corneal thickness (CCT). Each point on the graph represents the average CCT measured by optical coherence tomography from one adult mouse, 13 to 16 weeks old, with age-matching across genotypes and *error bars* = mean ± standard deviation. Plots compare CCT between mice supplied with DOPA in the drinking water from conception through 10-weeks-old to controls supplied with standard drinking water for the **a** B6, **b** B6.*Tyr*^*c-h*^, and **c** B6.*Tyr*^*c-2J*^ strains of mice (Student’s two-tailed *t*-test for each comparison).

The second candidate mechanism revolves around the possibility of a gene-environment interaction involving temperature and light. In this series of experiments, mice were reared in environmental control chambers in combinations of different temperatures and light cycles. Cohorts included pigmented B6, albino *Tyr*^*c-2J*^, and the temperature sensitive himalayan mutation (B6.Cg-*Tyr*^*c-h*^/J)^31,32^. At ambient temperatures, *Tyr*^*c-h*^ homozygotes are only partially pigmented on the coolest parts of the body (such as the ears, nose, tail, and eyes) and CCT is intermediate between B6 and *Tyr*^*c-2J*^ mice (Figure 6 – Source Data). For CCT of mice raised at ambient temperature, genetic complementation tests again confirmed the influence of *Tyr*-mediated albinism on CCT (*p* = 0.337 for [pigmented B6 x *Tyr*^*c-h*^ F1] vs. [pigmented B6 x *Tyr*^*c-2J*^ F1]; *p* < 0.002 for [pigmented B6 x *Tyr*^*c-h*^ F1] vs. [albino *Tyr*^*c-h*^ x *Tyr*^*c-2J*^ F1]; and *p* < 0.002 for [pigmented B6 x *Tyr*^*c-2J*^ F1] vs. [albino *Tyr*^*c-h*^ x *Tyr*^*c-2J*^ F1]; one-way ANOVA with Tukey post-test; *n* = 4–14 mice per group, (Figure 6 – Source Data)). At decreased temperatures *Tyr*^*c-h*^ homozygotes can generate pigment more broadly, and at increased temperatures the albinism is accentuated (Figure 6).

**Figure 6.**
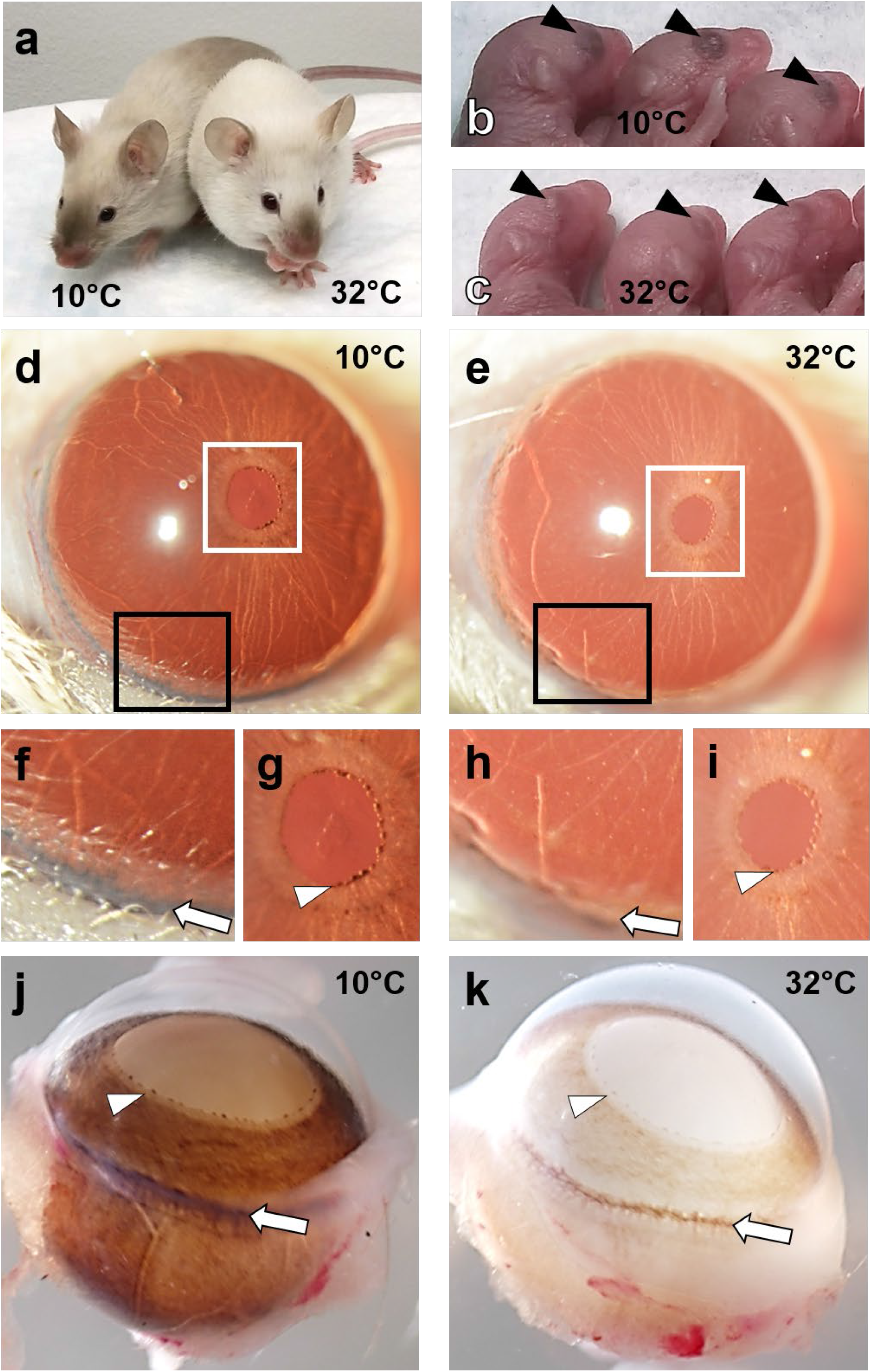
The influence of temperature on ocular pigment for the B6.*Tyr*^*c-h*^ strain of mice. **a** When reared at decreased temperatures (10°C; *left*) *Tyr*^*c-h*^ homozygotes can generate pigment more broadly and when reared at increased temperatures (32°C; *right*) the albinism is accentuated. There are also readily apparent differences, indicated by black arrowheads, in ocular pigment of *Tyr*^*c-h*^ homozygote neonates born at **b** decreased temperatures (10°C) compared to **c** increased temperatures (32°C). Slit-lamp transillumination images photographed with identical setting show that *Tyr*^*c-h*^ homozygote eyes have **d** increased pigment in mice reared at decreased temperatures (10°C) compared to **e** mice raised at increased temperatures (32°C). Magnified views of the black and white boxes in panel **d** are shown in **f** and **g**, respectively. Magnified views of the black and white boxes in panel **e** are shown in **i** and **j**, respectively. There is increased pigment in the limbal region (*white arrow*) of *Tyr*^*c-h*^ homozygotes **f** reared at decreased temperatures (10°C) compared to those **h** reared at increased temperatures (32°C). There is increased pigment in the pupillary rough (white arrowhead) of *Tyr*^*c-h*^ homozygotes **g** reared at decreased temperatures (10°C) compared to those **i** reared at increased temperatures (32°C). There is increased pigment in enucleated eyes from mice *Tyr*^*c-h*^ homozygotes **j** reared at decreased temperatures (10°C) compared to those **k** reared at increased temperatures (32°C). The pupillary rough is indicated by the white arrowhead and the limbus is indicated by the white arrow.

Regarding temperature, comparison of B6.*Tyr*^*c-h*^ mice, as well as B6 and B6.*Tyr*^*c-2J*^ controls, raised at 10°C vs. 32°C in environmental control chambers with a standard light cycle, showed that CCT followed pigment status; the thin CCT of hypopigmented *Tyr*^*c-h*^ mice reared at increased temperature (i.e., lower TYR activity) was rescued by rearing the *Tyr*^*c-h*^ mice at decreased temperature (i.e., higher TYR activity; *Tyr*^*c-h*^ at 32°C vs. *Tyr*^*c-h*^ at 10°C; Δ 10.9 μm; *p* < 0.001; *n* = 13–15 mice per condition; one-way ANOVA with Sidak test; Figure 7; Figure 7 – Source Data 1−2). In testing the effect of temperature on corneal thickness in controls, there was a small (Δ 2.8 μm) and nominally significant (*p* = 0.03) decreased CCT in mice raised at 10ºC compared to mice raised at 32ºC when considering all C57BL/6J and *Tyr*^*c-2J*^ mice across the experiment (*n* = 16 mice vs. *n* = 19 mice, Type II ANOVA) and there was no interaction between genotype and temperature. In sum, changing environmental temperature changed CCT of the *Tyr*^*c-h*^ mice as predicted.

**Figure 7.**
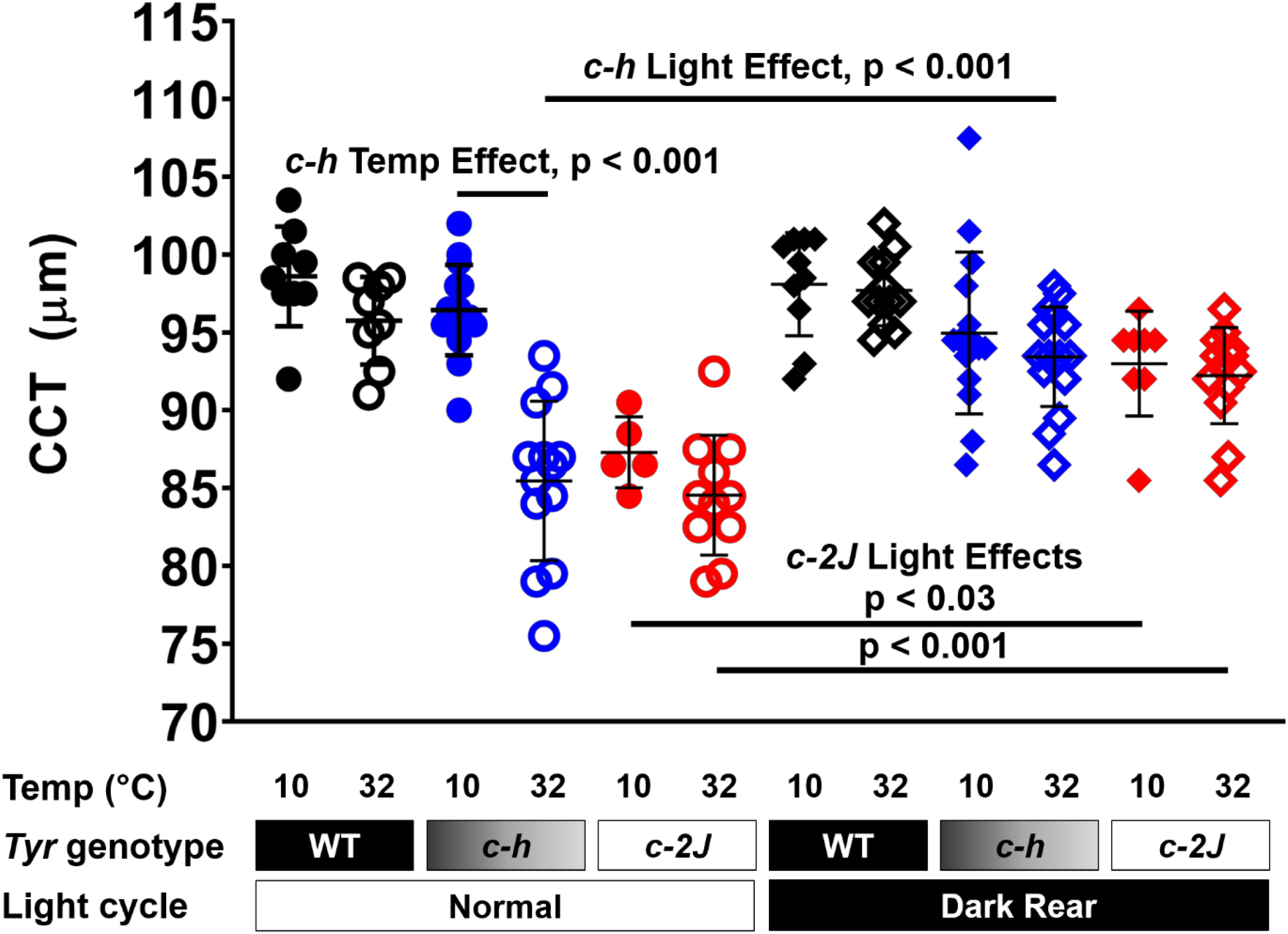
The influence of temperature and light on central corneal thickness (CCT) for the B6, B6.*Tyr*^*c-h*^, and B6.*Tyr*^*c-2J*^ strains of mice. Each point on the graph represents the average CCT measured by optical coherence tomography from one adult mouse, 10 to 15 weeks old, with age-matching across genotypes and *error bars* = mean ± standard deviation. Cohorts were subjected to standard lighting or dark-rearing. Each grouping of *Tyr* genotypes (WT B6 = *black*; *c-h* temperature-sensitive himalayan = *blue*; *c-2J* albino = *red*) has mice raised at 10°C or 32°C (one-way ANOVA with Sidak test).

Regarding light, comparison of B6.*Tyr*^*c-h*^ mice, as well as B6 and B6.*Tyr*^*c-2J*^ controls, raised at 10°C vs. 32°C in environmental control chambers with dark-rearing of mice from conception to 10–15 weeks of age rescued the thin CCT phenotype associated with albinism (*p* < 0.001 for [*Tyr*^*c-h*^ at 32°C standard light] vs. [*Tyr*^*c-h*^ at 32°C dark rear]; *p* < 0.03 for [*Tyr*^*c-2J*^ at 10°C standard light vs. *Tyr*^*c-2J*^ at 10°C dark rear]; *p* < 0.001 for [*Tyr*^*c-2J*^ at 32°C standard light vs. *Tyr*^*c-2J*^ at 32°C dark rear]; *n* = 5–16 per condition; one-way ANOVA with Sidak test; Figure 7; Figure 7 – Source Data 1−2). Rearing pigmented B6 mice in constant light is known to lead to an increase in axial length^33^. OCT examinations of independent cohorts of C57BL/6J and *Tyr*^*c-2J*^ mice for the purpose of measuring axial length show that *Tyr*^*c-2J*^ mice have a 65.6 μm greater axial length on average compared to C57BL/6J mice (3.453 mm vs. 3.388 mm, respectively; *p* < 0.001, Student’s two-tailed *t*-test; *n* = 10 male and 10 female mice per strain; Figure 8; Figure 8 – Source Data 1−2).

**Figure 8.**
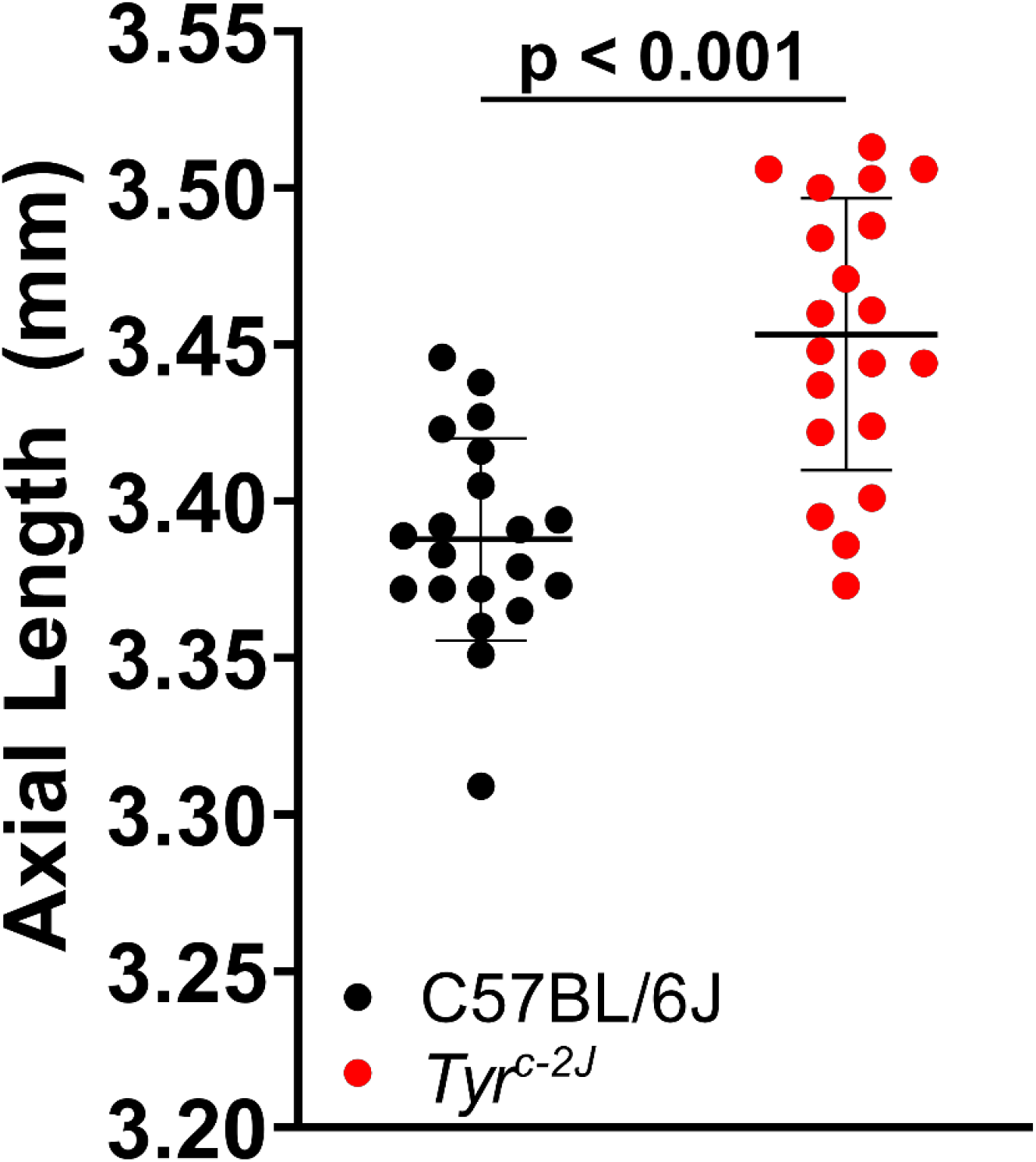
The influence of *Tyr* on ocular axial length. Each point on the graph represents the average axial length measured by optical coherence tomography from one 12-week-old adult mouse and *error bars* = mean ± standard deviation. The plot compares axial length between the B6 and B6.*Tyr*^*c-2J*^ strains of mice, measured from the outer surface of the cornea to the retinal pigment epithelium/choroid interface (Student’s two-tailed *t*-test).

## Discussion

Using a phenotype-driven quantitative genetic analysis of CCT, physical mapping led to identification of a small critical region containing 10 genes, of which we ruled out three (*Fzd4, Rab38, Ctsc*) and found via an analysis of a *Tyr* allelic series (*c, c-2J, c-h*) that *Tyr* is the causative gene underlying the *Cctq1a* QTL. *Tyr* is by no means an unknown gene—it was in fact one of the first known mammalian genes whose initial discovery predates the word “gene”^27,34^,35. However, *Tyr* was a surprising gene to find linked to CCT. TYR is an oxidase whose only known biological role relates to melanin synthesis^27,36^.

Although there are small numbers of pigmented cells in the corneal limbus, the cornea is by and large not only non-pigmented, but transparent. Furthermore, previously published data indicate that *Tyr* is not even expressed in the adult cornea^18^, which is consistent with the current study which also found near-zero expression in the cornea at 3 weeks of age (RNA-Seq max expression ∼1.3 FPKM; Supplementary Data 7). Thus, there is no molecular rationale for proposing that TYR has a direct function in corneal cells. If not for the current experiments, there would also be sparse biological rationales for proposing that TYR might influence corneal anatomy through any mechanism. Regardless, the current agnostic QTL study led to the conclusion that *Tyr* contributes to the primary genetic influence on CCT, at least in the context of KS x SJL hybrids. Experiments using multiple alleles on matched genetic backgrounds, in isolation and genetic complementation crosses, conclusively confirmed allelism of *Tyr-Cctq1a*; albino mice lacking TYR function have thin corneas. Rationalizing that albinism would expose the developing eye to increased light, one of the mechanistic experiments performed here compared the effect of dark-rearing on the CCT of albino vs. pigmented C57BL/6J mice. The results showed that the thin CCT phenotype of albino B6.*Tyr*^*c-h*^ mice raised at 32°C and B6.*Tyr*^*c-2J*^ mice was rescued by dark-rearing. Thus, we are led to propose an epiphenomenon, whereby developmental light exposure interacts with genotype as an important determinate of corneal thickness (Figure 9).

**Figure 9.**
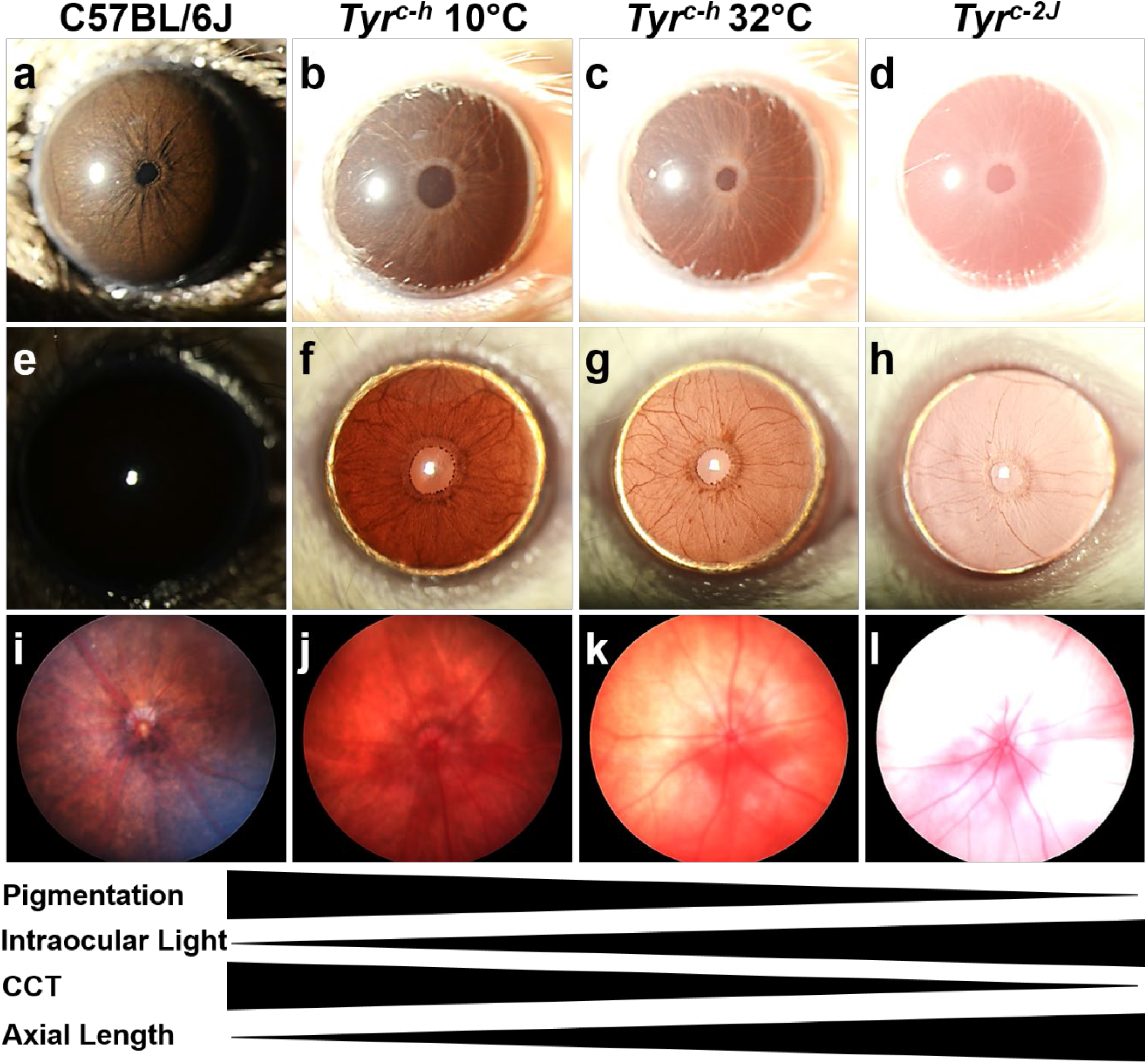
Model of the influences of genotype and light exposure on ocular development. **a-d** Slit-lamp images photographed with identical camera settings showing eyes from **a** B6 mice, **b** *Tyr*^*c-h*^ mice reared at 10°C, **c** *Tyr*^*c-h*^ mice reared at 32°C, and **d** *Tyr*^*c-2J*^ mice. **e-h** Slit-lamp iris transillumination images photographed with identical camera settings showing eyes from **e** B6 mice, **f** *Tyr*^*c-h*^ mice reared at 10°C, **g** *Tyr*^*c-h*^ mice reared at 32°C, and **h** *Tyr*^*c-2J*^ mice. **i-l** Micron retinal fundus images photographed with identical camera settings showing eyes from **i** B6 mice, **j** *Tyr*^*c-h*^ mice reared at 10°C, **k** *Tyr*^*c-h*^ mice reared at 32°C, and **l** *Tyr*^*c-2J*^ mice. CCT– central corneal thickness.

All current CCT measurements were done with mice 10–15 weeks of age, which is slightly past the age at which the cornea of B6 mice reaches its final adult thickness (∼P55)^37^. Transcriptomic changes in the cornea related to *Tyr* genotype were detectable at 3 weeks of age. Thus, the timeframe for when *Tyr* can impact the cornea is presumably during anterior chamber development at some point preceding 3 weeks of age. Two mechanisms, which are both conjectural, might feasibly contribute to this early acting phenomenon. 1) Corneal development might be a component of refractive development. Emmetropization typically occurs in the first months following eyelid opening, with impacts from both the amount of light and its focus on the retina^33,38^. Thus, it is feasible that albinism could in effect cause blur (from light not being absorbed by melanin and reflecting within the eye), which induces relative myopia and CCT thinning with rearing in normal lights, but not in dark-rearing. 2) Corneal development might be influenced by central or corneal circadian outputs. Circadian outputs arise from both central and peripheral clocks, with the suprachiasmatic nucleus (SCN) being a central master pacemaker that receives light signals from retinal ganglion cells and subsequently coordinates phasing to peripheral tissues^39,40^. Notably, the SCN receives retinal input from the retinohypothalamic tract, which is known to be expanded in albinos^41-43^. Thus, one possibility is that factors increasing SCN output (such as albinism or cycling light conditions) might lead to decreased CCT and those decreasing the signaling (such as dark rearing) might lead to increased CCT. Refractive and circadian mechanisms could also be acting in an intertwined way^44^. Additional experiments are needed to distinguish these, and possibly other, mechanisms relevant to our current findings.

A leading candidate for contributing to the molecular mechanism causing thin CCT in albino mice was DOPA, which is a cofactor for TYR^27,36^, a substrate for TH leading to dopamine (reviewed in ^28^), and an endogenous ligand for the G-protein-coupled receptor GPR143^45^. DOPA can modulate refractive development^27, 29, 40^ and the circadian system^46^, as well development of multiple ocular tissues^47,48^. Notably, mice with a conditional knock-out of *Th* in the retina have decreased CCT^30^. The current experiments with *Tyr* mutant mice were not able to detect a role for DOPA in influencing CCT of these albino strains, though an important caveat to point out is that only a single dosing schedule for DOPA supplementation was currently tested.

It is unclear whether albinism or pigmentation influences CCT in humans, though our current experiments suggest this is likely. In humans, loss of function mutations in *TYR* cause oculocutaneous albinism type 1 (OCA 1)^36,49^. It’s unclear from the literature if humans with OCA 1 have decreased CCT; it may be difficult to ascertain because of relatively small patient populations and confounding variables such as eye rubbing^50^.

The current study has several implications with respect to mouse genetics and mouse models of disease: 1) The results highlight the potential for environmental influences on ocular development, which have been quantitated here for CCT, but may extend to other tissues as well. 2) The results indicate that albino mice should be tested for potentially being a naturally occurring model of myopia. 3) Because CCT is a complex trait, there is little reason to suspect that different inbred albino mouse strains would necessarily have the thinnest CCT in comparison to other inbred strains that are pigmented^51^, only that they would have thinner CCT as albino mice compared to pigmented mice *within* an inbred strain. However, for any experiment using such an albino strain, or cohorts in which an allele such as the common *Tyr*^*c*^ allele is segregating, attention to the possibility of environmental influences is warranted. 4) *Tyr* has been linked with many ophthalmic traits in mice^41-43,47,52-57^; in some instances, a consideration of gene-environment interactions in the mechanism of these various models may be warranted. And finally, 5) our study uncovers a genetic peculiarity. *Cctq1* was originally reported as a single QTL on chromosome 7, detected in an F2 intercross of KS and SJL inbred mice^16^. In studies of successive generations of congenic mice, N4F2 mice heterozygous for the *Cctq1* alleles showed the original differential phenotype (over-dominant, *increased* CCT compared to littermate controls)^16^, whereas in N10–N15 intercrosses, mice homozygous for the SJL allele showed the differential phenotype (recessive, *decreased* CCT compared to littermate controls). As the sub-congenics were intercrossed and analyzed for CCT, KS.SJL-*Cctq1a*^*SJL*^*;Cctq1b*^*KS*^ mice consistently had thinner corneas than KS.SJL-*Cctq1a*^*KS*^*;Cctq1b*^*KS*^ mice of the same generation, but there were also fluctuations in absolute value related to generation and interval size. The likely explanation for these observations is that there was more than one CCT-modifying gene in the original *Cctq1* interval, which was in fact found to be the case by the resolution of *Cctq1* into *Cctq1a* and *Cctq1b*. This is consistent with the findings of human GWAS, indicating that there are likely hundreds of CCT-influencing genes dispersed throughout the genome, many of which will be physically close to one another. In mice, it is known that as a locus is narrowed using congenics, genes can be segregated away from nearby modifiers and the overall phenotype of the original QTL can be reduced, disappear, or reverse its apparent effect^58-60^. The current data seem to exemplify this phenomenon.

In summary, our phenotype-driven genetic study of CCT identified *Tyr* as a significant regulator of CCT in mice. The molecular findings of this study were unexpected. We propose that the results can be explained by an epiphenomenon whereby a gene:environment interaction; i.e., *Tyr*-mediated albinism allowing increased exposure of the eye to light has an important influence on corneal development.

## Materials and Methods

### Experimental animals

All animals were treated in accordance with the ARVO Statement for the Use of Animals in Ophthalmic and Vision Research. The majority of mice were housed and bred at the University of Iowa Research Animal Facility with approval for experimental protocols conferred by the Institutional Animal Care and Use Committee of the University of Iowa. Two cohorts of mice, the C57BL/6J (JAX Stock No. 000664) and B6.Cg-*Tyr*^*c-2J*^/J (JAX Stock No. 000058) used for in vivo axial length measurements (Figure 8) were purchased from The Jackson Laboratory at 10-weeks-old and subsequently housed at the University of California San Francisco until 12-weeks-old with approval for experimental protocols conferred by the Institutional Animal Care and use Committee at the University of California San Francisco. Mouse strains used in this study include: SJL/J (Stock No. 000686), C57BLKS/J (JAX Stock No. 000662), C57BL/6J (JAX Stock No. 000664), B6.Cg-*Tyr*^*c-2J*^/J (JAX Stock No. 000058), B6;129-*Fzd4*^*tm1Nat*^/J (JAX Stock No. 012823), B6.Cg-*Tyr*^*c-h*^/J (JAX Stock No. 000104; Imported from Dr. Brian Brooks at the NIH), B6.Cg-*Ctsc*^*tm1Ley*^ (Imported from Dr. Christine Pham at Washington University), KS.Cg-SJL^*Cctq1*^, B6-*Ctsc*^*tm1Mga*^, and B6-*Tyr*^*tm4Mga*^. All experiments included male and female mice.

### Chromosome 7 QTL analysis

The chromosome 7 quantitative trait locus analysis was performed with R/qtl, using the two-dimensional genome-wide scan (scantwo). Significance thresholds were determined empirically by permutation testing, using 1000 permutations. The validity of a multiple QTL model was tested by performing a multiple regression analysis. Phenotypic variance was estimated and the full model was statistically compared to reduced models in which one QTL was dropped.

### Constructing congenic mice

A ∼38.3cM genomic region (*i*.*e*., *Cctq1*) spanning from *D7Mit318* (SSLP marker at 42.3 cM) to *rs13479545* (SNP marker at 81.2 cM) was transferred from SJL/J mice (abbreviated throughout as SJL; thick cornea) onto the genetic background of C57BLKS/J mice (abbreviated throughout as KS; thin cornea) by reiterative backcrossing. Mice carrying the SJL alleles within the region (*i*.*e*., KS.SJL-*Cctq1*^*Het*^) were selected at each generation by using a panel of six markers that were tested and found to be polymorphic between KS and SJL mice. At the N10 generation, mice were intercrossed. At this point, *Cctq1* was treated as a digenic locus, renamed to *Cctq1a* and *Cctq1b* (see Results and Figure 2). *Cctq1a* encompassed the region spanning from *D7Mit318* (42.3 cM) to *D7Mit220* (55.7 cM). *Cctq1b* spanned from *D7Mit105* (70.3 cM) to *rs13479545* (81.2 cM). All genotype combinations of *Cctq1a* and *Cctq1b* (9 possible combinations; *i*.*e*., homozygosity for KS alleles, heterozygous, or homozygosity for SJL alleles at each locus) were analyzed for their effect on CCT.

Sub-congenic mice harboring reduced *Cctq1a* intervals (KS alleles at *Cctq1b*) were also generated. At N12, *Cctq1a* was reduced to a 15.9 cM region spanning from *D7mit347* to *D7mit321*. These N12 sub-congenic mice are referred to throughout as KS.SJL-*Cctq1a*(15.9cM). At N15, *Cctq1a* was reduced to a 9.9 cM region spanning from *rs3672782* to *D7mit321*. These sub-congenic N15 mice are referred to throughout as KS.SJL-*Cctq1a*(9.9cM). Sub-congenic mice were intercrossed, and all genotypes were assessed for the CCT phenotype. The KS.SJL-*Cctq1* line has been sperm cryopreserved.

### CCT phenotyping

All measurements were recorded from adult mice. Cohorts were determined based on either 1) mice received from The Jackson Laboratory, or 2) genotypes of mice born in the first litters of a genetic cross. Mice were injected with a standard mixture of ketamine/xylazine (intraperitoneal injection of 100 mg ketamine + 10 mg xylazine / kg body weight). During induction of anesthesia, mice were provided supplemental indirect warmth by a heating pad. Immediately following anesthesia, eyes were hydrated with balanced salt solution (BSS) and corneal images were obtained with a Bioptigen optical coherence tomographer (SD-OCT; Bioptigen, Inc., USA). A 12mm telecentric bore was used to image the anterior segment of each eye. The bore was positioned such that the pupil of the eye was centered in the volume intensity projection. Scan parameters were as follows: radial volume scans 2.0 mm in diameter, 1000 A-scans/B-scan, 100 B-scans/volume, 1 frame/B-scan, and 1 volume. Central corneal thickness (CCT) was measured for each eye using vertical angle-locked B-scan calipers by a technician masked to genotype and/or treatment. Mice were included in the analysis if the difference between the right and left eyes was less than 7 μm and if both eyes were free from opacity. The average CCT with standard deviation for each genotype was statistically compared using Student’s two-tailed *t*-test for comparison of two cohorts or one-way ANOVA with a Tukey post-test for comparison of three or more cohorts.

### Recombination mapping

For genetic mapping of the gene underlying *Cctq1a*, additional polymorphic markers were identified and tiled into the region. Intercrosses of N10 mice were continued and mice with informative recombination events were analyzed for the CCT phenotype. Based on the allelic effects of the QTL on the CCT phenotype (see Results), the genomic boundaries of the QTL (and hence, the region of the underlying gene) were deduced by comparing the phenotype of the recombinant mice with the location of the recombination event within the critical interval. The following is a complete list of all the polymorphic markers used for genotyping (listed in order from centromeric to telomeric): *D7Mit318, rs13479346, rs13479362, D7Mit347, D7Mit62, rs6271685, rs108403472, Cctq1a-STR5, D7Mit31, rs3672782, rs32438580, rs3663323, rs13479392, rs13479393, rs6247100, rs13479395, D7Mit301, D7Mit321, D7Mit220, D7Mit238, D7Mit105, rs13479535, rs13479536*, and *rs13479545*. All MIT marker primer sequences are available on the Mouse Genome Informatics website^61^. *Cctq1a-STR5* is a short tandem repeat polymorphism with genotyping primers designed in-house: 1) GCAAATTAATCAGCACATTTC and 2) CATTTCCTCTAGGTGATTGC.

### Exome sequence analysis

High quality genomic DNA was harvested from KS and SJL spleen tissue using a Qiagen DNeasy Blood and Tissue kit following the manufacturer’s instructions; an RNA digestion step was included. DNA samples were sent to BGI Americas for sequencing and passed their quality control standards. Libraries were constructed with an Agilent SureSelect 50Mb Mouse Exome Capture Kit and were sequenced with 50X coverage using an Illumina HiSeq2000. Standard bioinformatics analysis was conducted in which the data were filtered (by removing adaptor contamination and low-quality reads from raw reads), aligned, and SNPs were called using the GRChm38 mouse reference genome. The resulting VCF file was processed at the University of Iowa to annotate the consequence for each variant using a locally-developed pipeline.

### RNASeq analysis

N12F3 KS.SJL-*Cctq1a(15*.*9cM)*^*KS*^, KS.SJL-*Cctq1a(15*.*9cM)*^*SJL*^, and KS inbred mice were euthanized at three weeks of age by cervical dislocation. Immediately upon death, mice were enucleated, and the eyes were placed in RNase free dishes containing RNA*later* RNA stabilization reagent. Corneas from mice were dissected in RNA*later* and pooled to make one sample (6 corneas per sample); three samples were collected per genotype. Cornea samples were either stored at -80°C in RNA*later* or processed immediately. Corneas were transferred from RNA*later* to 0.7 mL of lysis/binding buffer from the *mir*Vana miRNA isolation kit (Ambion) and homogenized for 1 minute using a tissue tearer. The homogenate was then passed through a QIAshredder column (Qiagen) and the lysate was collected. For the remainder of the procedure, the samples were processed using the *mir*Vana kit for total RNA according to the manufacturer’s instructions. The quality and concentration of the RNA was analyzed using a NanoDrop 2000 and the Agilent Model 2100 Bioanalyzer. All samples had RNA integrity numbers of 9.5 or greater, indicating high quality RNA with little degradation of the samples. Samples were barcoded and stranded libraries were prepared by the Genomics Division of the Iowa Institute of Human Genetics. The nine libraries were pooled together, split into two equal parts, and run on two lanes of an Illumina HiSeq to obtain 100 base pair, paired-end sequence reads.

Reads were mapped to the mm10 mouse genome build using Tophat2 (ver 2.0.11^62^). The ‘-r’ parameter was set to 135, and the ‘--no-coverage-search’ option was used. Transcript abundance was quantified using Cufflinks (ver. 2.1^63^) for RefSeq transcript models from the Illumina iGenomes mm10 package. Ribosomal RNA and mitochondrial gene loci, obtained from UCSC Genome Table Viewer, were masked from the Cufflinks analysis, and the ‘--max-bundle-frags’ parameter was set to 20000000. Differential expression between genotypes was performed using Cuffdiff (ver. 2.1.1^64^). Genes identified from each comparison were subsequently filtered for only those with a q-value ≤ 0.001 and a mean FPKM (Fragments Per Kilobase of transcript per Million mapped reads) value ≥ 1 in at least one of the strains. Functional enrichment analysis was performed with WebGestalt [32, 33] with analysis parameters detailed in Supplementary File 2 – Figure Supplement 1.

### Constructing *Ctsc* null mice

B6-*Ctsc*^*KO*^ mice were generated by the Genome Editing Facility at The University of Iowa on a pure C57BL/6J (JAX Stock No. 000664) background by targeting *Ctsc* Exon 1 with CRISPR/Cas9 using guide sequence: CGTGCGCTCCGACACTCCTGCC. Founders were crossed with C57BL/6J mice and offspring analyzed for germline transmission of *Ctsc* mutations. From four founders, we observed transmission of four separate *Ctsc* mutations, all predicted to be null based on Sanger sequencing results. Four *Ctsc* mutations were propagated in separate intercross mouse lines and were validated as null mutations by Western Blot using an antibody against CTSC (Catalog #AF1034; R&D Systems, Inc.; Minneapolis, MN; Supplementary Data 10). One of these mutations, *Ctsc*^*tm1Mga*^, is a 46 bp exon 1 coding sequence deletion that was used for additional downstream analysis.

### Constructing *Tyr* null mice

To generate mice with *Tyr* null mutations on a pure C57BL/6J (JAX Stock No. 000664) background, the Genome Editing Facility at the University of Iowa targeted *Tyr* Exon 1 with CRISPR/Cas9 using two guide sequences simultaneously: 1) CCATGGATGGGTGATGGGAG and 2) TTCAAAGGGGTGGATGACCG. Founders were crossed with C57BL/6J mice and offspring analyzed by Sanger sequencing for germline transmission of *Tyr* mutations. This experiment generated more alleles than we could reasonably work with (15 unique mutations identified via sequencing; Supplementary Data 12). For each unique mutation, we set up a complementation cross with B6.*Tyr*^*c-2J*^ and screened progeny coat color for alleles conferring novel function. F1 progeny were screened for 14 alleles; the mice harboring the remaining allele did not produce F1 progeny. All 14 alleles produced a standard albino coat color in trans with the *Tyr*^*c-2J*^ mutation, indicating failure to complement. Going forward, we chose to complete additional studies for one allele, *Tyr*^*tm4Mga*^, which is a 4 bp deletion in the coding sequence of exon 1, causes a frameshift and leads to a premature stop codon predicted to cause RNA-mediated decay (i.e., a presumed null mutation). Accordingly, homozygotes of this strain are albino.

### L-DOPA supplementation studies

Breeder cages of C57BL/6J, B6.Cg-*Tyr*^*c-2J*^/J, and B6.Cg-*Tyr*^*c-h*^/J were provided with water bottles containing water only (control) or water with 200mg/L of L-DOPA, with 30mg/L of benserazide to minimize the conversion of L-DOPA to dopamine in the peripheral nervous system. Fresh water bottles were prepared and supplied every Monday, Wednesday, and Friday. Litters were weaned into cages where they continued to receive the same treatment of supplemented water until 13–16 weeks old. Cohorts were determined by the number of mice born under each treatment condition.

### Environmental chamber

Breeder cages of C57BL/6J, B6.Cg-*Tyr*^*c-2J*^/J, and B6.Cg-*Tyr*^*c-h*^/J were housed in rodent environmental control chambers (Powers Scientific) either at 10°C or 32°C and with either a standard light cycle (12 h on/12 h off) or dark-rearing. Resulting pups were born and group-housed with age-matched mice regardless of genotype at the altered temperature and light cycle until adulthood, when CCT was analyzed by OCT at 10–15 weeks old. Cohorts were determined by the number of mice born under each treatment condition.

### Axial length phenotyping

Envisu R4300 spectral-domain optical coherence tomography (SD-OCT, Leica/Bioptigen Inc., Research Triangle Park, NC, USA) was employed to measure the ocular axial length in adult (12-week-old) mice^65^. Twenty mice of each genotype were ordered from The Jackson Laboratory and all forty mice were phenotyped. Mice were anesthetized with ketamine/xylazine (100 mg/kg and 5mg/kg, respectively; intraperitoneal) and their eyes dilated before placing the animal in a cylindrical holder. The eye was hydrated with Genteal (Alcon, Fort Worth, TX, USA) and positioned in front of the OCT light source. Correct alignment of the eye was achieved by placing the Purkinje image in the center of the pupil. The images were acquired in rectangular volume and radial volume scans. The axial length was calculated by measuring the distance from the anterior corneal surface to the RPE/choroid interface for both the left and right eyes of a given mouse. Measurements from the left and right eye of each mouse were averaged to give a single measurement per animal. Measurements from all eyes were included in the analysis. To minimize the possible effect of body weight on ocular size, we ensured that body weight of littermates was within a narrow range in each of the comparative groups. The average axial length with standard deviation for each genotype was statistically compared using Student’s two-tailed *t*-test.

### Statistics

Power calculations were performed using the Piface Java applet by Russell V. Lenth^66^ with an analysis for a two-sample *t*-test with an alpha = 0.05 or a one-way ANOVA with an alpha = 0.05. Assumptions entered for CCT phenotype for a two-sample *t*-test are sigma1/sigma2 = 2.5 (the standard deviation in μm of CCT measured previously in naïve adult mice), true difference of means = 8 (the measured CCT difference in μm between KS.SJL-*Cctq1a*^*SJL*^ vs. KS.SJL-*Cctq1a*^*WT*^), and sample size = 4 mice per group, which results in a power of 96.18% to detect an 8 μm difference (7.8% difference in CCT on a B6 background) between groups. Assumptions entered for CCT phenotype for a one-way ANOVA testing three genotypes are SD[WT], SD[HET], SD[MUT] each = 2.5 (the standard deviation in μm of CCT measured previously in naïve adult mice), SD[Residual] = 5.2 (the standard deviation in μm of CCT measured previously between groups with a CCT difference of 10 μm), sample size = 5 mice per group, and a Tukey/HSD post-hoc analysis, which results in a power of 97.84% to detect an 8 μm difference between groups. Assumptions entered for the axial length phenotype for a two-sample *t*-test are sigma1/sigma2 = 0.25 (the standard deviation in mm of axial length measured previously in naïve adult mice), true difference of means = 0.33 (10% difference in axial length in mm), and sample size of 20 mice per group, which results in a power of 98.24% to detect a 0.33 mm difference between groups. If additional mice resulted from genetic crosses, phenotypes were collected and included in data analyses. A multiple regression analysis, in which each locus and the interaction component were sequentially dropped from the 2-QTL model, was used to analyze the presence of two interacting loci on chromosome 7. Student’s two-tailed *t*-test was used to evaluate the difference between two independent genotypes for CCT (*t*-values and degrees of freedom for each comparison are listed in Figure 3 – Source Data 3) and axial length (*t*-value = 5.415; degrees of freedom = 38). A one-way ANOVA with a Tukey post-test was used to evaluate the difference between three or more independent genotypes for CCT (*f*-values and degrees of freedom for each comparison are listed in Figure 3 – Source Data 3). A one-way ANOVA with Sidak test was used to evaluate four comparisons to determine the effect of two environmental conditions on CCT in B6.cg-*Tyr*^*c-h*^ and B6.cg-*Tyr*^*c-2J*^ mice (f-value and degrees of freedom are listed in Figure 7 – Source Data 2) because previous experiments identified significant differences between *Tyr* genotypes. No outliers were excluded.

## ACKNOWLEDGEMENTS

This work was supported in part by Merit Review Award (101RX001481) from the U.S. Departments of Veterans Affairs RR&D Service to MGA and by internal faculty development funds to MGA. DRL was supported in part by a NIH/NEI training grant, EY021436. AHB was supported in part by a NIH/NIDDK training grant, DK112751. Equipment maintenance was supported in part by the University of Iowa Multidisciplinary Investigations in Visual Science service cores (a resource supported by a P30 grant from the NIH/NEI to the University of Iowa, EY025580). We acknowledge use of the University of Iowa Genome Editing Core Facility (a core resource supported in part by grants from the NIH and from the Roy J. and Lucille A. Carver College of Medicine) directed by William Paradee, PhD, and we thank Norma Sinclair, Patricia Yarolem and Joanne Schwarting for their technical expertise in generating transgenic mice. The authors thank Karl Broman for analyzing chromosome 7 of our original F2 dataset with us. The contents do not represent the views of the U.S. Department of Veterans Affairs or the U.S. Government.

The authors declare no competing interests.

## AUTHOR CONTRIBUTIONS

K.J.M., D.R.L., and M.G.A. conceived and designed the experiments. K.J.M., D.R.L., C.J.V., A.H.B., L.M.D., N.P., H.E.M., M.N.A., and M.G.A. performed the experimental work. K.J.M., D.R.L., and M.G.A. performed experimental data analysis and interpreted the results with contributions from all authors. S.S.W. analyzed the RNA-seq data supervised by T.E.S. W.J.P. constructed the transgenic mice. S.K. performed the axial length experiment supervised by K.S.N. K.W. verified and performed statistical analyses. K.J.M., D.R.L. and M.G.A. wrote the manuscript with input from all authors. M.G.A. supervised the study and provided funding.

## FIGURE SUPPLEMENT LEGEND

**Table 1 – Table Supplement 1.**
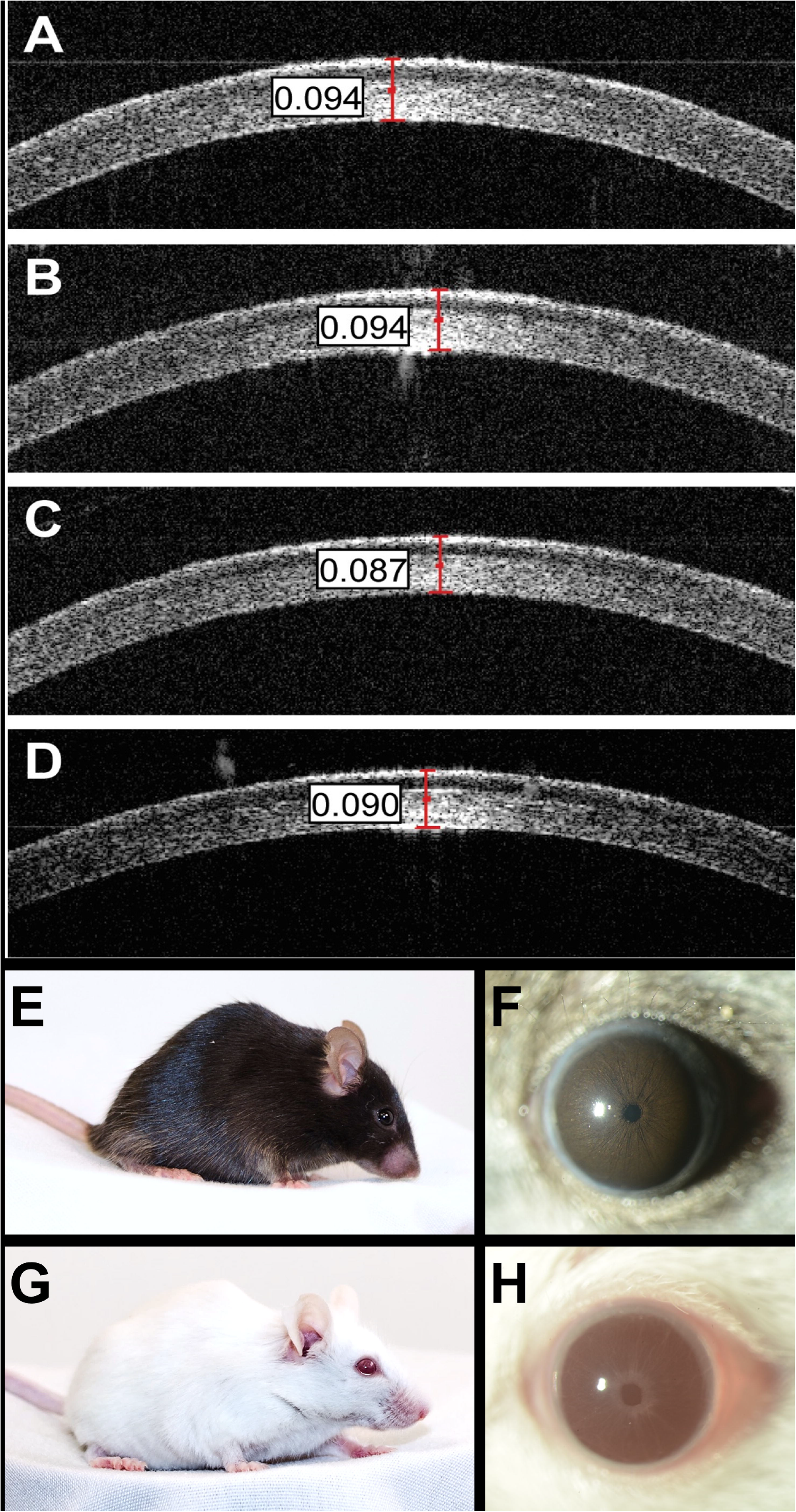
Imaging of KS.SJL-*Cctq1a* mice. Representative optical coherence tomography images with caliper measurements (in millimeters) of central corneas of (**A**) KS inbred mice, (**B**) KS.SJL-*Cctq1a*^*KS*^,*Cctq1b*^*KS*^ N10F2 “congenic control” mice, (**C**) KS.SJL-*Cctq1a*^*SJL*^ N10F2 mice, and (**D**) KS.SJL-*Cctq1a*^*KS*^,*Cctq1b*^*SJL*^ N10F2 mice. Representative (**E, G**) photographs and (**F, H**) slit-lamp images of (**E, F**) KS.SJL-*Cctq1a*^*KS*^,*Cctq1b*^*KS*^ and (**G, H**) KS.SJL-*Cctq1a*^*SJL*^,*Cctq1b*^*KS*^ congenic mice.

**Supplementary File 2 – Figure Supplement 1.**
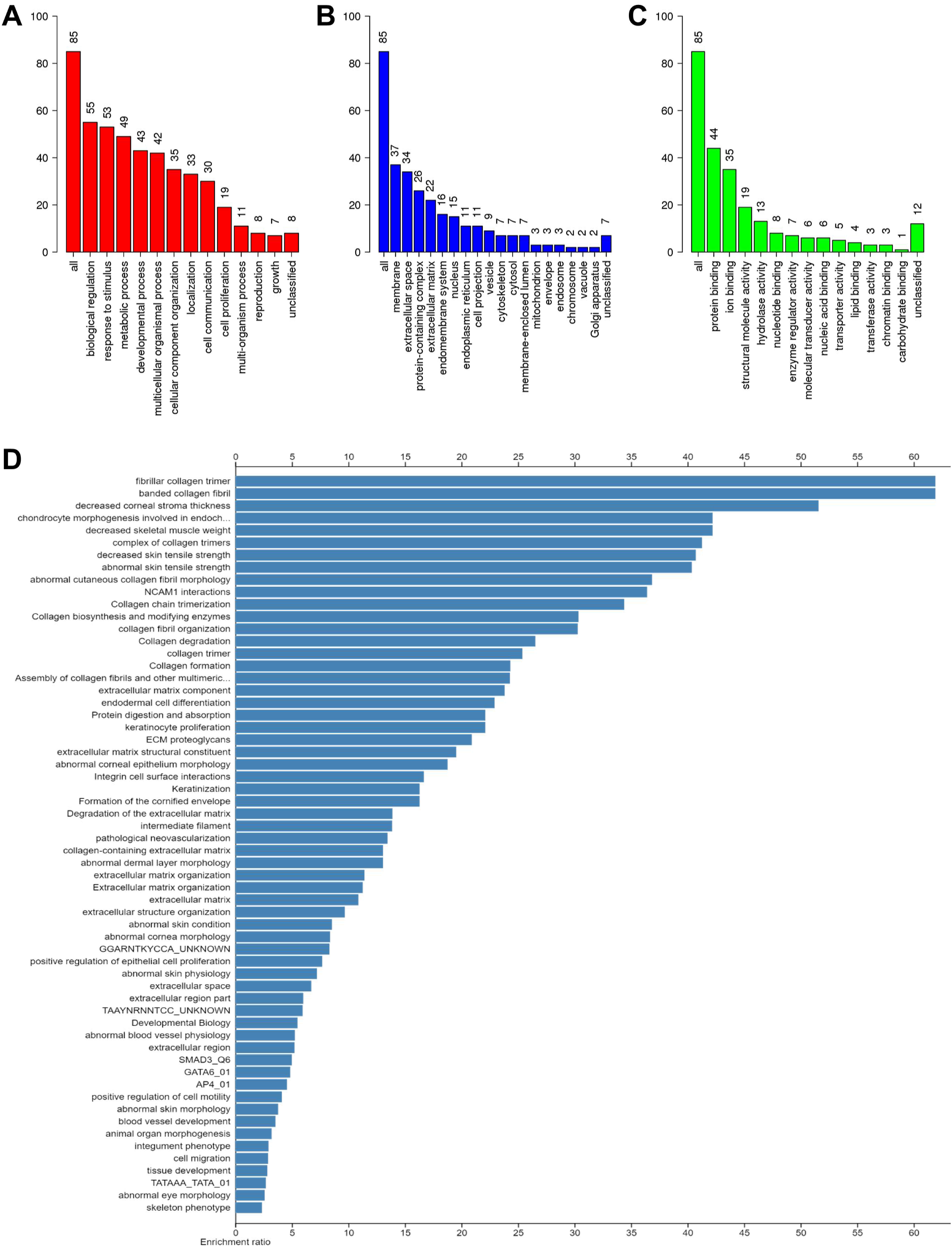
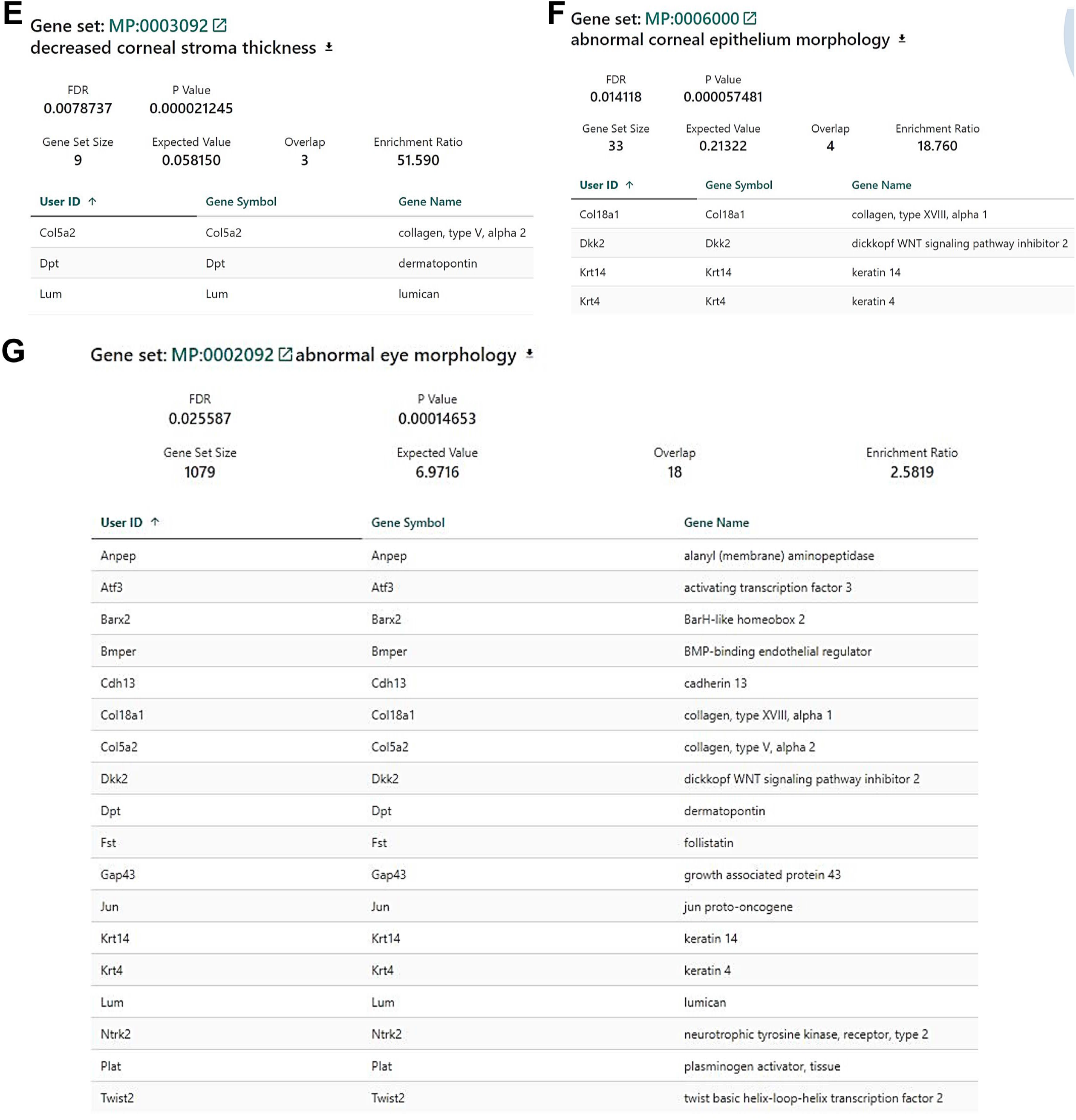
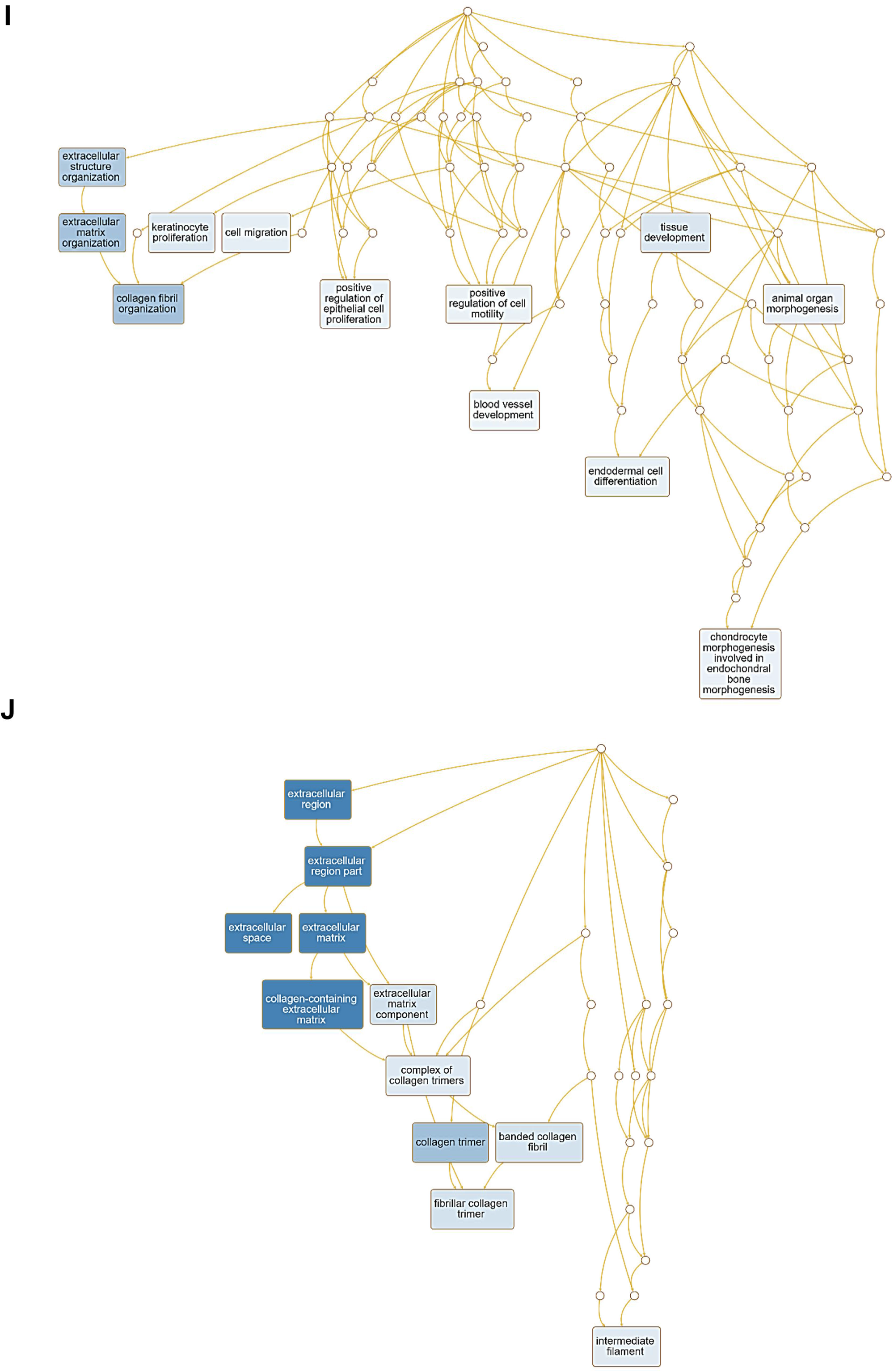
WebGestalt over-representation analysis output. Parameters for the over-representation analysis are as follows: 1) Minimum of 5 gene IDs in the category, 2) Maximum of 2,000 gene IDs in the category, 3) False Discovery Rate with Benjamini-Hochberg correction, and 4) False Discovery Rate significance level of < 0.05. The interesting gene input list contains 87 user IDs in which 85 user IDs are unambiguously mapped to 85 unique entrezgene IDs and 2 user IDs cannot be mapped to any entrezgene ID. The GO Slim summary is based upon the 85 unique entrezgene IDs. Among 85 unique entrezgene IDs, 83 IDs are annotated to the selected functional categories and in the reference list, which are used for the over-representation analysis. The reference list, containing genes expressed in the cornea by RNASeq and listed in Supplementary File 3, can be mapped to 13,404 entrezgene IDs and 12,846 IDs are annotated to the selected functional categories that are used as the reference for the over-representation analysis. The interesting gene list of 85 unambiguously mapped unique entrezgene IDs are as follows: *4833423E24Rik*; *Il33*; *Plvap*; *C130074G19Rik*; *Alox12e*; *Col18a1*; *Car12*; *Has2*; *Krt17*; *Krt4*; *Aadac*; *Mmp3*; *Barx2*; *Spink5*; *Loxl4*; *Lgi2*; *Krt8*; *Krt19*; *Il1rn*; *Trpm6*; *Krt14*; *Atf3*; *Serpinb3a*; *Nr4a1*; *Car13*; *Fmo2*; *Fst*; *Col14a1*; *Atp12a*; *Kcnh1*; *Cdh3*; *Jun*; *Plat*; *Adam28*; *Krt75*; *Cdh13*; *Anpep*; *Ppic*; *Col11a1*; *Rasl11b*; *Col16a1*; *Col6a3*; *Twist1*; *Adamts2*; *Ctsc*; *Dkk2*; *Nxph4*; *Plod1*; *Mme*; *Upk3b*; *Sema3c*; *Fibin*; *Gap43*; *Spon1*; *Itih2*; *Vcam1*; *Srpx*; *Matn4*; *Bmper*; *C1ql2*; *Smoc2*; *Adcyap1r1*; *C1qtnf7*; *Agtr2*; *Mfap4*; *Ntrk2*; *Olfml3*; *Sparcl1*; *Tmem132c*; *Lum*; *Col5a2*; *Cpne7*; *Twist2*; *Gsg1l*; *Dpep1*; *A2m*; *Angptl7*; *Col6a1*; *Abca8b*; *Col5a1*; *Cd40*; *Mamdc2*; *Abca8a*; *Gm2115*; *Dpt*. Based on the above parameters, 124 categories are identified as over-represented categories, in which the 40 most significant categories and representatives in the reduced sets are shown in this report. The Gene Ontology Slim summary for the uploaded interesting gene IDs is shown for (**A**) Biological Process, (**B**) Cellular Component, and (**C**) Molecular Function. The results of over-representation analysis are shown in panel (**D**). Gene sets from the over-representation analysis are shown for select categories: (**E**) decreased corneal stroma thickness, (**F**) abnormal corneal epithelium morphology, (**G**) abnormal eye morphology, and (**H**) abnormal cornea morphology. (**I**) Biological process directed and (**J**) cellular compartment directed acyclic graphs.

## SOURCE DATA FILES

**Figure 1 – Source Data 1**. Table formatted for analysis with R/qtl containing genotypes and central corneal thickness measurements of KS x SJL F2 mice.

**Figure 1 – Source Data 2**. Multiple regression analysis for *Cctq1a* and *Cctq1b* comparing the full model to a model in which the indicated QTL or interaction is omitted.

**Table 1 – Source Data**. Table containing *Mrdq1a* and *Mrdq1b* interval genotypes for individual mice along with corresponding central corneal thickness measurements in KS.SJL-*Cctq1* N10F2 mice.

**Figure 2 – Source Data**. Table containing genotypes from N10F2+ mice harboring a recombination event within the *Mrdq1a* interval.

**Figure 3 – Source Data 1**. Table containing central corneal thickness, corneal epithelium thickness, and corneal stroma thickness measurements for all individual mice. Genetic crosses are separated into individual tabs.

**Figure 3 – Source Data 2**. Graphs of corneal epithelium and stroma thickness for the same mice shown in figure 3.

**Figure 3 – Source Data 3**. Table containing CCT results and statistics for all experimental iterations.

**Figure 3 – Source Data 4**. CRISPR targeting strategy, sanger sequencing of novel mutations, and western blot for *Ctsc*.

**Figure 3 – Source Data 5**. Photographs and slit-lamp images of *Fzd4*^*tm1Nat*^/*Cctq1a* F1 mice.

**Figure 4 – Source Data 1**. CRISPR targeting strategy and sanger sequencing of novel mutations for *Tyr*.

**Figure 4 – Source Data 2**. Table containing central corneal thickness, corneal epithelium thickness, and corneal stroma thickness measurements for all individual mice.

**Figure 4 – Source Data 3**. Graphs of corneal epithelium and stroma thickness for the same mice shown in figure 4.

**Figure 4 – Source Data 4**. Table containing CCT results and statistics for all experimental iterations.

**Figure 5 – Source Data 1**. Table containing central corneal thickness measurements and treatment type for all individual mice. Data from C57BL/6J, B6.*Tyr*^*c-h*^, and B6.*Tyr*^*c-2J*^ are presented in individual tabs.

**Figure 5 – Source Data 2**. Table containing CCT results and statistics for all experimental iterations.

**Figure 6 – Source Data**. Table containing CCT results and statistics for all experimental iterations.

**Figure 7 – Source Data 1**. Table containing central corneal thickness measurements for all individual mice, rearing temperature, rearing light cycle, and strain.

**Figure 7 – Source Data 2**. Table containing CCT results and statistics for all experimental iterations.

**Figure 8 – Source Data 1**. Table containing axial length measurements for all individual mice.

**Figure 8 – Source Data 1**. Table containing CCT results and statistics for all experimental iterations.

## SUPPLEMENTARY FILES

**Supplementary File 1**. Exome results for SJL/J and C57BLKS/J.

**Supplementary File 2**. Results of cornea RNASeq from 3-week-old C57BLKS/J and KS.SJL-*Cctq1a*.

## Notes

### Competing Interest Statement

The authors have declared no competing interest.

