## Supplementary material for "*Tyr* is Responsible for the *Cctq1a* QTL and Links Developmental Environment to Central Corneal Thickness Determination": Figure 1 -- Source Data 2

**Multiple regression analysis comparing the full model to a model in which the indicated QTL or interaction is omitted**

| QTL*^a^* | df*^b^* | Type III SS*^c^* | LOD(λ)*^d^* | %var*^e^* | F value*^f^* | Pvalue(χ2)*^g^* | Pvalue(F)*^h^* |
| --- | --- | --- | --- | --- | --- | --- | --- |
| 7@49.0 | 6 | 702.2 | 8.252 | 30.78 | 7.075 | < 0.0001 | 4.17e-06 |
| 7@74.3 | 6 | 557.1 | 6.804 | 24.42 | 5.613 | < 0.0001 | 6.29e-05 |
| 7@49.0:7@74.3 | 4 | 499.2 | 6.196 | 21.88 | 7.545 | < 0.0001 | 3.10e-05 |

*^a^* Chromosome and centimorgan position of the QTL

*^b^* Degrees of freedom

*^c^* Type III sum of squares

*^d^* log_10_ likelihood ratio comparing the hypothesis of a QTL at position λ versus that of no QTL

*^e^* Phenotypic variance (%) attributed to the indicated QTL or interaction

*^f^* F statistic

*^g^* *p* value for the χ2 distribution

*^h^* *p* value for the F statistic
