## Supplementary material for "*Tyr* is Responsible for the *Cctq1a* QTL and Links Developmental Environment to Central Corneal Thickness Determination": Figure 3 -- Source Data 2

**Testing the influence of lead *Cctq1a* positional gene candidates on central corneal epithelium thickness.** Each point on the graph represents the epithelial thickness from one mouse and *error bars* = mean ± standard deviation. A) Homozygosity of *Cctq1a^SJL^* in KS.SJL-*Cctq1a* N15F7 sub-congenic mice has no effect on epithelium thickness compared to littermate controls having *Cctq1a^KS^* genotypes. B) The *Fzd4^tm1Nat^* allele has no effect on epithelium thickness, in neither the heterozygous nor homozygous state, compared to *Fzd4^WT^* littermate controls. C) The *Fzd4^tm1Nat^* allele in trans with a *Cctq1a^SJL^* allele has no effect on epithelium thickness compared to littermate controls having a *Fzd4^WT^* allele in trans with a *Cctq1a^SJL^* allele. D) Homozygosity of the *Ctsc^tm1Ley^* allele results in a significantly decreased epithelium thickness compared to littermate controls. E) The *Ctsc^tm1Ley^* allele in trans with a *Cctq1a^SJL^* allele results in a significantly decreased epithelium thickness compared to littermate controls having a *Ctsc^tm1Ley^* allele in trans with a *Cctq1a^KS^* allele. F) *Ctsc^KO^* mice on a pure B6 background, harboring one of four *tmMga* alleles predicted to result in a null protein, have an unchanged epithelium thickness compared to *Ctsc^WT^* littermate controls. G) The *Ctsc^tm1Mga^* allele, made on a pure B6 background, in trans with a *Cctq1a^SJL^* allele has no effect on epithelium thickness compared to littermate controls having a *Ctsc^WT^* allele in trans with a *Cctq1a^SJL^* allele. H) Homozygosity of the *Tyr^c-2J^* allele has no effect on epithelium thickness compared to C57BL/6J mice. I) The *Tyr^c-2J^* allele in trans with a *Cctq1a^SJL^* allele has no effect on epithelium thickness compared to littermate controls having a *Tyr^c-2J^* allele in trans with a *Cctq1a^KS^* allele.

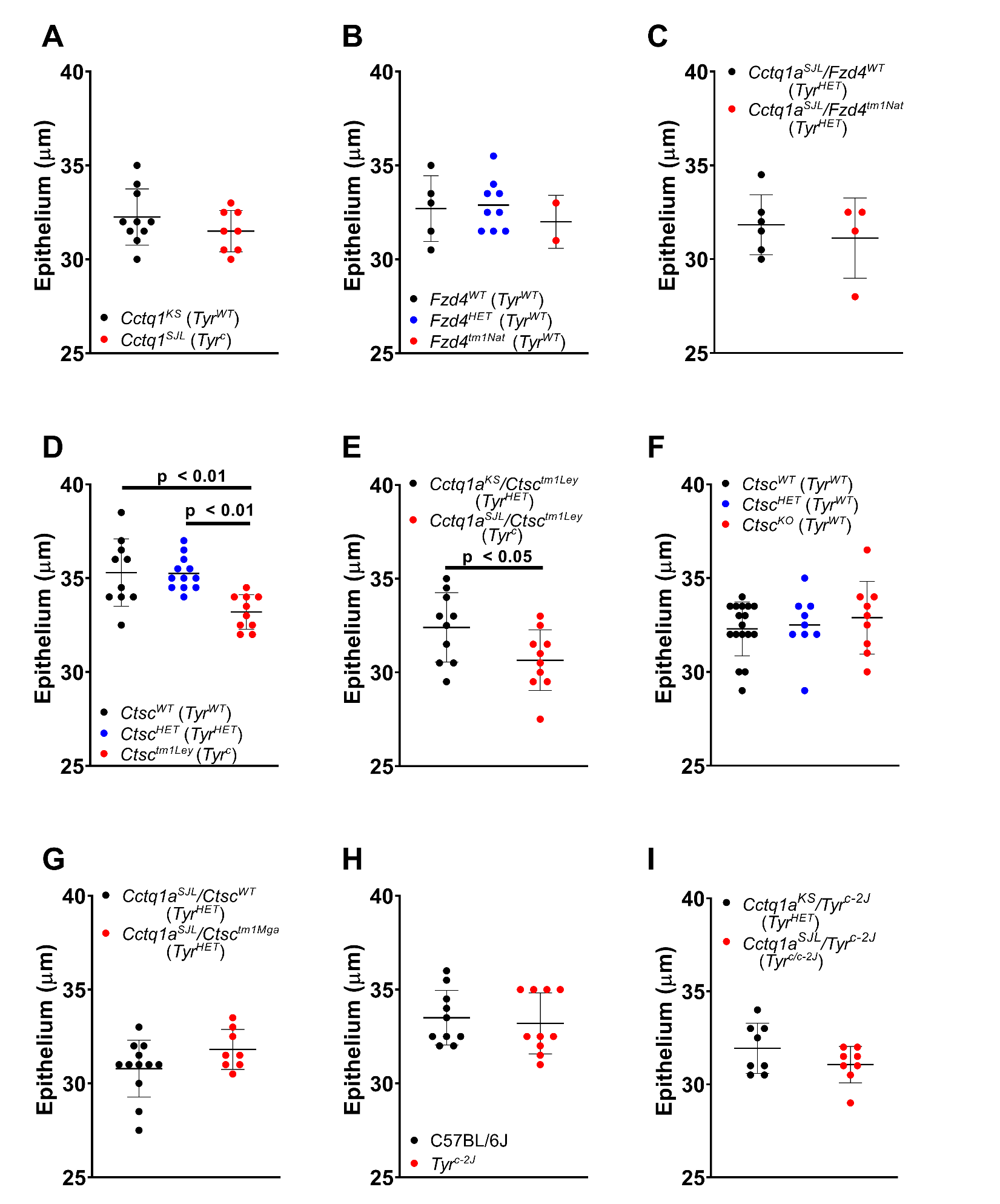

**Testing the influence of lead *Cctq1a* positional gene candidates on central corneal stroma thickness.** Each point on the graph represents the stromal thickness from one mouse and *error bars* = mean ± standard deviation. A) Homozygosity of *Cctq1a^SJL^* in KS.SJL-*Cctq1a* N15F7 sub-congenic mice results in a significantly decreased stroma thickness compared to littermate controls having *Cctq1a^KS^* or genotypes. B) The *Fzd4^tm1Nat^* allele has no effect on stroma thickness, in neither the heterozygous nor homozygous state, compared to *Fzd4^WT^* littermate controls. C) The *Fzd4^tm1Nat^* allele in trans with a *Cctq1a^SJL^* allele has no effect on (complements) stroma thickness compared to littermate controls having a *Fzd4^WT^* allele in trans with a *Cctq1a^SJL^* allele. D) Homozygosity of the *Ctsc^tm1Ley^* allele results in a significantly decreased stroma thickness compared to littermate controls. E) The *Ctsc^tm1Ley^* allele in trans with a *Cctq1a^SJL^* allele results in a significantly decreased (fails to complement) stroma thickness compared to littermate controls having a *Ctsc^tm1Ley^* allele in trans with a *Cctq1a^KS^* allele. F) *Ctsc^KO^* mice on a pure B6 background, harboring one of four *tmMga* alleles predicted to result in a null protein, have an unchanged stroma thickness compared to *Ctsc^WT^* littermate controls. G) The *Ctsc^tm1Mga^* allele, made on a pure B6 background, in trans with a *Cctq1a^SJL^* allele has no effect on (complements) stroma thickness compared to littermate controls having a *Ctsc^WT^* allele in trans with a *Cctq1a^SJL^* allele. H) Homozygosity of the *Tyr^c-2J^* allele results in a significantly decreased stroma thickness compared to C57BL/6J mice. I) The *Tyr^c-2J^* allele in trans with a *Cctq1a^SJL^* allele results in a significantly decreased (fails to complement) stroma thickness compared to littermate controls having a *Tyr^c-2J^* allele in trans with a *Cctq1a^KS^* allele.

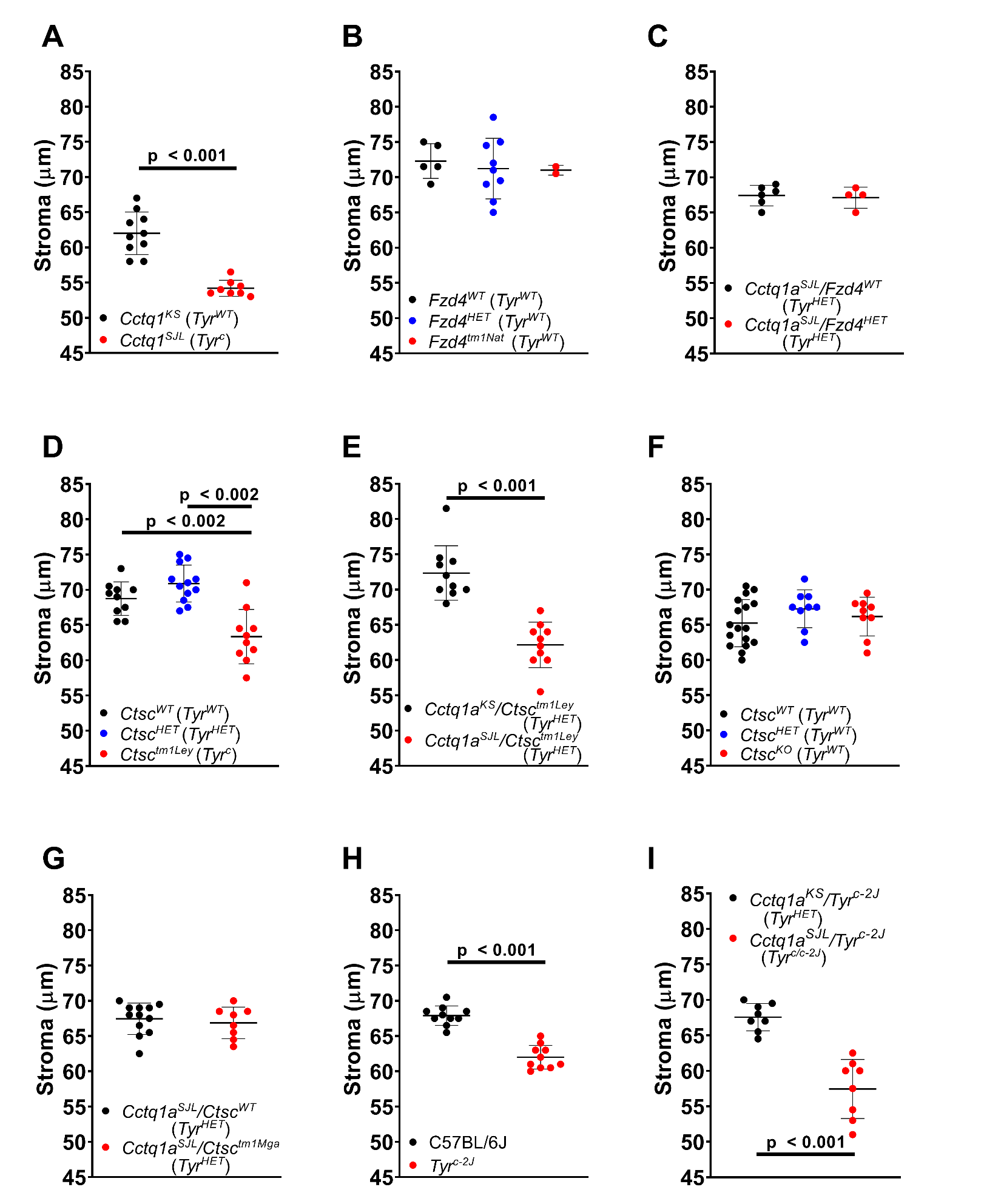
