## Supplementary material for "*Tyr* is Responsible for the *Cctq1a* QTL and Links Developmental Environment to Central Corneal Thickness Determination": Figure 3 -- Source Data 4

### CRISPR targeting strategy for *Ctsc*

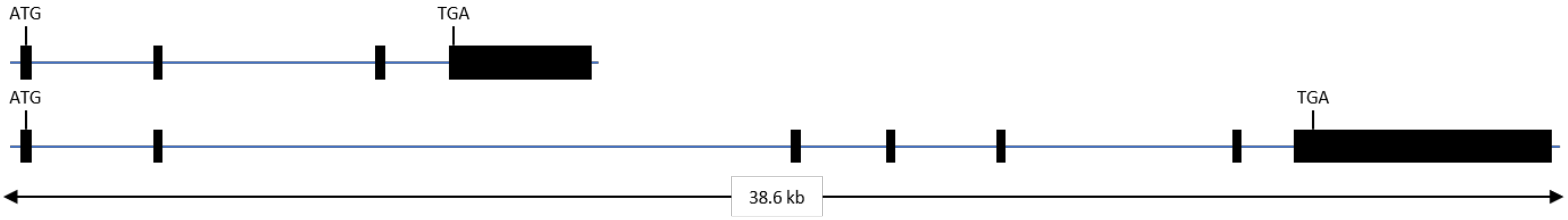


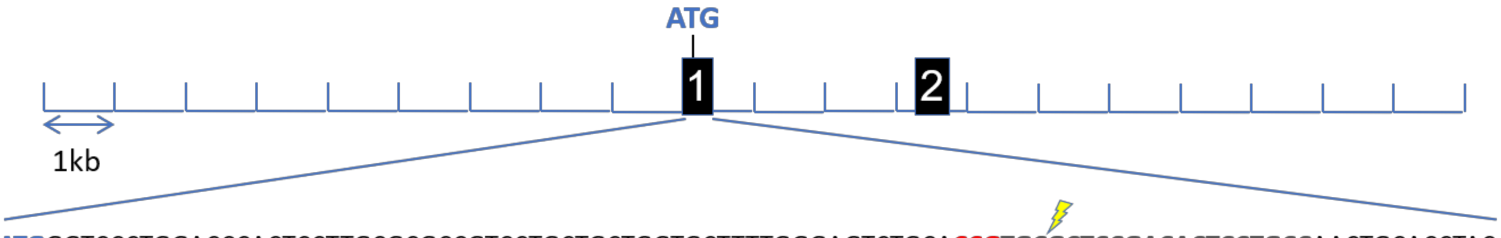


5’ **ATG**GGTCCCTGGACCCACTCCTTGCGCGCCGTCCTGCTGCTGGTGCTTTTGGGAGTCTGCA**CCG****TGCGCTCCGACACTCCTGCC**AACTGCACCTAC. . . 3’

**PAM** **Guide RNA**

↓ **Pronuclear Injection (C57BL/6J embryos)**
- synthetic crRNA:tracrRNA
- Cas9 Protein

↓ **Non-homologous end joining**

5’ **ATG**GGTCCCTGGACCCACTCCTTGCGCGCCGTCCTGCTGCTGGTGCTTTTGGGAGTCTGCA**CCG****TGC****{indel}****GCTCCGACACTCCTGCC**AACTGCACCTAC. . . 3’

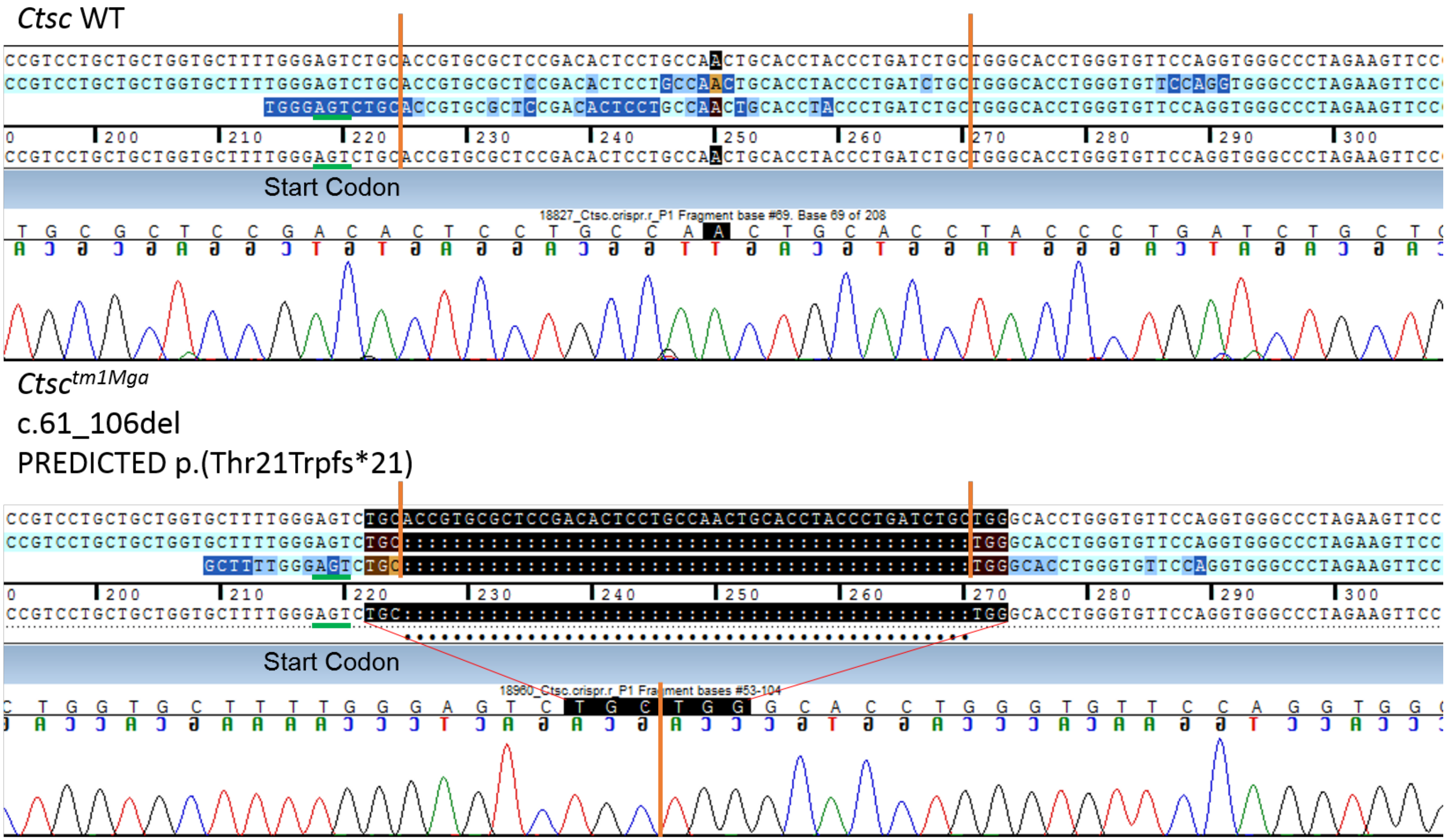
Ctsc WT
Start Codon
18827_Ctsc.crispr.r_P1 Fragment base #69. Base 69 of 208
Ctsc tm1Mga
c.61_106del
PREDICTED p.(Thr21Trpfs*21)
Start Codon
18960_Ctsc.crispr.r_P1 Fragment bases #53-104

*Ctsc* WT

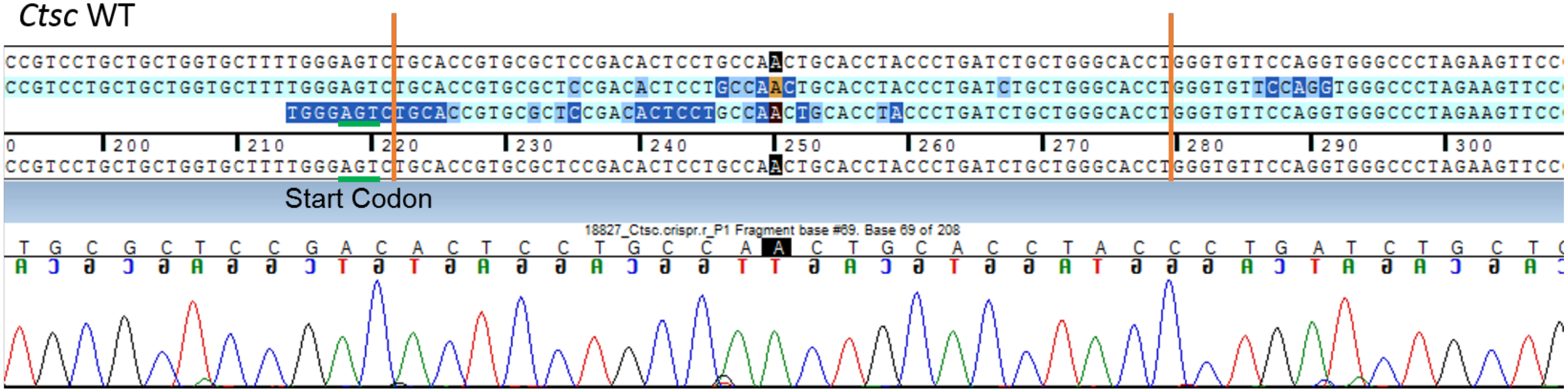


*Ctsc*$^{tm2Mga}$

c.[58_71del; 95_115del]

PREDICTED p.(Cys20Argfs*15)

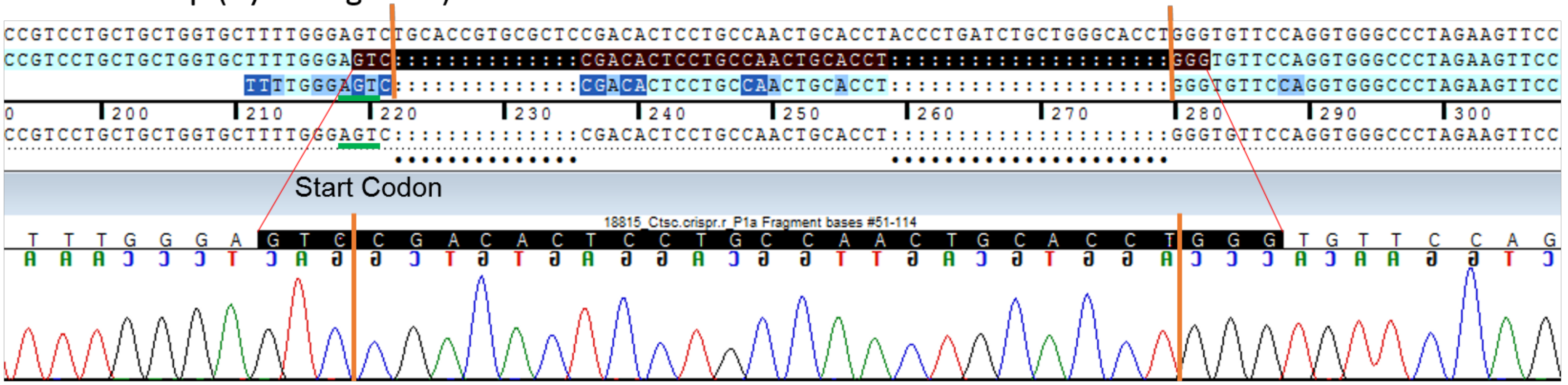

*Ctsc* WT

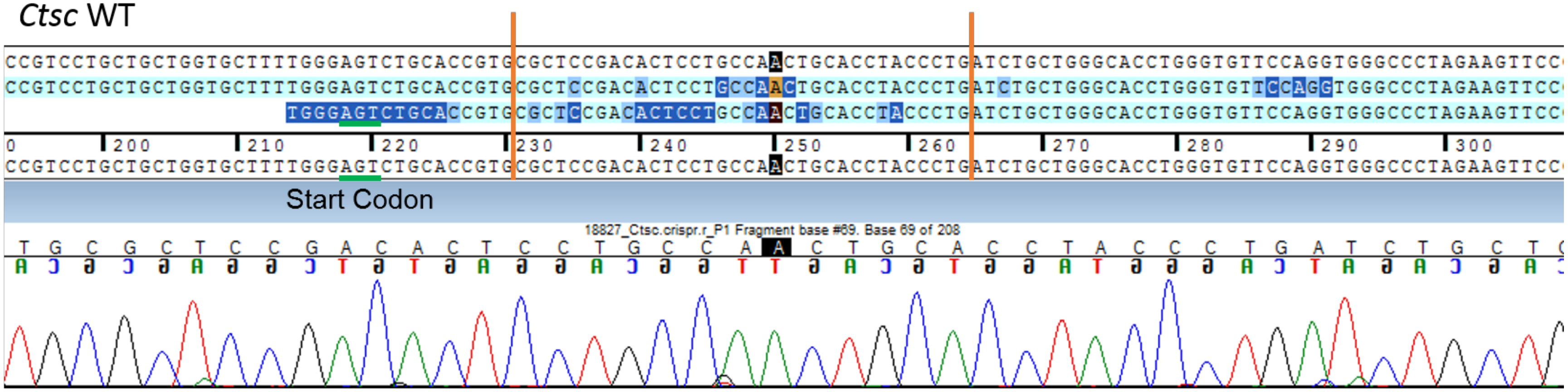


*Ctsc*$^{tm3Mga}$

c.[59G>A; 67_100del]

PREDICTED p.([Cys20Tyr; Arg23Ilefs*23])

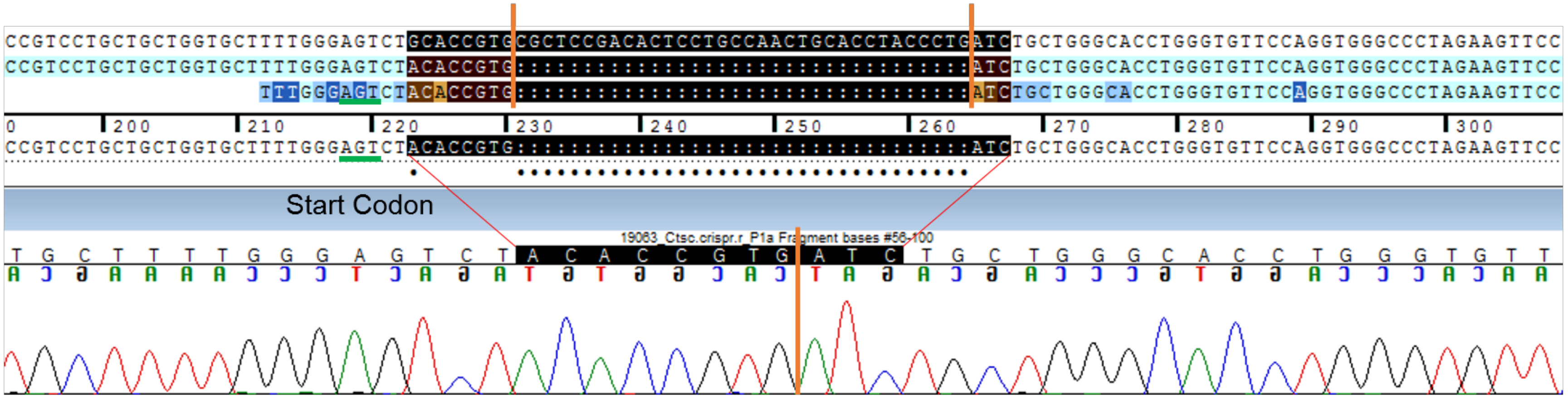

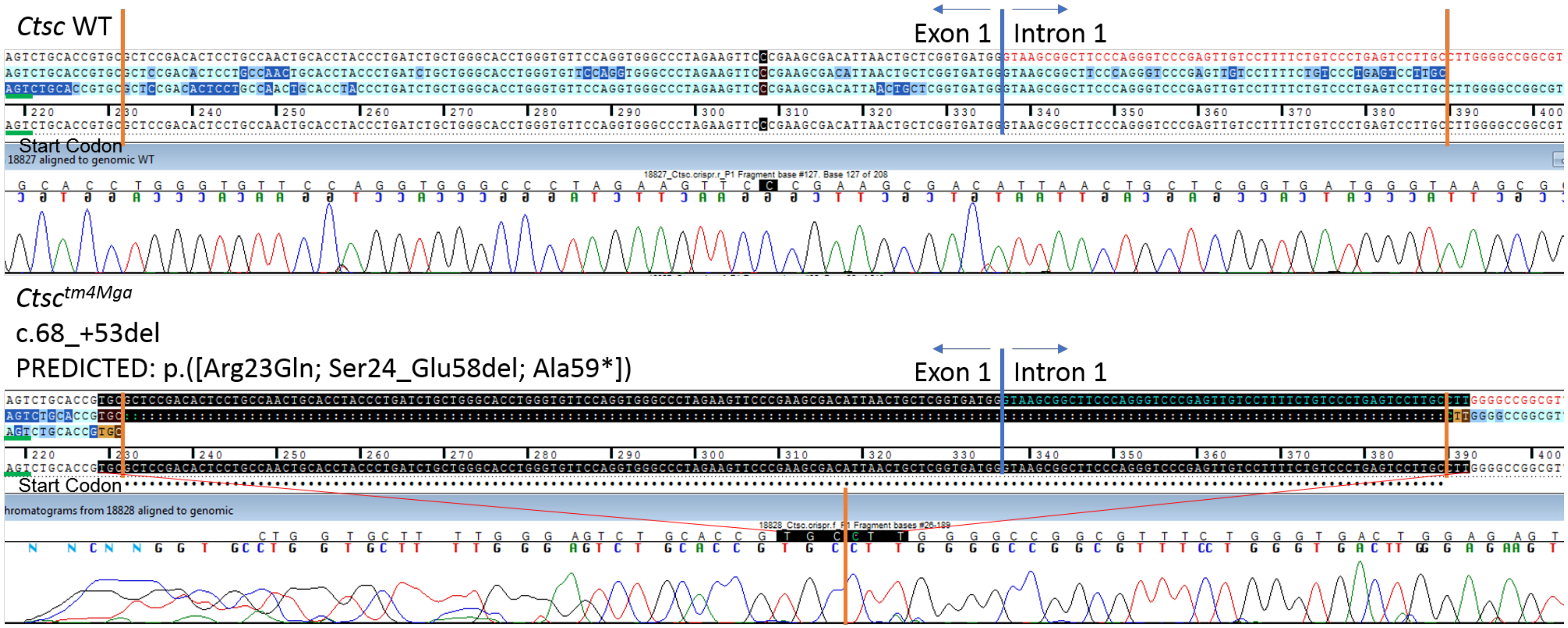
Ctsc WT
Exon 1
Intron 1
Start Codon
18827 aligned to genomic WT
Ctsc tm4Mga
c.68_+53del
PREDICTED: p.([Arg23Gln; Ser24_Glu58del; Ala59*])
Exon 1
Intron 1
Start Codon
hromatograms from 18828 aligned to genomic

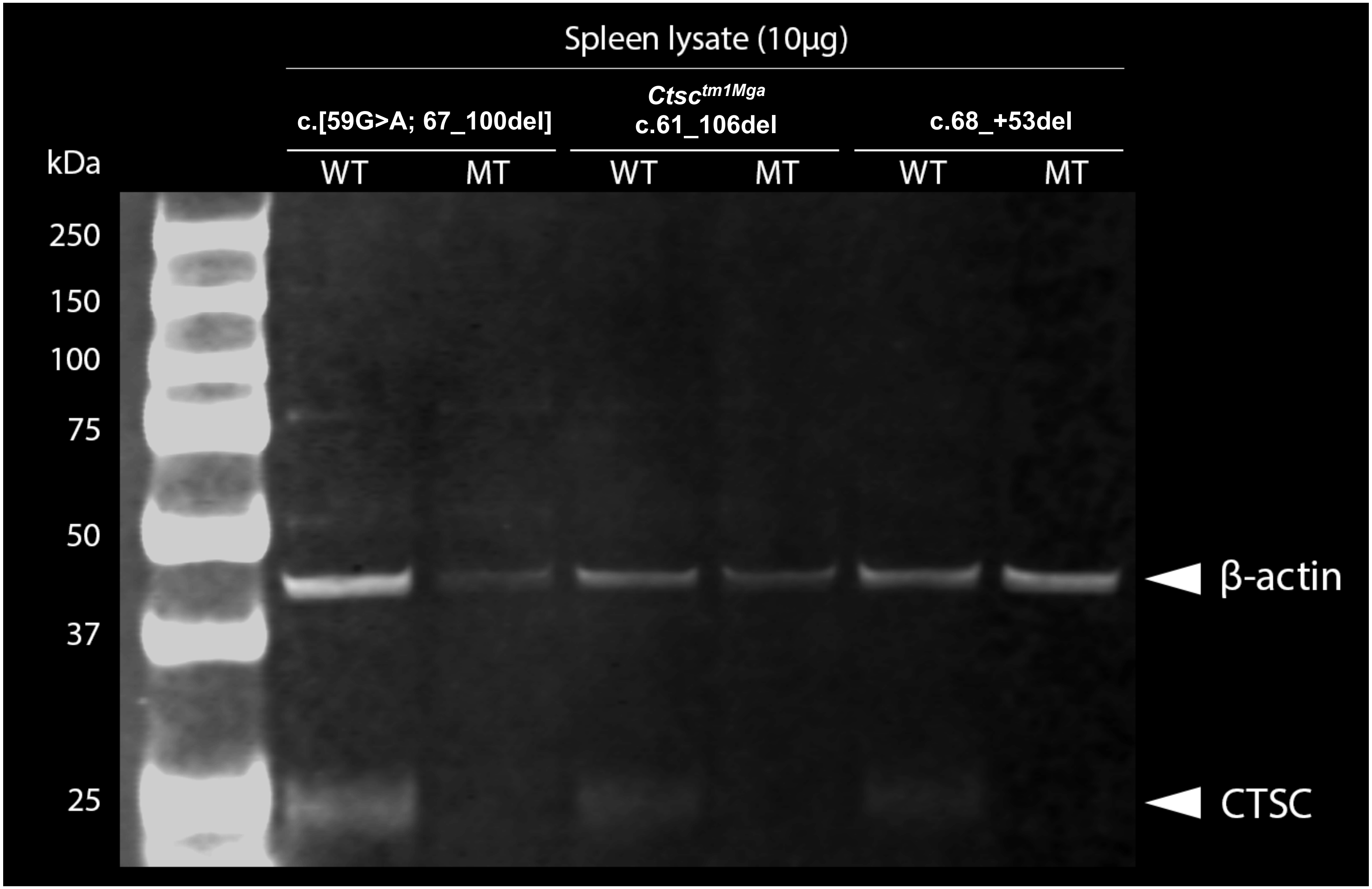
Spleen lysate (10µg)
c.[59G>A; 67_100del]
Ctsc tm1Mga
c.61_106del
c.68_+53del
WT
MT
WT
MT
WT
MT
kDa
250
150
100
75
50
37
25
β-actin
CTSC
