## Supplementary material for "*Tyr* is Responsible for the *Cctq1a* QTL and Links Developmental Environment to Central Corneal Thickness Determination": Figure 3 -- Source Data 5

Representative (**A, C**) photographs and (**B, D**) slit-lamp images of (**A, B**) *Fzd4^WT^*/*Cctq1a^SJL^* F1 mice and (**C, D**) *Fzd4^HET^*/*Cctq1a^SJL^* F1 mice. All mice pictured are heterozygous for the albinism-causing *Tyr^c^* allele.

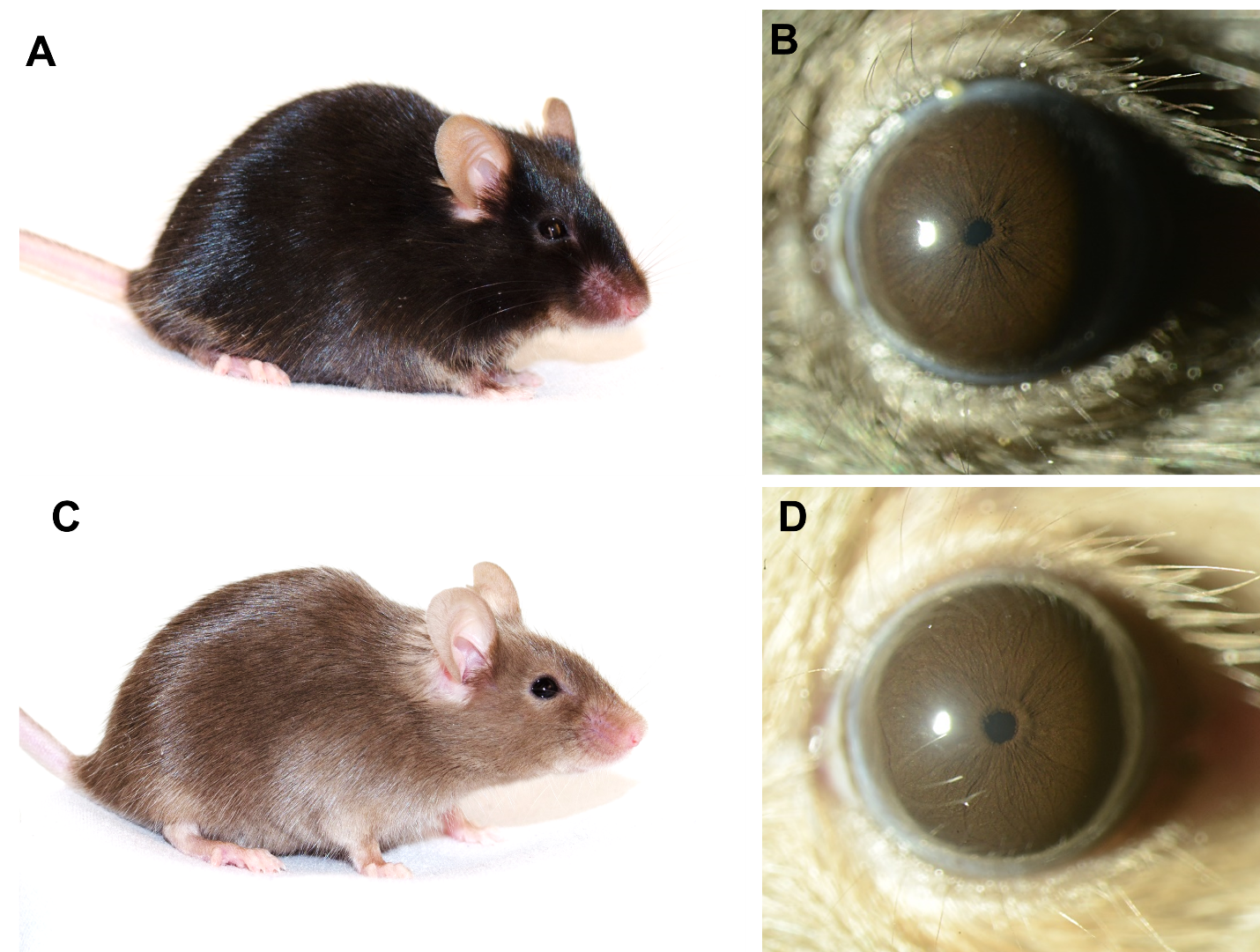
