## Supplementary material for "*Tyr* is Responsible for the *Cctq1a* QTL and Links Developmental Environment to Central Corneal Thickness Determination": Figure 4 -- Source Data 1

### CRISPR targeting strategy for *Tyr*

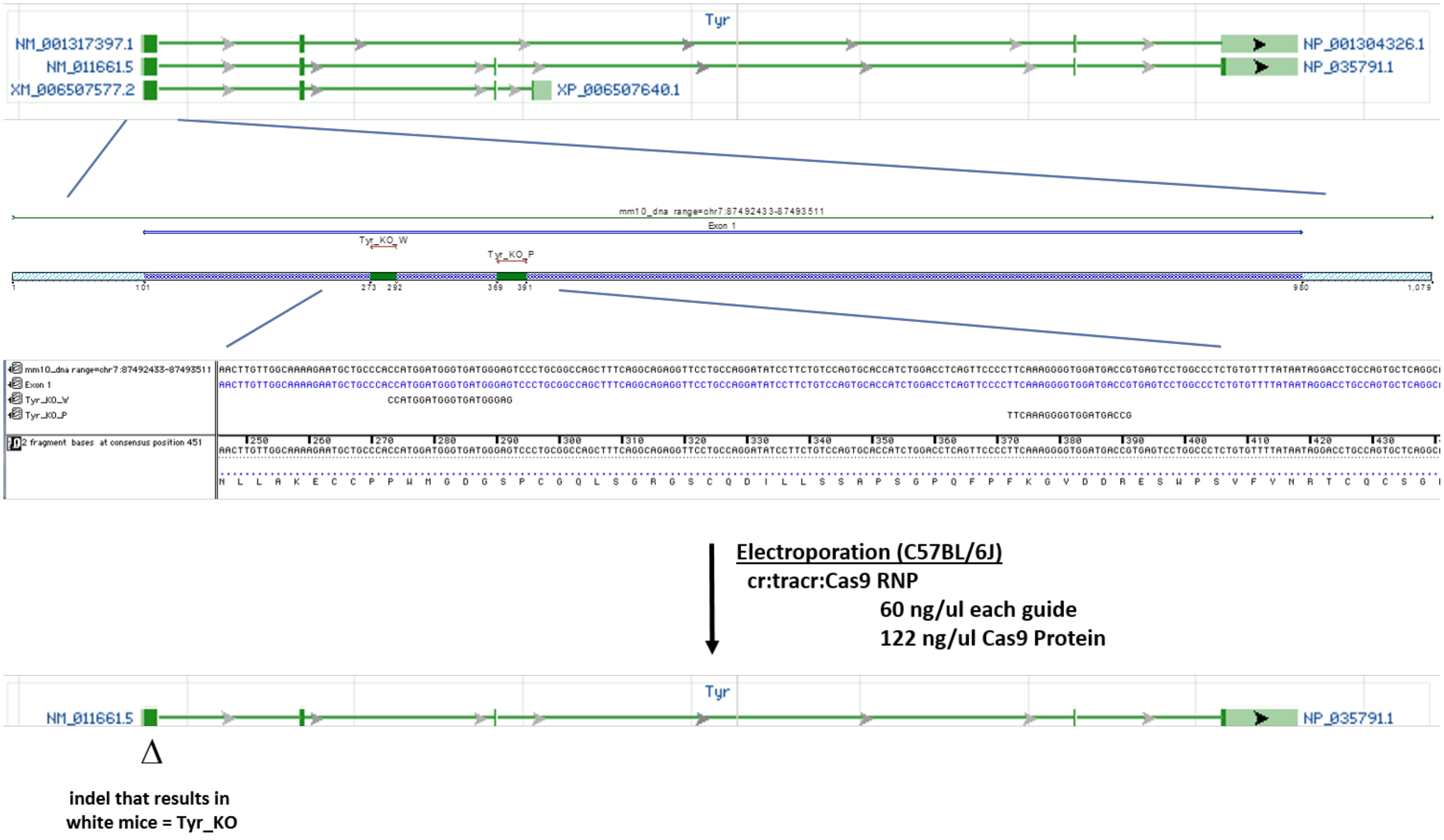

*Tyr* WT

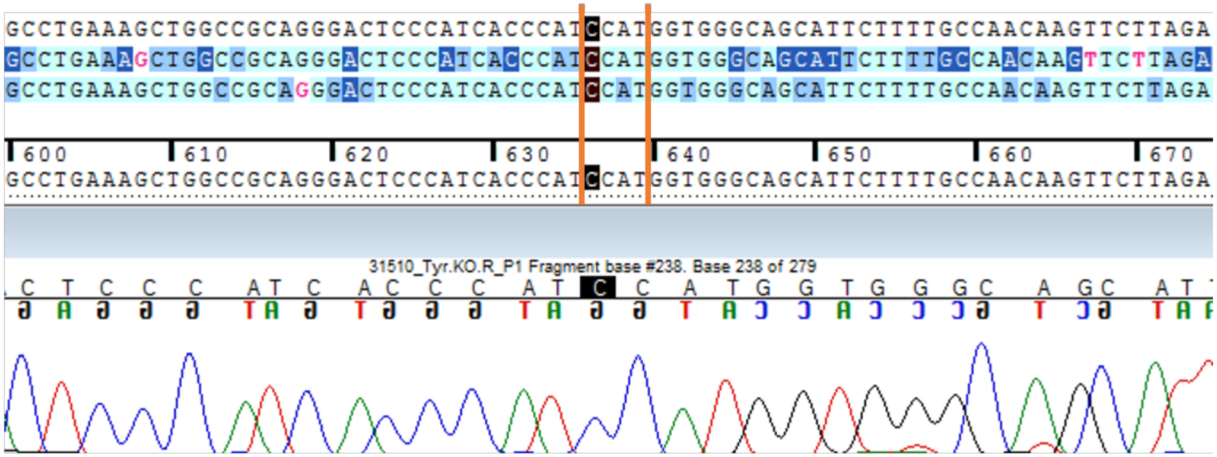


*Tyr*$^{tm4Mga}$

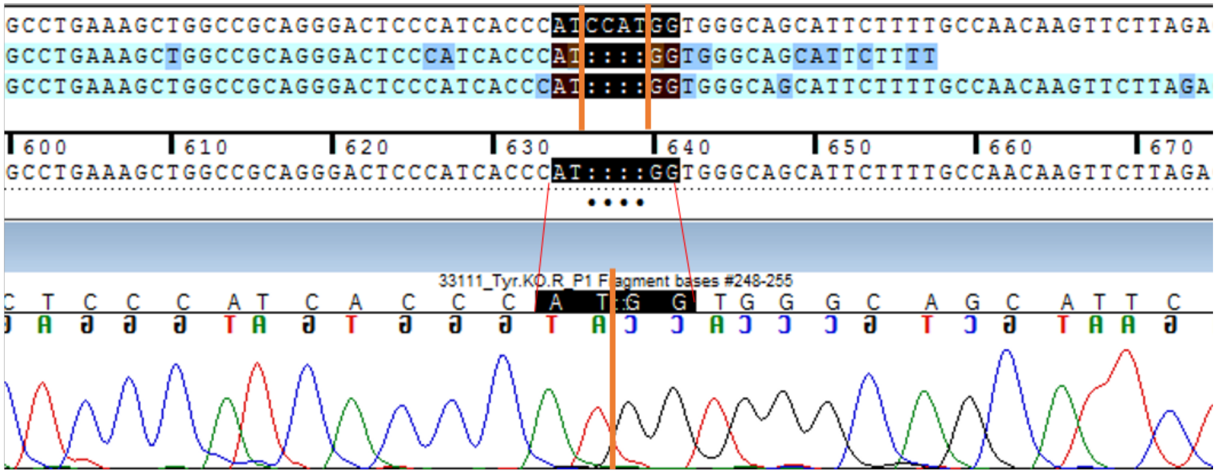


4bp-del

Fails to complement *Tyr*$^{c\text{-}2J}$ (albino pups)

*Tyr* WT

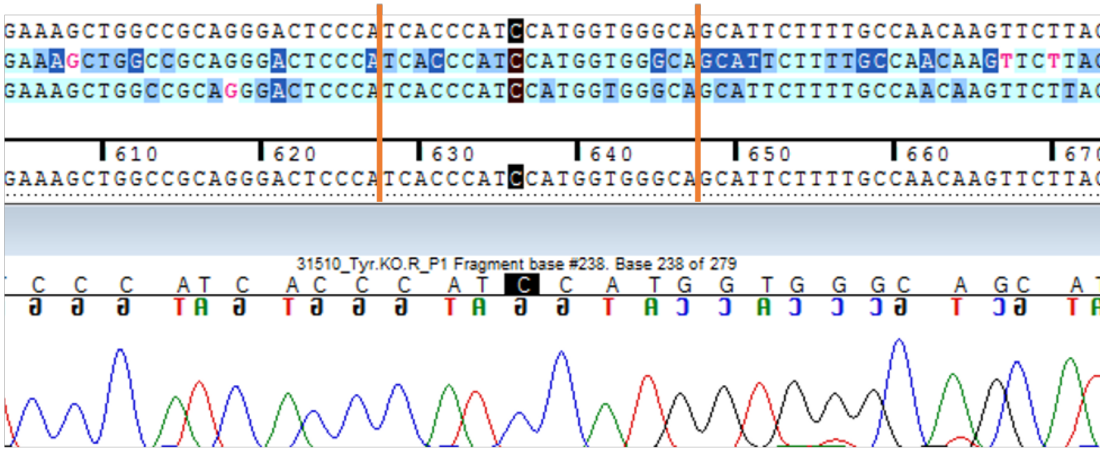


*Tyr*$^{tm5Mga}$

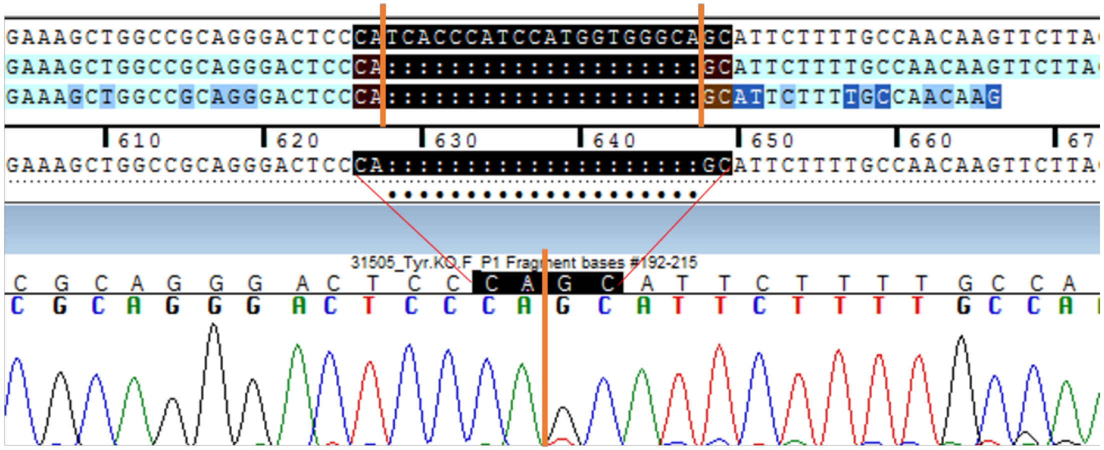


20bp-del

Fails to complement *Tyr*$^{c\text{-}2J}$ (albino pups)

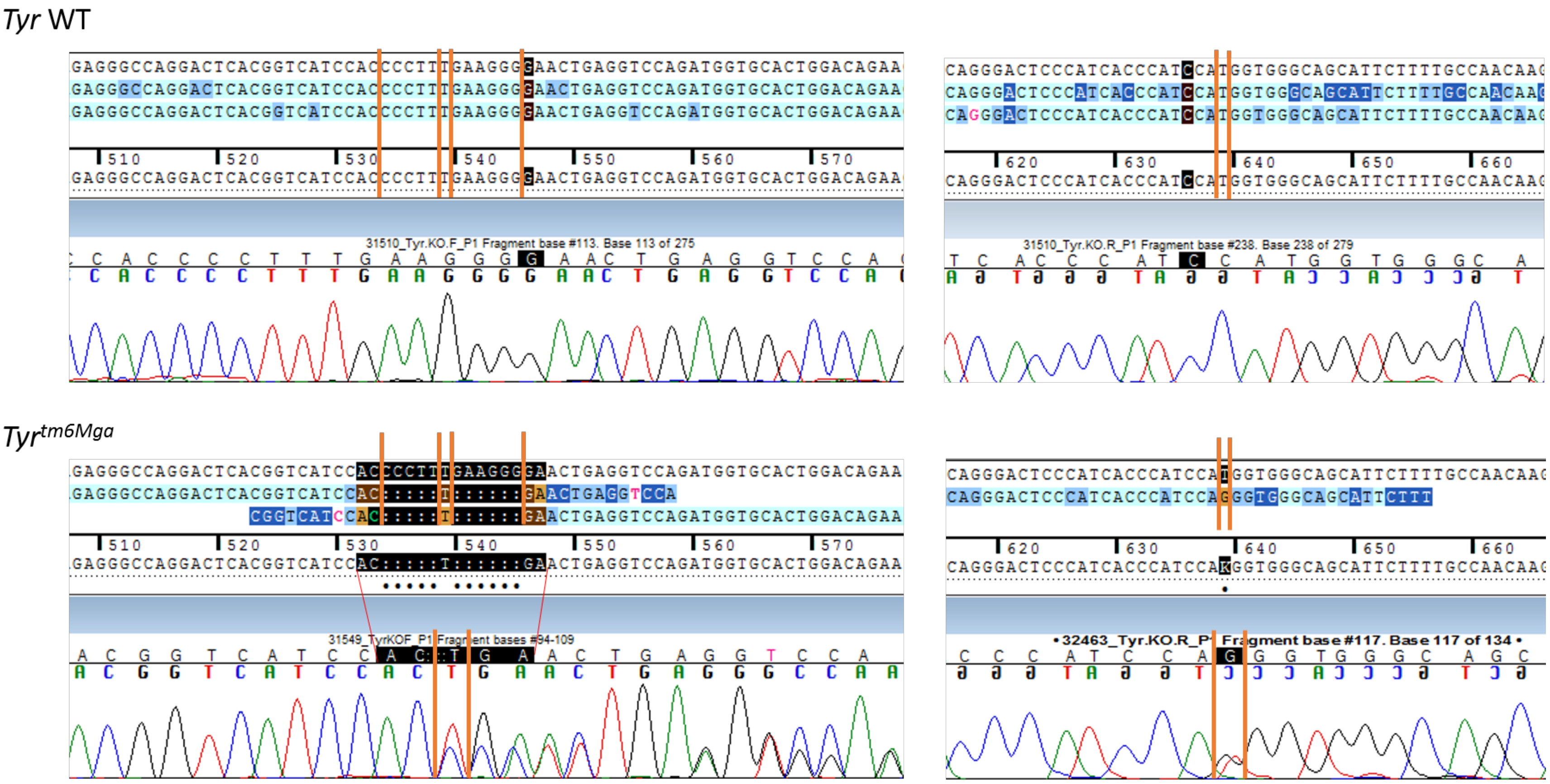


5bp-del and 6bp-del and 1bp T>G

Fails to complement *Tyr*$^{c-2J}$ (albino pups)

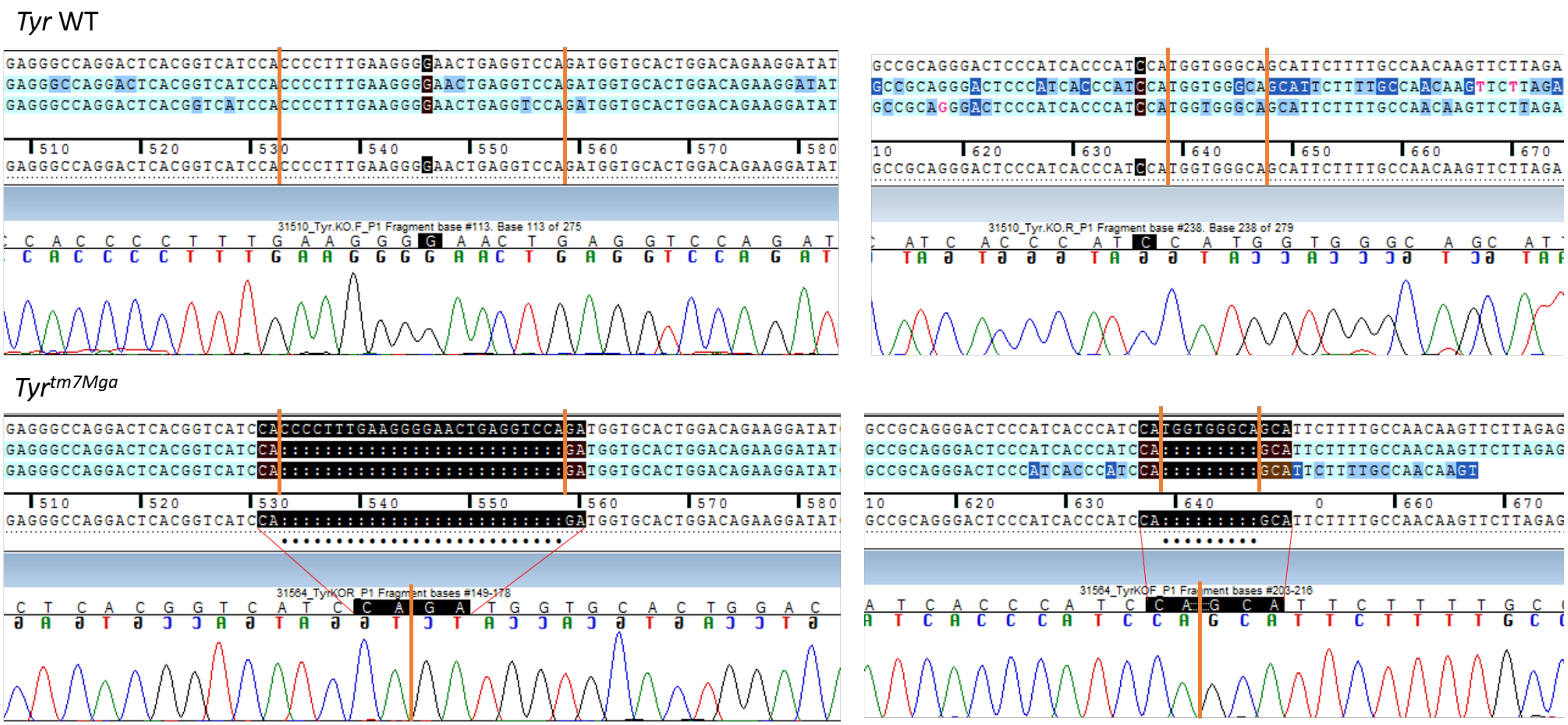


26bp-del and 9bp-del

Fails to complement *Tyr*$^{c\text{-}2J}$ (albino pups)

*Tyr* WT

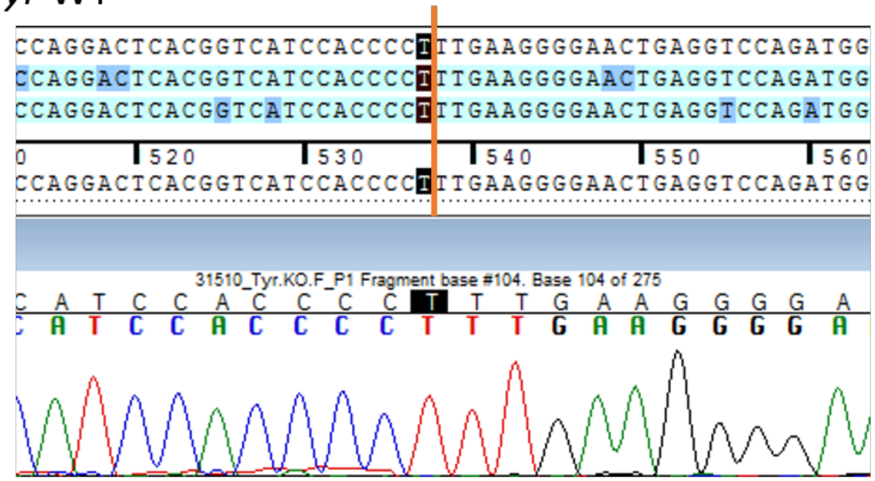


*Tyr*$^{tm8Mga}$

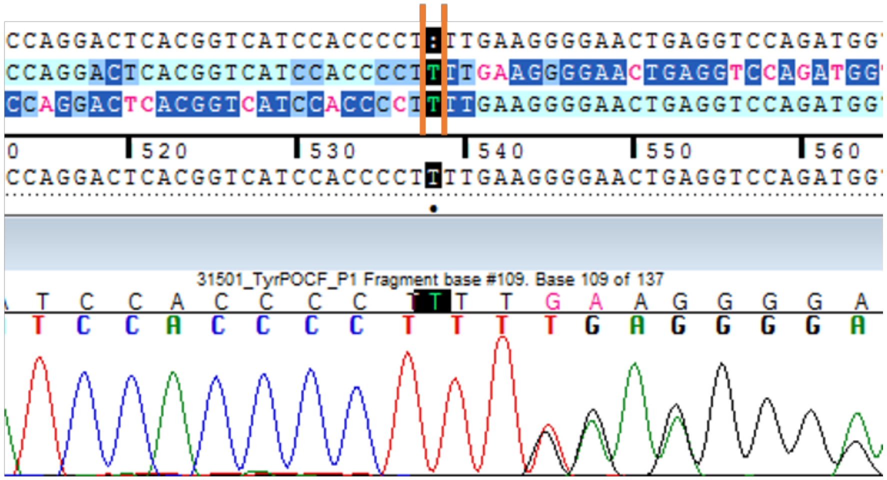


1bp-ins T

Fails to complement *Tyr*$^{c\text{-}2J}$ (albino pups)

*Tyr* WT

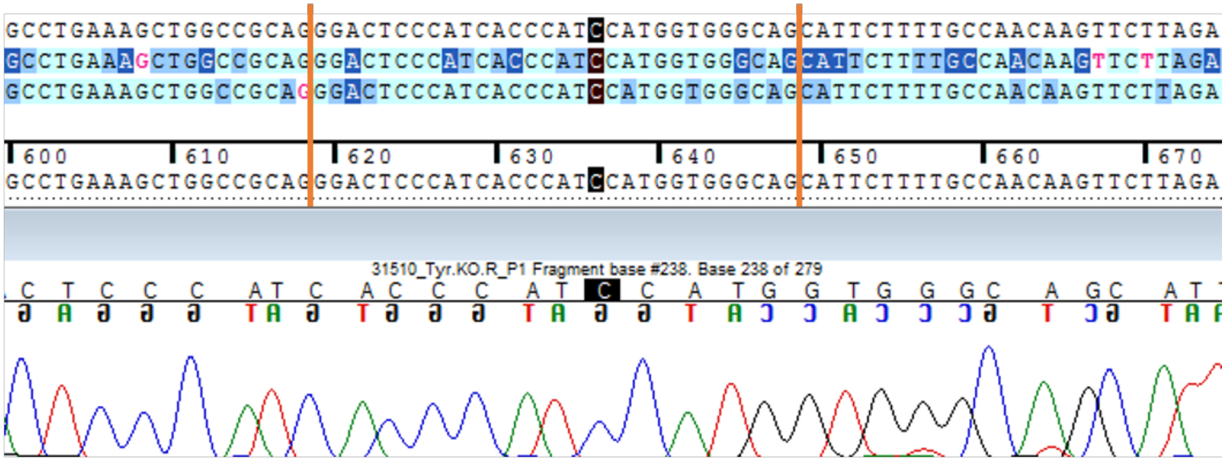


*Tyr*$^{tm9Mga}$

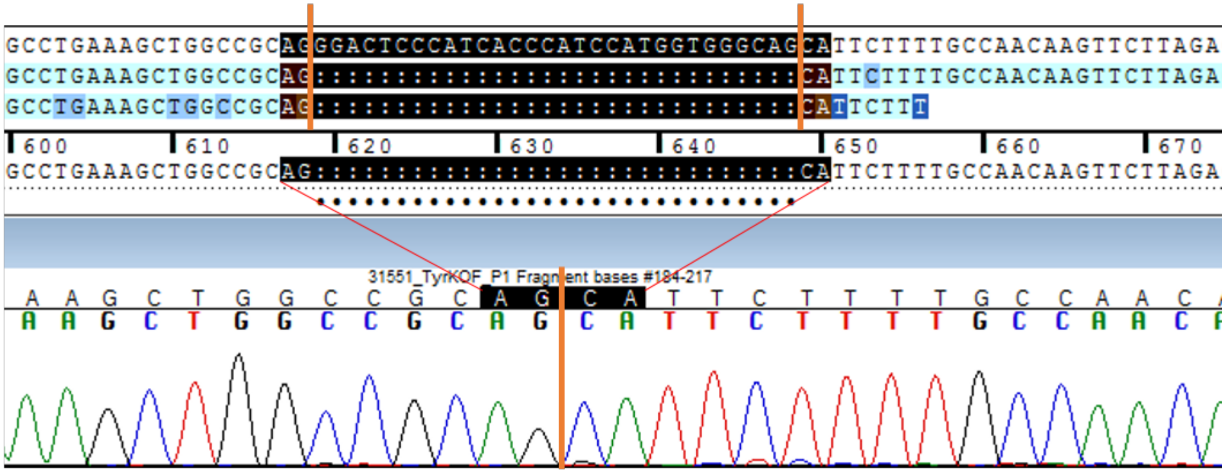


30bp-del

Fails to complement *Tyr*$^{c\text{-}2J}$ (albino pups)

*Tyr* WT

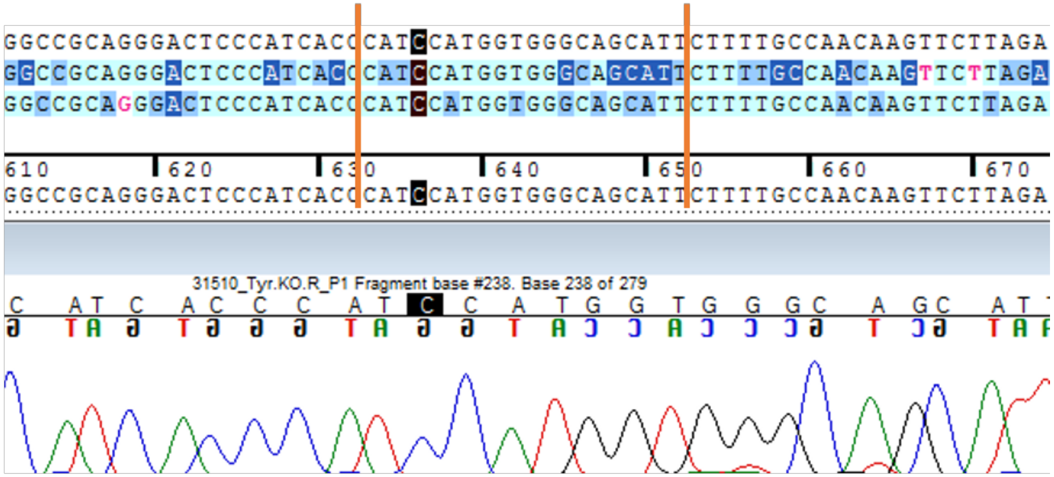


*Tyr*$^{tm10Mga}$

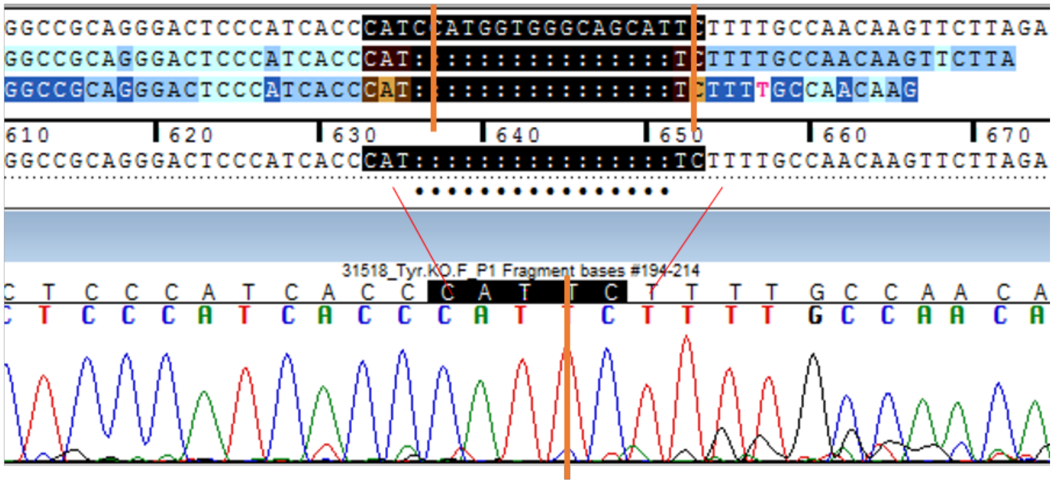


16bp-del

Fails to complement *Tyr*$^{c\text{-}2J}$ (albino pups)

*Tyr* WT

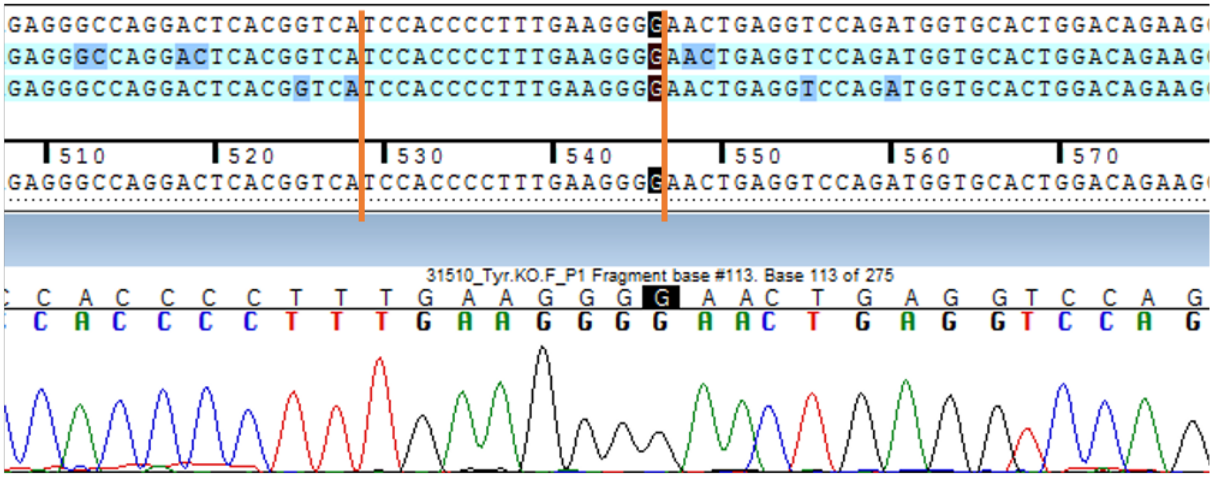


*Tyr*$^{tm11Mga}$

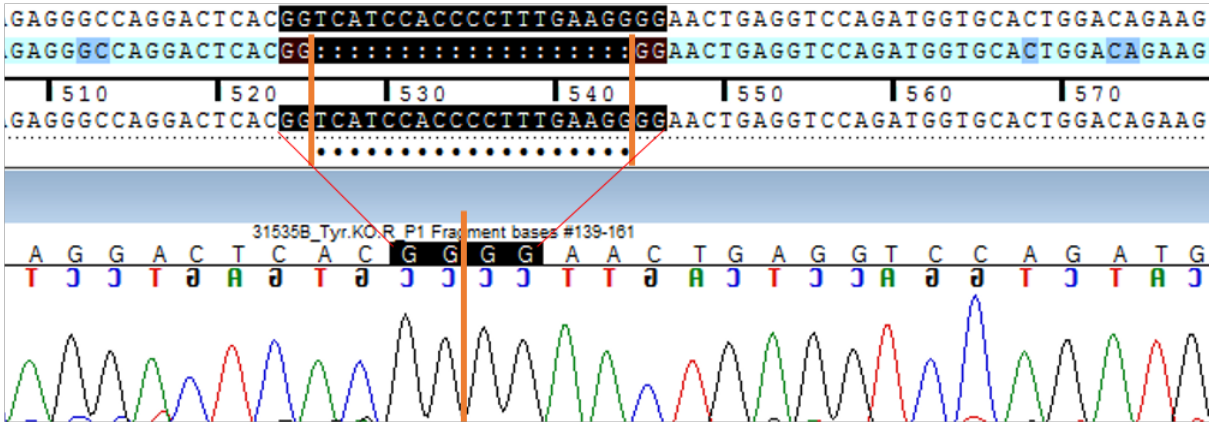


19bp-del

Fails to complement *Tyr*$^{c-2J}$ (albino pups)

*Tyr* WT

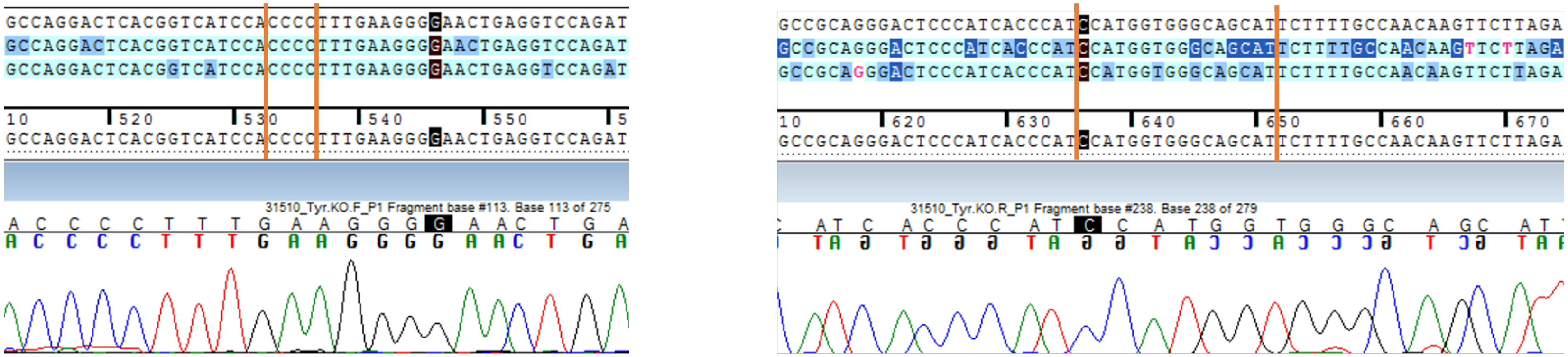


*Tyr*$^{tm12Mga}$

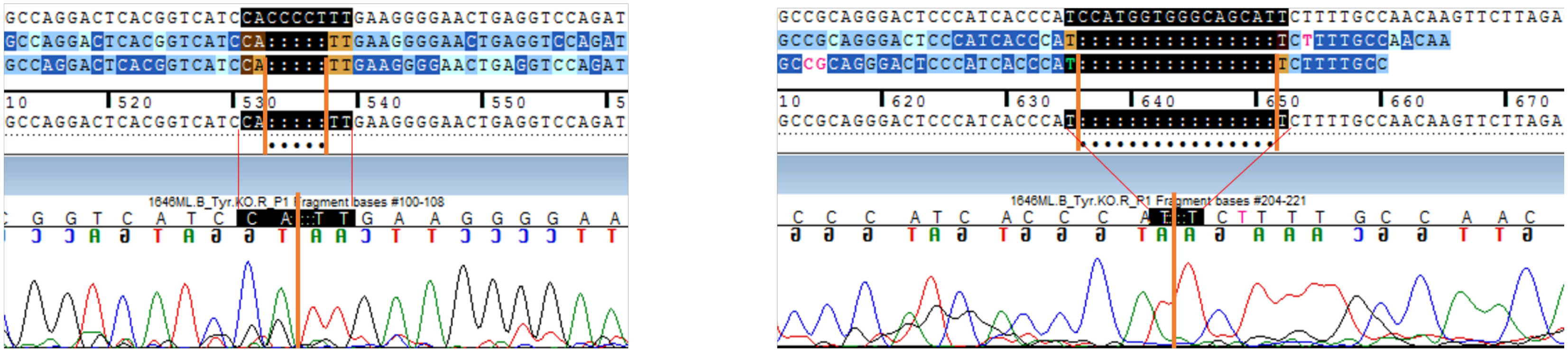


5bp-del and 16bp-del and ??

Did not produce pups in complementation cross to *Tyr*$^{c\text{-}2J}$

*Tyr* WT

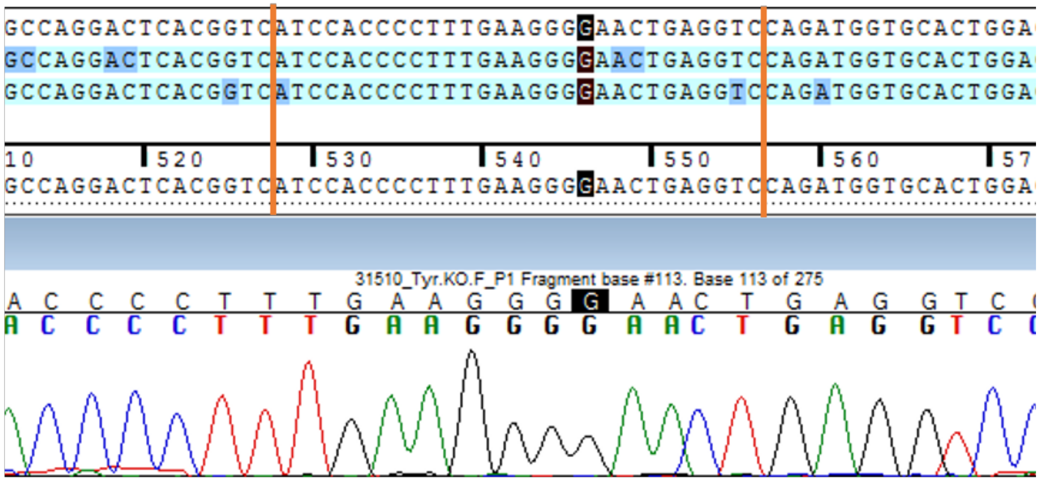


*Tyr*$^{tm13Mga}$

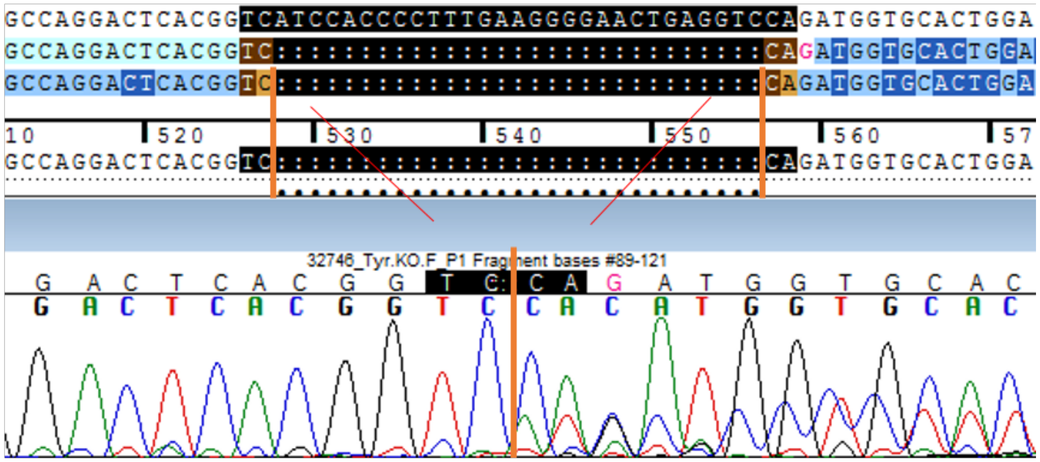


29bp-del

Fails to complement *Tyr*$^{c\text{-}2J}$ (albino pups)

*Tyr* WT

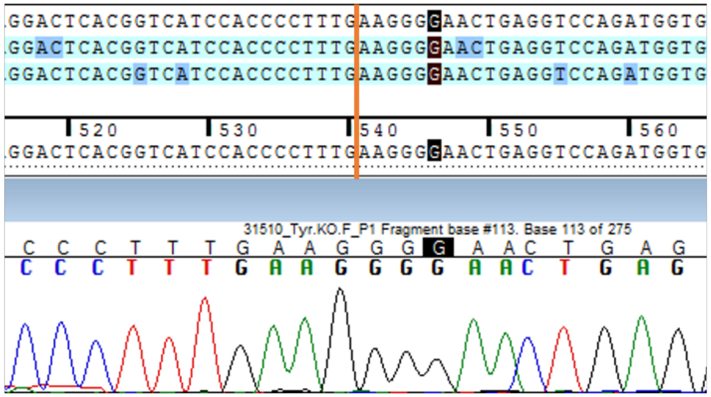


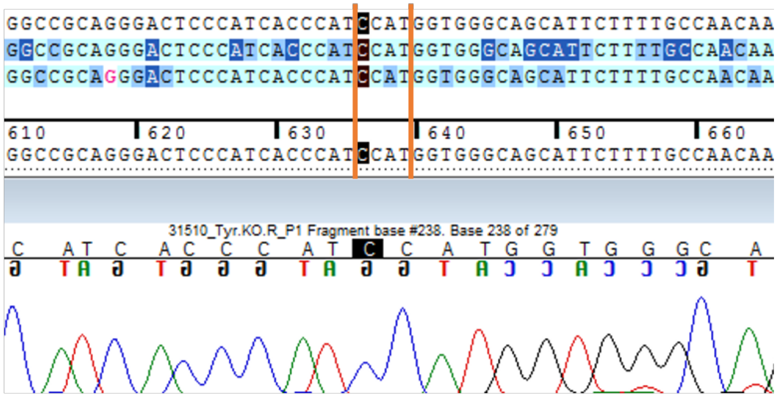


*Tyr^tm14Mga*

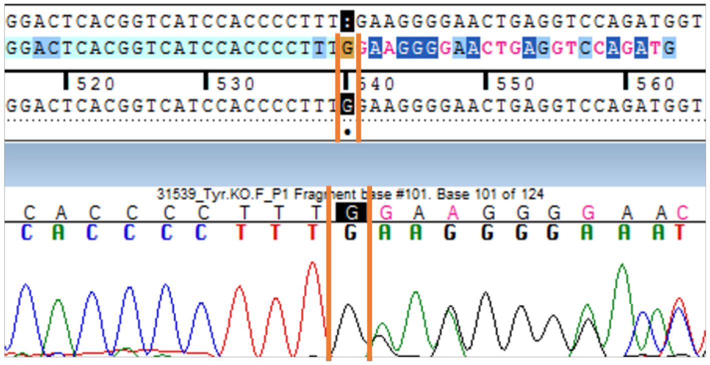


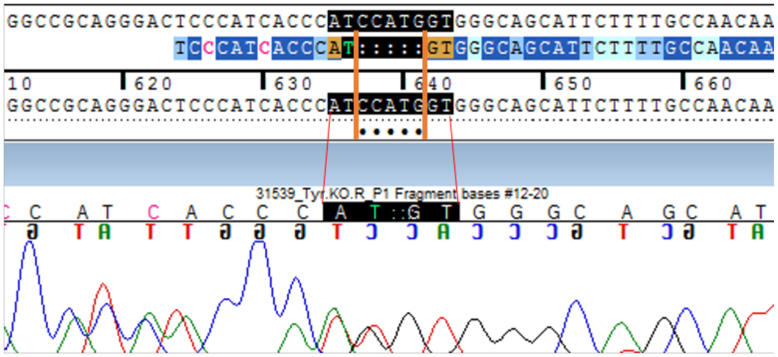


1bp-ins G and 5bp-del and ??

Fails to complement *Tyr^c-2J* (albino pups)

*Tyr* WT

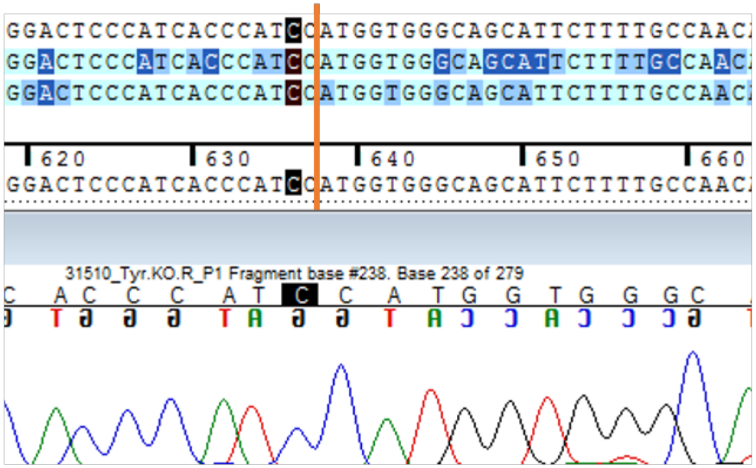


*Tyr*$^{tm15Mga}$

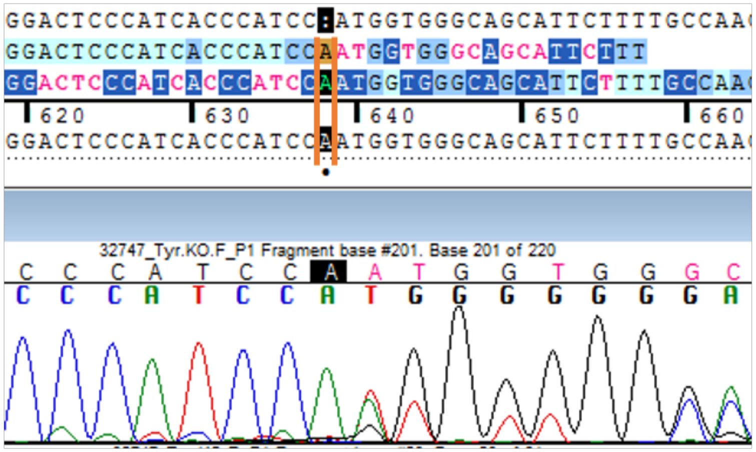


1bp-ins A

Fails to complement *Tyr*$^{c\text{-}2J}$ (albino pups)

*Tyr* WT

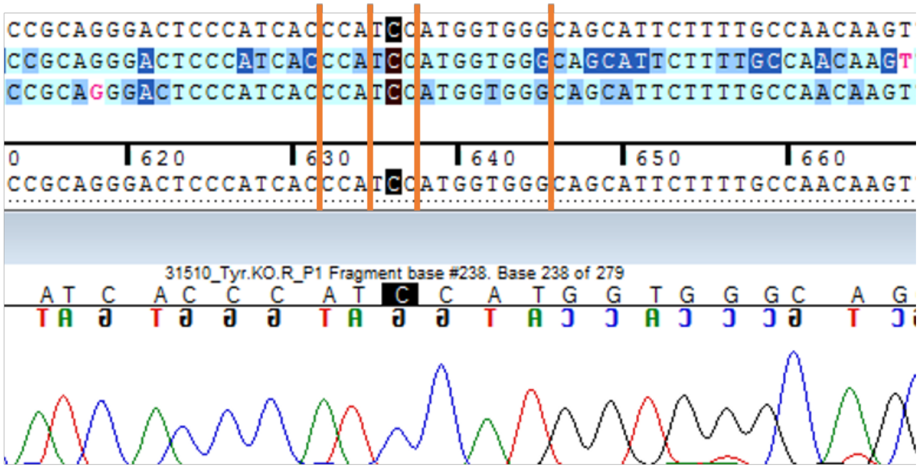


*Tyr*$^{tm16Mga}$

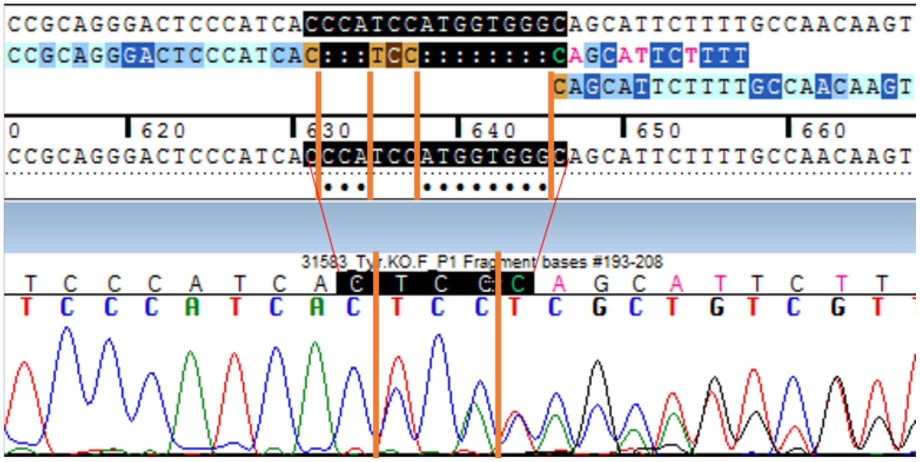


3bp-del and 8bp-del

Fails to complement *Tyr*$^{c\text{-}2J}$ (albino pups)

*Tyr* WT

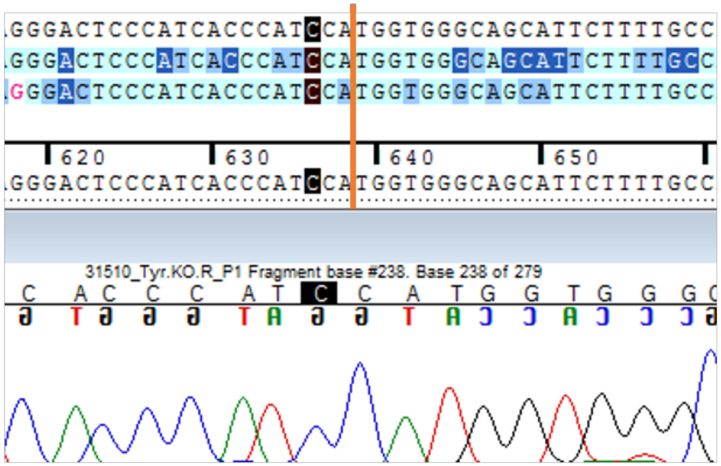


*Tyr*$^{tm17Mga}$

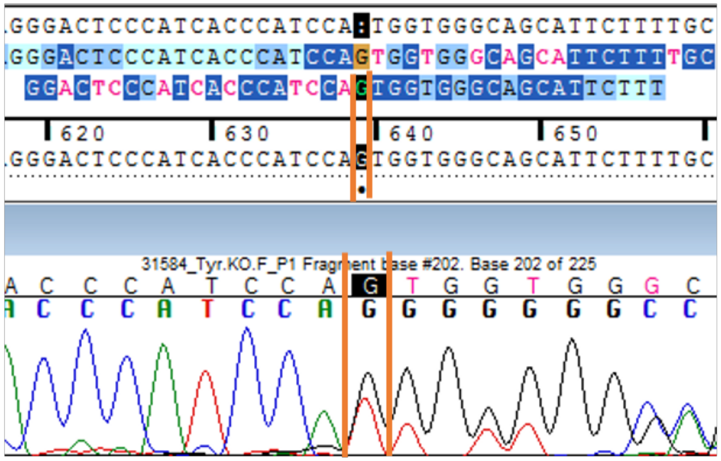


1bp-ins G

Fails to complement *Tyr*$^{c\text{-}2J}$ (albino pups)

*Tyr* WT

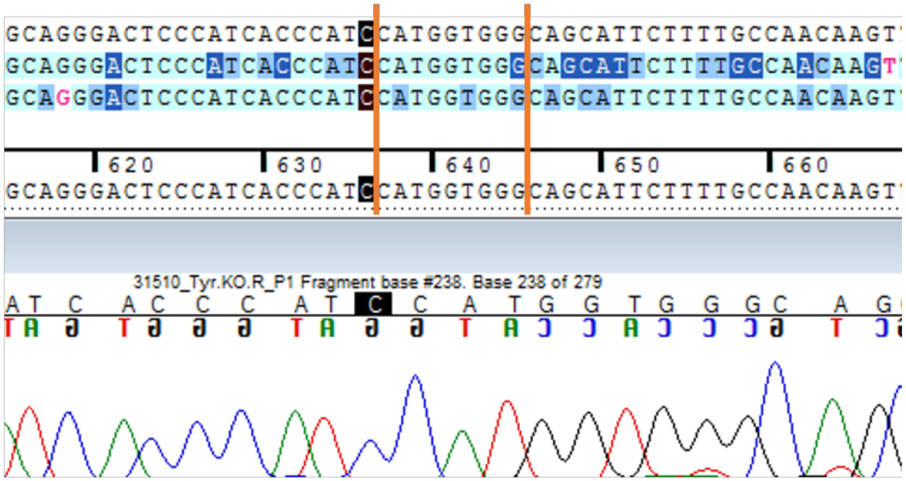


*Tyr*$^{tm18Mga}$

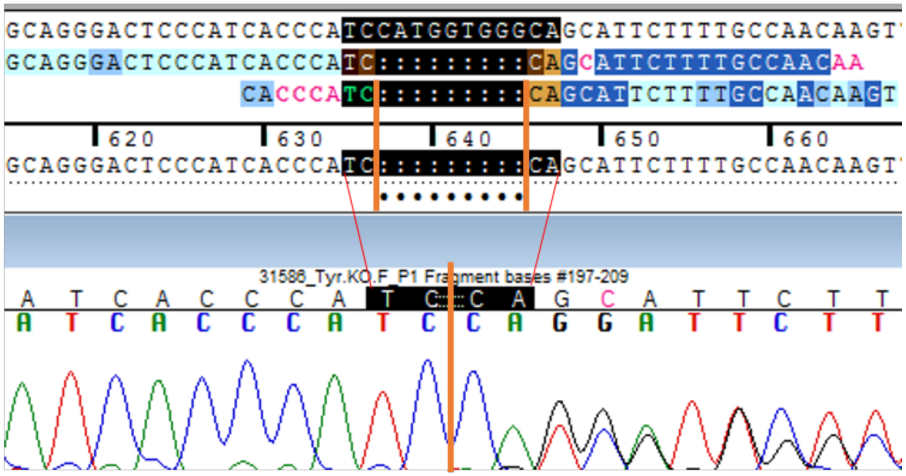


9bp-del

Fails to complement *Tyr*$^{c\text{-}2J}$ (albino pups)
