## Supplementary material for "*Tyr* is Responsible for the *Cctq1a* QTL and Links Developmental Environment to Central Corneal Thickness Determination": Figure 4 -- Source Data 3

The *Tyr^tm4Mga^* allele has no effect on epithelium thickness, in neither the heterozygous nor homozygous state, compared to *Tyr^WT^* littermate controls (*left*). Homozygosity of the *Tyr^tm4Mga^* allele results in a significantly decreased stroma thickness compared to littermate controls (*right*).

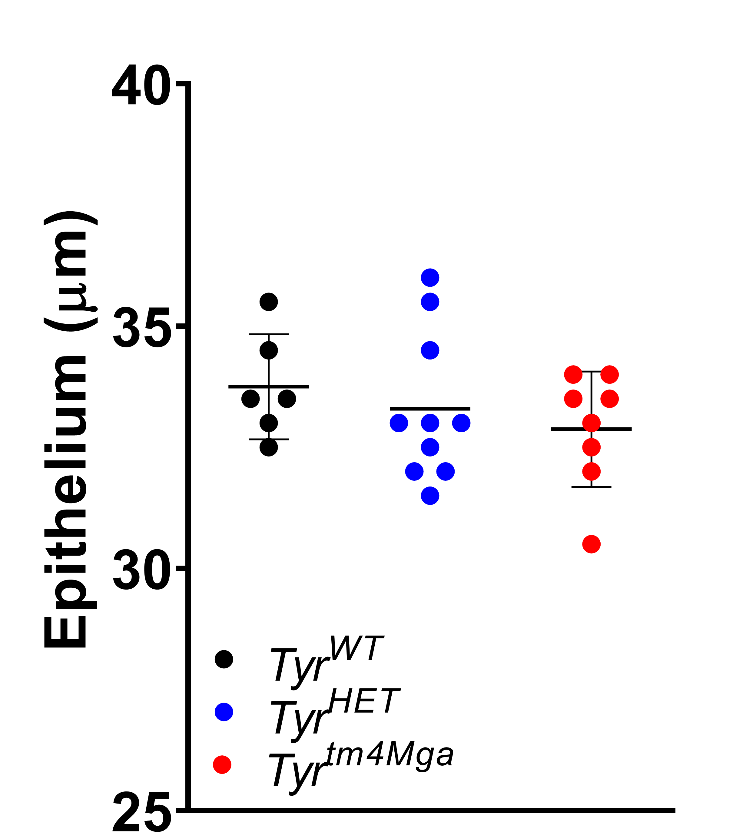

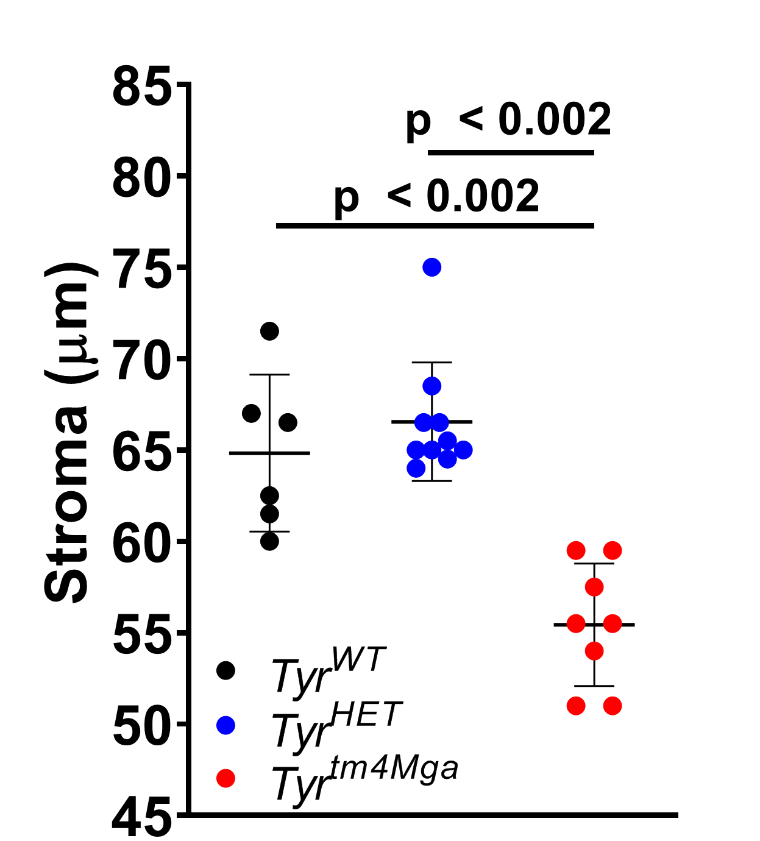
